# RosettaAntibodyDesign (RAbD): A General Framework for Computational Antibody Design

**DOI:** 10.1101/183350

**Authors:** Jared Adolf-Bryfogle, Oleks Kalyuzhniy, Michael Kubitz, Brian D. Weitzner, Xiaozhen Hu, Yumiko Adachi, William R. Schief, Roland L. Dunbrack

## Abstract

A structural-bioinformatics-based computational methodology and framework have been developed for the design of antibodies to targets of interest. RosettaAntibodyDesign (RAbD) samples the diverse sequence, structure, and binding space of an antibody to an antigen in highly customizable protocols for the design of antibodies in a broad range of applications. The program samples antibody sequences and structures by grafting structures from a widely accepted set of the canonical clusters of CDRs (North et al., *J. Mol. Biol*., 406:228-256, 2011). It then performs sequence design according to amino acid sequence profiles of each cluster, and samples CDR backbones using a flexible-backbone design protocol incorporating cluster-based CDR constraints. Starting from an existing experimental or computationally modeled antigen-antibody structure, RAbD can be used to redesign a single CDR or multiple CDRs with loops of different length, conformation, and sequence. We rigorously benchmarked RAbD on a set of 60 diverse antibody–antigen complexes, using two design strategies – optimizing total Rosetta energy and optimizing interface energy alone. We utilized two novel metrics for measuring success in computational protein design. The design risk ratio (DRR) is equal to the frequency of recovery of native CDR lengths and clusters divided by the frequency of sampling of those features during the Monte Carlo design procedure. Ratios greater than 1.0 indicate that the design process is picking out the native more frequently than expected from their sampled rate. We achieved DRRs for the non-H3 CDRs of between 2.4 and 4.0. The antigen risk ratio (ARR) is the ratio of frequencies of the native amino acid types, CDR lengths, and clusters in the output decoys for simulations performed in the presence and absence of the antigen. For CDRs, we achieved cluster ARRs as high as 2.5 for L1 and 1.5 for H2. For sequence design simulations without CDR grafting, the overall recovery for the native amino acid types for residues that contact the antigen in the native structures was 72% in simulations performed in the presence of the antigen and 48% in simulations performed without the antigen, for an ARR of 1.5. For the non-contacting residues, the ARR was 1.08. This shows that the sequence profiles are able to maintain the amino acid types of these conserved, buried sites, while recovery of the exposed, contacting residues requires the presence of the antigen-antibody interface. We tested RAbD experimentally on both a lambda and kappa antibody–antigen complex, successfully improving their affinities 10 to 50 fold by replacing individual CDRs of the native antibody with new CDR lengths and clusters.

**Author Summary:** Antibodies are proteins produced by the immune system to attack infections and cancer and are also used as drugs to treat cancer and autoimmune diseases. The mechanism that has evolved to produce them is able to make 10s of millions of different antibodies, each with a different surface used to bind the foreign or mutated molecule. We have developed a method to design antibodies computationally, based on the 1000s of experimentally determined three-dimensional structures of antibodies available. The method works by treating pieces of these structures as a collection of parts that can be combined in new ways to make better antibodies. Our method has been implemented in the protein modeling program Rosetta, and is called RosettaAntibodyDesign (RAbD). We tested RAbD both computationally and experimentally. The experimental test shows that we can improve existing antibodies by 10 to 50 fold, paving the way for design of entirely new antibodies in the future.

## Introduction

Antibodies are a key component of the adaptive immune system and form the basis of its ability to detect and respond to foreign pathogens through binding of molecular epitopes. Antibodies are increasingly a focus of biomedical research for drug and vaccine development in addition to their numerous applications in biotechnology by private companies, government, and academia [1–7]. Experimentally, antibodies may be discovered and optimized through in vitro phage and yeast display [8,9], screening with large antibody libraries [10–13] and/or affinity maturation through error-prone PCR [14–16]. They may also be derived in vivo through a combination of animal immunization and antibody screening through ELISA or Western blots, and humanization of the animal antibody [17–19].

Although these methods have been successfully applied to create new antibodies, they can take many months to complete and can be prohibitively expensive. In addition, for many targets, these methods may not produce antibodies with desirable properties, because the antigen is difficult to target [20–22] or because the antibody is required to bind to a specific epitope for various functional reasons such as the neutralization of a target pathogen [23], initialization of a downstream signaling cascade [24], or the blocking of a binding protein from being able to engage the site [25]. We believe that computational design methods developed specifically for antibodies can be used in tandem with state-of-the-art experimental methods to save time, money, and increase our ability to design or enhance antibodies to many different targets.

Various computational methods including rational, structure-based design, protein design algorithms, and antibody-specific modeling techniques can aid in the design of antibodies [26–28]. General protein design methods have been applied to affinity maturation [29–31], improving stability [32–34], humanization [35,36], and the design of phage/yeast display libraries [37–40], while three software programs have been developed specifically for antibody computational design. Maranas and colleagues have developed the OptCDR [41] and OptMAVEn [42] methods, which sample and combine elements of antibody structure in an effort to assemble antibodies to bind to novel epitopes. OptCDR samples from clusters of the six CDRs in the presence of a fixed antigen position. This is followed by placement of side chains according to sequence preferences within each cluster, a rotamer search from a backbone-dependent rotamer library [43], and a CHARMM-based energy function. The method has not been experimentally tested. OptMAVEn divides antibody structures in a manner inspired by V(D)J recombination: antibody heavy- and light-chain V regions, CDR3s, and post-CDR3 segments from the MAPS database [44]. OptMAVEn has been tested experimentally and was used to design antibodies against a very hydrophobic heptamer peptide antigen with a repetitive sequence (FYPYPYA), starting from the structure of an existing antibody bound to a dodecamer peptide containing this sequence (PDB 4HOH [45]; only the heptamer has coordinates and was used in the design process) [46].

Lapidoth et al. have presented AbDesign [47] that follows a similar methodology to OptMAVEn, breaking up antibodies into V regions and CDR3 by analogy to V(D)J recombination. They clustered V region structures purely by length of the CDR1 and CDR2 segments in VH and VL, grouping sequences from distantly related germlines and different CDR conformations into clusters from which sequence profiles were derived. As implemented in the Rosetta Software Suite, AbDesign combinatorially builds antibodies and performs sequence design from position-specific scoring matrices of aligned antibody sequences of their length-based clusters of the V regions and CDR3 regions. Because CDR1 and CDR2 are grafted together, AbDesign has limited flexibility in terms of setting which CDRs to design and what CDR lengths or conformation combinations to sample. The computational benchmarking of AbDesign consisted of reporting the Cα RMSD from native for each CDR of the top design for each of 9 antibodies. Only the CDRs with the most common canonical conformations were reproduced; those with less common conformations were poorly predicted, making it difficult to evaluate the statistical significance of their results.

AbDesign was used recently to create lead antibodies against insulin and mycobacterial acyl-carrier protein, which were then synthesized and tested for binding [48]. Three weak binders were then subjected to random mutagenesis in a yeast-display library screen followed by manually chosen mutations, which resulted in antibodies with affinity in the 50-100 nM range. Two residues in the epitope of each of the two ACP-binding antibodies were mutated to test the designs. Only one of these four reduced binding significantly (by 75%; two others reduced binding by 10% and 20%). One mutation from valine to glutamic acid actually increased binding. Two residues outside the epitope of each of the two ACP designs were also mutated; three of these mutations increased binding of their respective antibodies 2 to 5 fold and one of them surprisingly abrogated binding. The equivocal computational benchmarking and experimental results for AbDesign suggest that further development of antibody computational design is warranted.

Taking advantage of the influx of new structures of antibodies in the PDB, we presented a new clustering of all CDR structures in the Protein Data Bank (PDB) in 2011, updating the Chothia classification developed in the 1980s and 1990s [49–52]. Our clustering was performed with a dihedral angle metric and an affinity-propagation clustering algorithm, and was presented with a systematic nomenclature [53], which is now in common use. From this classification, we developed the PyIgClassify database [54], which is updated monthly and contains CDR sequences and cluster identifications for all antibodies in the PDB. PyIgClassify includes identification of species and IMGT germline V regions [55], and is provided as a relational database for use in antibody structure prediction and design.

We hypothesized that the clusters in PyIgClassify could form the core of a knowledge-based approach to antibody design. In this paper, we test this hypothesis through a computational benchmark and experimental validation on two separate antibodies. Using the data from PyIgClassify, our main approach to design is to graft CDRs from populated clusters onto the antibody and to sample the sequence and structure space of that CDR according to the observed variation in sequence and structure of that cluster in the database. Our goal was to create a flexible, generalized antibody design framework and program that can be applied to numerous types of antibody design projects from affinity maturation to de novo design.

To create a reliable antibody design framework from our structural bioinformatics efforts, we leveraged the Rosetta Software Suite [56], a collaborative research project across many independent labs around the world. Rosetta has been developed and used for a variety of modeling and design tasks, such as loop modeling [57,58], protein–protein docking [45,59], structure refinement [60–63], de novo protein design [64], enzyme design [65–67], and interface design [68–70]. Rosetta provides frameworks for sampling and optimizing the conformations of the backbone and side chains of a protein–protein complex while simultaneously changing the sequence at specified positions in the interface (in this case, primarily in the CDRs) in order to optimize the total energy of the system. Alternatively, the program can optimize the interface energy, which is the difference between the energy of the relaxed complex and the sum of the energies of the separated components after relaxation.

The program and methodology we have developed is called RosettaAntibodyDesign or RAbD. In this paper, we describe RAbD and both experimental testing and extensive computational benchmarking. To develop RAbD: (1) we created a database of CDR structures annotated according to our CDR cluster nomenclature and added this database to Rosetta; (2) implemented user-controlled sampling of CDR structures from this database for antibody design; (3) developed new grafting methods using the cyclic coordinate descent algorithm [71] in Rosetta; (4) implemented an algorithm that utilizes sequence profiles for our CDR clusters for sampling amino acid changes during antibody design and exploits existing structure optimization and Monte Carlo design strategies in Rosetta; and (5) added antibody-specific analysis tools to Rosetta to provide data that can be used in selecting antibody designs for synthesis and testing. The RAbD Framework consists of around 50 new Rosetta classes and over 20,000 lines of code, all of which is used by the RAbD program and available in RosettaScripts.

A common method for computational benchmarking of protein design methods is the use of the concept of *sequence recovery* [72]. Sequence recovery tests how the sequences in the final design models match the native sequence, calculated as percent sequence identity for all or the subset of the designable residues. Rosetta’s sequence recovery tends to be in the 35– 40% range for full design of monomeric proteins [73], since many surface positions are tolerant to amino acid substitution, and the benchmark protocols do not include functional interactions with other proteins, nucleic acids, or ligands.

However, since our antibody design protocol includes potential changes in the overall structure of the CDRs by sampling different CDR lengths, clusters, and sequences, the standard sequence recovery metric is inadequate for testing computational antibody design. We have therefore expanded the concept of sequence recovery to include recovery of structural features of the designed antibodies. Although antibodies in the PDB are not likely to be the highest affinity possible to a given epitope, they bind strongly enough for crystallography. Thus, maximizing the recovery of CDR lengths, clusters, and sequences is a reasonable strategy to optimize sampling and scoring strategies for antibody design.

We have developed novel recovery metrics and a way of assessing the statistical significance of these metrics, which may be used in any protein design scenario. To do this, we borrow a concept from statistical epidemiology, the Risk Ratio. The Risk Ratio (RR) is defined as the ratio of two frequencies (or proportions): the frequency of event X in situation A (e.g., disease progression while taking a drug) and the frequency of event X in situation B (e.g., disease progression with no drug treatment). The Risk Ratio is similar to the odds ratio, which is simply the ratio of the odds of X to not-X in situation A and the odds of X to not-X in situation B. However, the interpretation of the odds ratio is often misleading and used incorrectly to inflate a sense of benefit or risk [74].

For antibody design, we have defined two design metrics – the *design risk ratio* and the *antigen risk ratio*. We define the *design risk ratio* as the ratio of the frequency of native CDR clusters, lengths, or residue identities in the top scoring designs divided by the frequency of the same native features sampled during the design trajectory. In this way, we can account for any uneven sampling of the native structure and sequence during the design process. The *antigen risk ratio* is the ratio of the frequency of the native CDR length, cluster, or residue identities achieved in the top scoring decoys in independent antigen-present and antigen-absent simulations. This metric accounts for any bias Rosetta may have for the native CDRs even in the absence of the antigen, possibly because of favorable framework–CDR or CDR–CDR interactions. In this paper, we utilize a benchmark of 60 κ and λ antigen-antibody complexes that we chose to be as diverse in CDR lengths and clusters as possible. We show that RAbD is able to achieve risk ratios greater than 1.0 for each CDR, and we show statistical significance of these results with 95% confidence intervals.

To enable repeatable analysis and comparison of native, modeled, and designed antibody structures output by RAbD, we developed a set of antibody-specific *FeatureReporters* and *Feature R Scripts* within the Rosetta Feature Reporter framework [73,75,76]. These have enabled comparison of antibody design strategies and benchmarks, were used in the design of the antibodies in this paper, and have aided in the general optimization of the antibody design framework.

Finally, we show results where RAbD and the feature analysis reporters were used to experimentally improve binding affinity of antibodies from two different antibody–antigen complexes.

## RESULTS

### Antibody Design Framework and Program

We created a general framework and application for antibody design within the Rosetta software suite written in C++. This highly customizable framework enables the tailored design of antibody CDRs, frameworks, and antigens using highly expanded core components of the *RosettaAntibody* framework [77–79] and our PyIgClassify clustering of antibody CDRs [53,54] as its base. As with other Rosetta design protocols, RosettaAntibodyDesign depends on a “Monte Carlo plus minimization” (MCM) procedure [80]. This means that at each stage of the simulation, a change in sequence and/or structure is sampled randomly, followed by energy minimization within the Rosetta energy function. If the resulting minimized structure (a “decoy”) has lower energy than the previous decoy in the protocol, then the new structure is accepted. If the energy of the new design is higher than the previous decoy, the new design is accepted with probability exp(-ΔE/RT) where ΔE is the change in energy. This energy can be either the total energy or the calculated interface energy, which is the energy of the complex minus the energies of the separated antigen and antibody after side-chain repacking [81], or a weighted combination of both. The RAbD algorithm samples the diverse sequence, structure, and binding space of an antibody-antigen complex (Fig. 1; Fig. A in S1 Supporting Information).

**Fig. 1.**
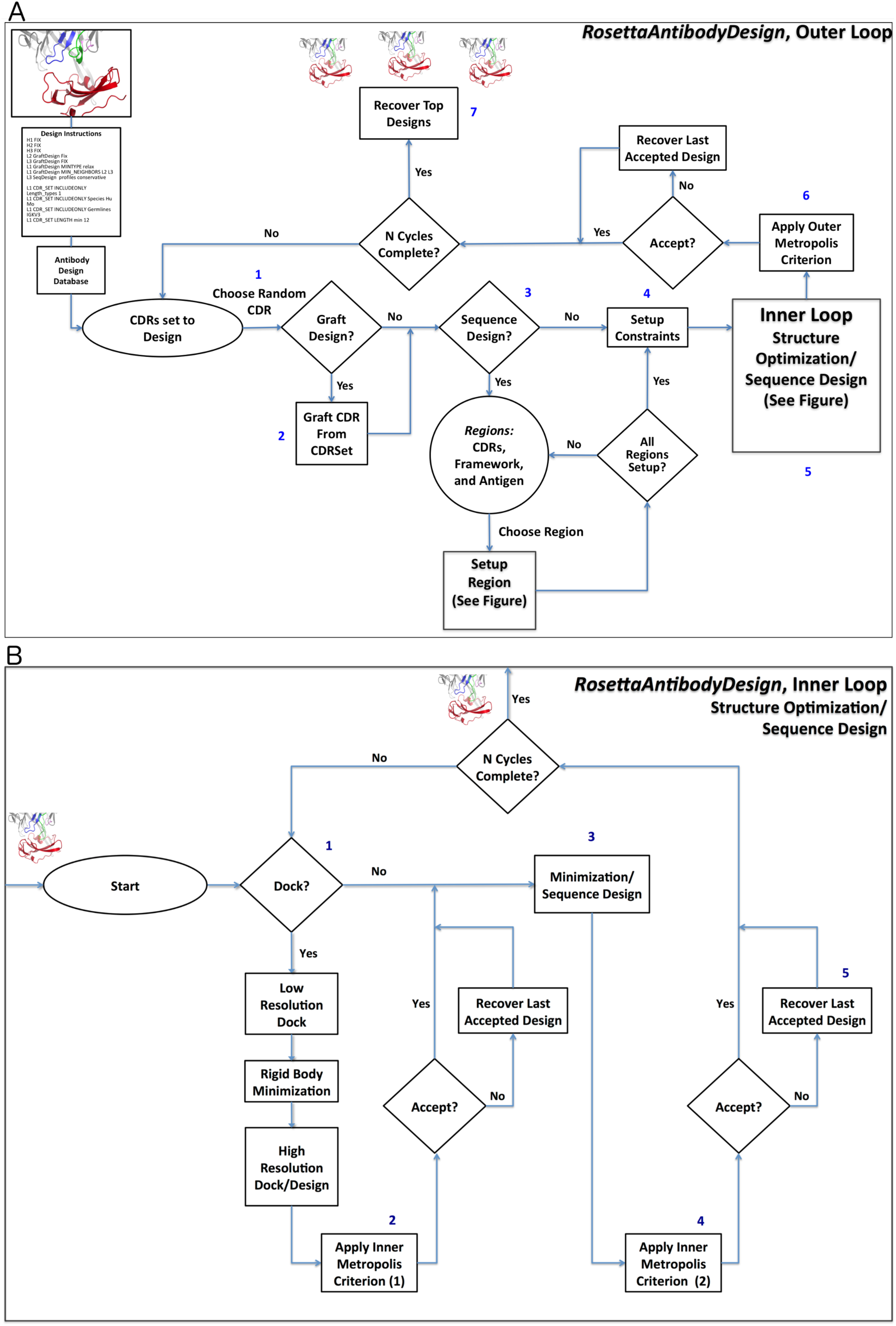
Schematic diagrams of. RosettaAntibodyDesign. A. The outer loop: The protocol starts by (1) Choosing a CDR from those that are set to design [L1, L2, etc.] randomly according to set weights (default is equal weighting) and (2) grafting a random structure for that CDR from the CDRSet, a set of CDR structures from the PDB that satisfy user-defined input rules. (3) Regional Sequence Design is then setup for all designable regions and (4) structural constraints on the CDRs and SiteConstraints on the antibody-antigen orientation, if any, are set. (5) N Inner cycles are then completed, followed by (6) the application of the Monte Carlo criterion to either accept or reject the preliminary designs. (7) Finally, the lowest energy designs are output. B. The inner loop: (1) The antigen-antibody interface is first optionally optimized by running N cycles of *RosettaDock* [45]. Interface residues set to undergo sequence design will be designed. (2) The inner Monte Carlo criterion is then applied. The conformations of the CDR, its stem, and surrounding residues, and CDRs are then optimized according to the instruction file. (3) Residues from neighboring regions are designed if enabled (Fig. 2 shows this packing/design shell). (4) The inner Monte Carlo criterion is then applied again and (5) the lowest energy decoy found in the inner loop is returned to the outer loop.

The protocol begins with the three-dimensional structure of an antibody–antigen complex. This structure may be an experimental structure of an existing antibody in complex with its antigen or a predicted structure of an existing antibody docked computationally to its antigen. As a prelude to de novo design, the best scoring results of low-resolution docking of a large number of unrelated antibodies to a desired epitope on a target antigen structure may be used. It should be noted that design on predicted structures is generally less reliable than design on high-resolution crystal structures due to possible inaccuracies in the model. The RosettaAntibodyDesign protocol is driven by a set of command-line options and an optional set of design instructions provided as an input file for increased control. Details and example command lines and instruction files are provided in the Supplemental Methods section.

RAbD enables the grafting of CDRs from diverse clusters of different lengths within the PyIgClassify database, sampling from the sequence and structural variation within each cluster. Broadly, the RAbD protocol consists of alternating outer and inner Monte Carlo cycles. Each outer cycle (of *N_outer_* cycles) (Fig. 1A) consists of randomly choosing a CDR (L1, L2, etc.) from those CDRs set to design, randomly choosing a cluster and then a structure from that cluster from the database according to the input instructions. The CDR is then grafted onto the antibody framework in place of the existing CDR (*GraftDesign*). The program then performs *N_inner_* rounds of the inner cycle (Fig. 1B), consisting of sequence design (*SeqDesign*) and local structure optimization. Sequence design is performed by Rosetta’s side-chain repacking algorithm: a residue is chosen randomly and the energy of each of its rotamers is evaluated (both internal energy and interaction with the environment); if the residue is set to be designed, then the rotamers of multiple residue types are tested; the side chain is then placed in the rotamer (and residue type) with lowest energy. This is repeated for residues in the grafted CDR as well as residues in neighboring CDRs and the framework (where only the native residue types are used). Once this design is completed, local structure optimization is performed with Rosetta’s standard local energy minimization routines. Amino acid changes are typically sampled from profiles derived for each CDR cluster in PyIgClassify. Conservative amino acid substitutions (according to the BLOSUM62 substitution matrix) may be performed when too few sequences are available to produce a profile (e.g., for H3). Each inner cycle structurally optimizes the backbone and repacks side chains of the CDR and its neighbors in order to optimize interactions of the CDR with the antigen and other CDRs (Fig. 2). Backbone dihedral angle constraints derived from the cluster data are applied to limit deleterious structural perturbations. After each inner cycle is completed, the new sequence and structure are accepted according to the Metropolis Monte Carlo criterion. After *N_inner_* rounds of the inner cycle, the program returns to the outer cycle, at which point the energy of the resulting design is compared to the previous design in the outer cycle. The new design is accepted or rejected according to the Monte Carlo criterion. After *N_outer_* cycles (default of 25), the lowest energy design observed during the run is output by the program as the final design. In practice, the whole procedure is performed in parallel on a cluster to produce 100s or 1000s of output structures (decoys). This ensemble of designs is then analyzed to choose specific sequences for experimental testing, typically based on both total energy and interface energy, which are reported in the decoys, or the needs of the specific project. Decoy discrimination, analysis, and selection are critical to the experimental success of the final designs.

**Fig. 2.**
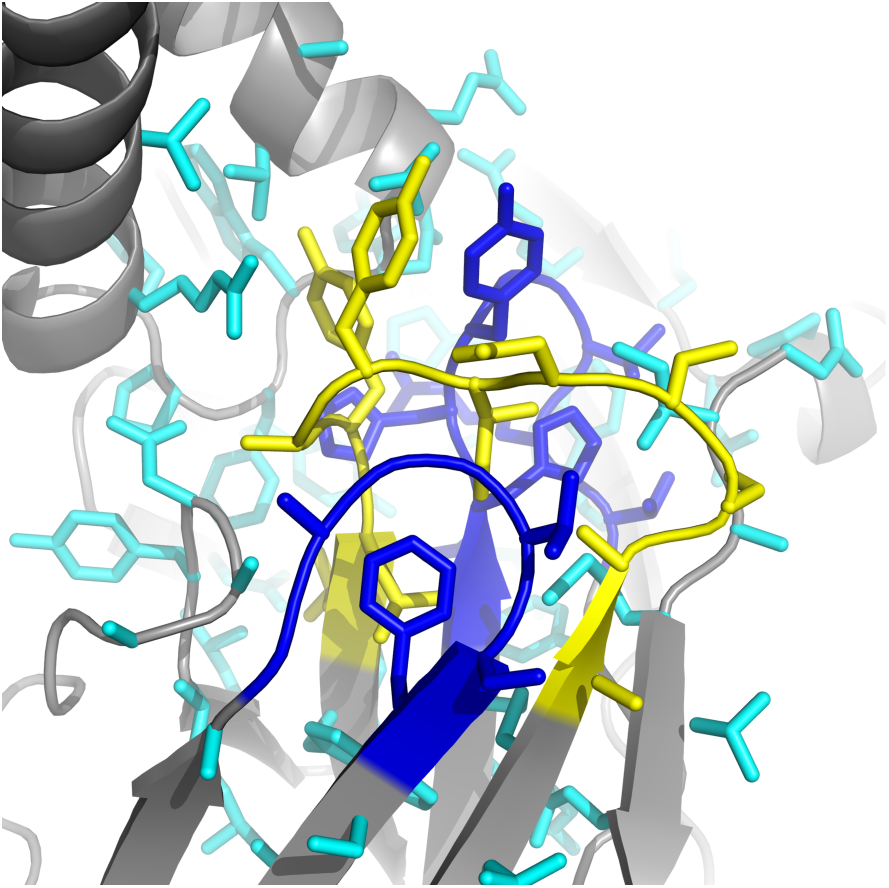
Packing shell setup. During the inner optimization cycle, a packing shell is created (cyan) around the chosen CDR (in this case, L1 in yellow), and its neighbors (in this case, L3 and the DE loop (L4) in blue). By default, 6 Å is used as the packing shell distance. During the inner loop, all side chains are optimized and amino acid changes are made to any CDRs or regions set to sequence. The chosen CDR and its neighbors additionally undergo backbone optimization during this stage according to the minimization type chosen.

RAbD can be tailored for a variety of design projects and design strategies. This is accomplished through the use of a set of command-line options and an optional CDR Instruction File. The CDR Instruction File (Fig. 3) uses a simple syntax and enables control over what lengths, clusters, germlines, and organism of each CDR will be sampled (the CDRSet) and which structural optimizations are used to minimize the score of each design. Each instruction can be set for all of the CDRs using a specific keyword, or they can be set individually. For example, in a redesign project, we may want to design an antibody with a particular CDR that is longer than the existing CDR in order to create new contacts with the antigen that are not present in the starting structure. Alternatively, we may simply want to optimize the sequence of a particular CDR or set of CDRs using the cluster profiles from PyIgClassify. These examples can be accomplished easily through the CDR Instruction File, and this flexibility has been used to design the antibodies described below.

**Fig. 3.**
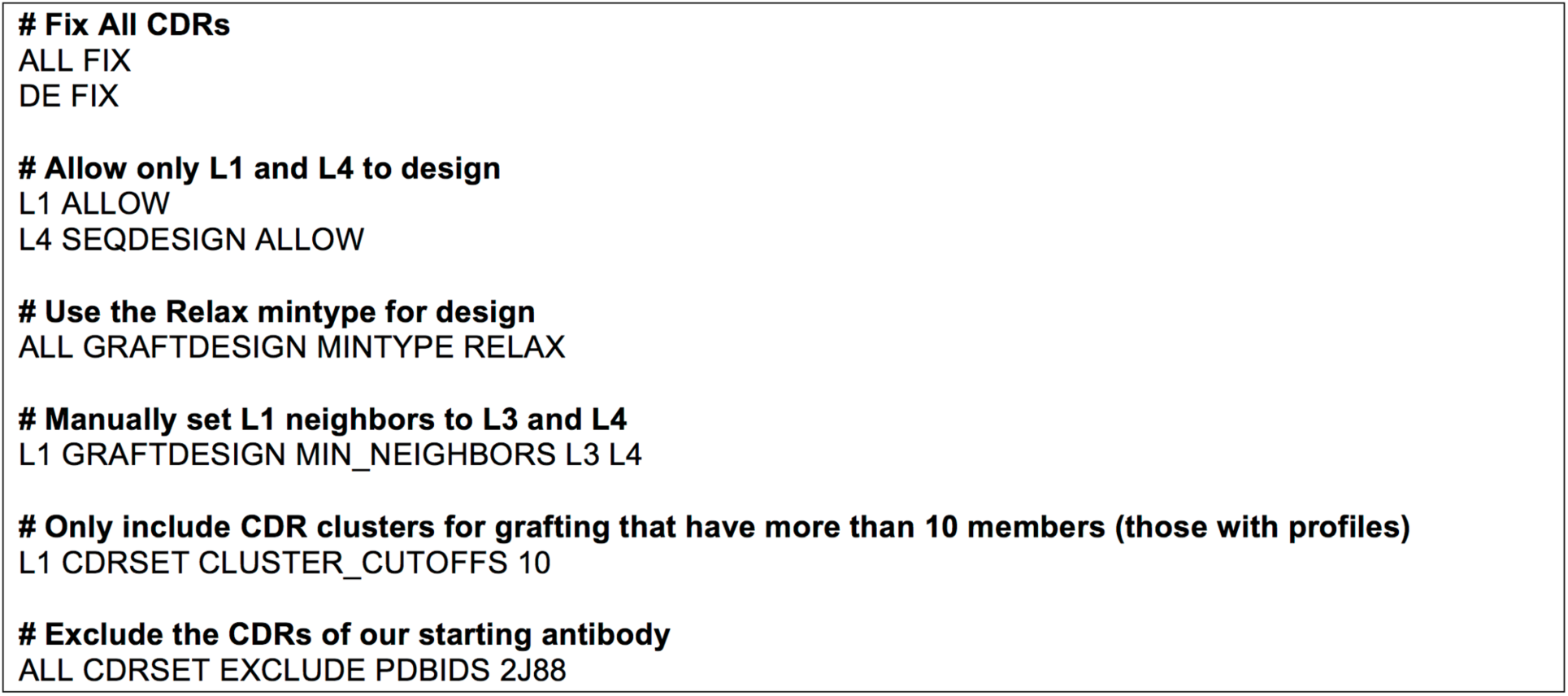
Example CDR Instruction file used for successful 2J88 antibody design L14_7. Lines beginning with # are comments and are ignored by the program. Further details are provided in Methods.

A core component of the RAbD protocol is an SQLITE3 antibody design database that houses all structures, CDR-clustering information, species, germline, and sequence profile data used for design. The database benchmarked in this paper comes from the August 2017 release of PyIgClassify, but up-to-date versions that reflect the current Protein Data Bank (PDB) can also be obtained from the PyIgClassify website (http://dunbrack2.fccc.edu/PyIgClassify). If RAbD uses non-redundant databases without outliers (the default), defined as CDRs greater than 40° or 1.5 Å RMSD from one of our cluster centroids (not applied to H3), this database comprises 657 L1 sequences, 471 L2 sequences, 681 L3 sequences, 805 H1 sequences, 930 H2 sequences, and 985 H3 sequences. In order to improve framework-CDR compatibility in the final designs, λ and κ type antibodies are designed by limiting the resulting CDRSet to only those CDRs derived from the same light chain type as the antibody undergoing design (Fig. 2).

### Computational Benchmarking of RAbD

To develop metrics for recovery of CDR lengths and clusters, we must account for the fact that CDR lengths and clusters are not evenly distributed in nature, the PDB, or in the PyIgClassify database and are not necessarily sampled evenly during RAbD’s Monte-Carlo trajectories. The probability of choosing the native cluster and length during sampling directly influences the statistical significance of the final recovery of the native length and cluster.

To account for this phenomenon, we borrow a concept from statistical epidemiology, the Risk Ratio. The Risk Ratio (RR) is defined as the ratio of two frequencies: the frequency of event X in situation A (e.g., disease progression while taking a drug) and the frequency of event X in situation B (e.g., disease progression with no drug treatment). The Risk Ratio is similar to the odds ratio, which is simply the ratio of the odds of X to not-X in situation A and the odds of X to not-X in situation B. However, the interpretation of the odds ratio is often misleading and used incorrectly to inflate a sense of benefit or risk [74]. In standard protein design scenarios, we may define the risk ratio as the frequency of the native structure (or sequence) in the top scoring designs divided by the frequency of the native structure (or sequence) sampled during the protocol. If we perform design simulations on an existing high-affinity antibody–antigen complex, it is reasonable to suppose that a successful protocol will recover the native CDR lengths, conformations, and sequences of a high-affinity antibody more often than they are sampled. We therefore define the design risk ratio (DRR) for CDR lengths and clusters as the frequency of the native length or cluster in the top scoring designs (the top decoys, one from each run of the program) divided by the frequency that the native length or cluster was sampled during the design simulations.

Because Rosetta might prefer some CDR conformations and lengths because they are lower energy, even in the absence of antigen, we also define, the antigen risk ratio (ARR), which is the frequency of the native CDR length or cluster in the top scoring designs in the presence of the antigen divided by the frequency of the native in the top scoring designs from independent simulations performed in the absence of the antigen. It is straightforward to calculate confidence intervals for the design and antigen risk ratios so that statistical significance of the results can be assessed (see Methods).

We tested two types of design methods: ‘opt-E’, which uses the Metropolis Monte Carlo criterion to optimize Total Rosetta Energy of the antibody-antigen complex, and ‘opt-dG’, which optimizes the calculated interface energy. The interface energy is equal to the Total Rosetta Energy of the complex minus the Total Rosetta Energy of the separated antigen and antibody, after side-chain repacking. For the opt-E method, we calculate both the DRR and ARR values. Since opt-dG includes a step of separating the antigen and antibody, an antigen-free simulation is not relevant to the calculation, and we therefore only calculate the DRR for the opt-dG designs. All 5 non-H3 CDRs were graft-designed, while all CDR sequences, including H3, were sequence-designed either preferentially using derived CDR cluster profiles or conservative design where cluster sequence data were sparse. All 5 non-H3 CDRs began each simulation with randomly inserted CDRs from the antibody design database. Prior to design calculations, the structure of each antigen-antibody complex was minimized into the Rosetta energy function with tight coordinate constraints on both backbone and side-chain regions [62] (see Methods for protocol). We used an up-to-date version of the antibody design database derived from the PDB as of August 2017. It contains 3,974 CDRs, while our original clustering in North et al. contained 1,346 CDRs (http://dunbrack2.fccc.edu/pyigclassify).

A diverse set of 46 κ and 14 λ antibody–antigen complexes were used for the computational benchmarks (Table 1; more details on the benchmark antibodies are provided in Table A in S1 Supporting Information). This set of antibody–antigen complexes includes a diverse set of CDR lengths and clusters, with many of the clusters commonly found in the PDB. The benchmark complexes were selected to satisfy several criteria: (1) resolution ≤ 2.5 Å; (2) buried surface area in the antigen-antibody complex > 700 Å^2^; (3) CDR1 and CDR2 within 40° of one of our cluster centroids; (4) contacts with CDRs in both the light chain and the heavy chain variable domains; (5) non-redundancy – antibodies which bind the same antigen were only selected if they bound to completely different sites on the antigen; 6) benchmark antibodies were prioritized so as to comprise as diverse a set of CDR lengths and clusters given the distribution of lengths and clusters present in the PDB. The benchmark contains 22 length classes and 35 clusters over the 5 non-H3 CDRs and lengths of H3 from 6 to 24 residues.

**Table 1.** Benchmark antibody complexes

|  | PDB | VL | H1 | H2 | H3 | L1 | L2 | L3 | Ag Length | Antigen |
| --- | --- | --- | --- | --- | --- | --- | --- | --- | --- | --- |
| 1 | 1a14 | $\kappa$ | H1-13-1 | H2-10-1 | H3-15 | L1-11-2 | L2-8-1 | L3-9-cis7-1 | 388 | Neuraminidase |
| 2 | 1a2y | $\kappa$ | H1-13-1 | H2-9-1 | H3-10 | L1-11-2 | L2-8-1 | L3-9-cis7-2 | 129 | Lysozyme C |
| 3 | 1fe8 | $\kappa$ | H1-13-1 | H2-9-1 | H3-9 | L1-11-1 | L2-8-1 | L3-9-cis7-1 | 196 | von Willebrand factor |
| 4 | 1ic7 | $\kappa$ | H1-13-1 | H2-9-1 | H3-7 | L1-11-1 | L2-8-1 | L3-9-cis7-1 | 129 | Lysozyme C |
| 5 | 1iqd | $\kappa$ | H1-13-1 | H2-10-1 | H3-10 | L1-12-1 | L2-8-1 | L3-9-1 | 156 | Coagulation factor VIII |
| 6 | 1n8z | $\kappa$ | H1-13-1 | H2-10-1 | H3-13 | L1-11-1 | L2-8-1 | L3-9-cis7-1 | 607 | ErbB-2 |
| 7 | 1ncb | $\kappa$ | H1-13-1 | H2-10-1 | H3-13 | L1-11-2 | L2-8-1 | L3-9-cis7-1 | 389 | Neuraminidase |
| 8 | 1osp | $\kappa$ | H1-13-7 | H2-9-3 | H3-14 | L1-11-2 | L2-8-2 | L3-9-cis7-1 | 257 | Ozd[ A |
| 9 | 1uj3 | $\kappa$ | H1-13-1 | H2-10-1 | H3-10 | L1-11-2 | L2-8-1 | L3-9-cis7-1 | 205 | Tissue factor |
| 10 | 1w72 | $\lambda$ | H1-13-1 | H2-10-2 | H3-15 | L1-11-3 | L2-8-1 | L3-11-1 | 274 | HLA-A1,b2-MG,peptide |
| 11 | 2adf | $\kappa$ | H1-13-1 | H2-10-1 | H3-11 | L1-11-2 | L2-8-1 | L3-8-1 | 196 | von Willebrand factor |
| 12 | 2b2x | $\kappa$ | H1-13-1 | H2-9-1 | H3-12 | L1-10-1 | L2-8-1 | L3-9-cis7-1 | 223 | Integrin alpha-1 |
| 13 | 2cmr | $\kappa$ | H1-13-3 | H2-10-1 | H3-12 | L1-11-1 | L2-8-1 | L3-9-cis7-1 | 226 | gp41 |
| 14 | 2dd8 | $\lambda$ | H1-13-10 | H2-10-1 | H3-11 | L1-11-3 | L2-8-1 | L3-10-1 | 202 | Spike glycoprotein |
| 15 | 2ghw | $\kappa$ | H1-13-1 | H2-10-2 | H3-10 | L1-11-1 | L2-8-1 | L3-9-cis7-1 | 203 | Spike glycoprotein |
| 16 | 2vxt | $\kappa$ | H1-13-1 | H2-10-1 | H3-6 | L1-11-1 | L2-8-1 | L3-9-cis7-1 | 157 | Interleukin-18 |
| 17 | 2xqy | $\kappa$ | H1-13-1 | H2-10-1 | H3-11 | L1-15-1 | L2-8-1 | L3-9-cis7-1 | 572 | Envelope glycoprotein-H |
| 18 | 2xwt | $\lambda$ | H1-13-1 | H2-10-1 | H3-12 | L1-13-1 | L2-8-2 | L3-11-1 | 239 | Thyrotropin receptor |
| 19 | 2ypv | $\kappa$ | H1-13-1 | H2-10-1 | H3-12 | L1-11-2 | L2-8-1 | L3-9-cis7-1 | 253 | Lipoprotein |
| 20 | 3bn9 | $\kappa$ | H1-13-1 | H2-10-2 | H3-21 | L1-11-1 | L2-8-1 | L3-9-cis7-1 | 241 | MT-SP1 |
| 21 | 3cx5 | $\kappa$ | H1-14-1 | H2-9-1 | H3-15 | L1-11-2 | L2-8-1 | L3-9-cis7-1 | 185 | Rieske Iron-sulfur protein |
| 22 | 3ffd | $\lambda$ | H1-13-1 | H2-10-2 | H3-11 | L1-12-3 | L2-12-2 | L3-13-1 | 108 | PTH-related |
| 23 | 3h3b | $\kappa$ | H1-13-1 | H2-10-1 | H3-13 | L1-17-1 | L2-8-1 | L3-9-cis7-1 | 194 | ErbB-2 |
| 24 | 3hi6 | $\kappa$ | H1-13-1 | H2-10-2 | H3-13 | L1-11-1 | L2-8-1 | L3-8-1 | 180 | Integrin alpha-L |
| 25 | 3k2u | $\kappa$ | H1-13-1 | H2-10-1 | H3-11 | L1-11-1 | L2-8-1 | L3-9-cis7-1 | 257 | HGF activator |
| 26 | 3l95 | $\kappa$ | H1-13-1 | H2-10-1 | H3-12 | L1-11-1 | L2-8-1 | L3-9-2 | 244 | NOTCH1 |
| 27 | 3mxw | κ | H1-13-1 | H2-10-1 | H3-12 | L1-11-1 | L2-8-1 | L3-9-cis7-1 | 169 | Sonic hedgehog protein |
| 28 | 3nid | κ | H1-13-1 | H2-10-1 | H3-12 | L1-11-2 | L2-8-1 | L3-9-cis7-1 | 457 | Integrin alpha-IIb |
| 29 | 3o2d | κ | H1-13-1 | H2-10-1 | H3-15 | L1-17-1 | L2-8-1 | L3-8-1 | 188 | CD4 |
| 30 | 3rkd | κ | H1-15-1 | H2-9-1 | H3-16 | L1-11-2 | L2-8-1 | L3-9-cis7-2 | 146 | Capsid protein |
| 31 | 3s35 | κ | H1-13-1 | H2-10-1 | H3-10 | L1-15-1 | L2-8-1 | L3-9-cis7-1 | 122 | VGFR2 |
| 32 | 3uzq | κ | H1-13-1 | H2-10-1 | H3-9 | L1-15-1 | L2-8-1 | L3-9-cis7-1 | 114 | Genome polyprotein |
| 33 | 3w9e | κ | H1-13-1 | H2-10-1 | H3-15 | L1-12-1 | L2-8-1 | L3-8-2 | 306 | Envelope glycoprotein D |
| 34 | 4cmh | κ | H1-13-1 | H2-10-1 | H3-13 | L1-11-1 | L2-8-1 | L3-9-cis7-1 | 256 | CD38 |
| 35 | 4dtg | κ | H1-13-1 | H2-10-2 | H3-14 | L1-16-1 | L2-8-1 | L3-9-cis7-1 | 66 | TFPI |
| 36 | 4dvr | κ | H1-13-1 | H2-10-1 | H3-12 | L1-11-1 | L2-8-1 | L3-8-1 | 313 | gp160 |
| 37 | 4etq | κ | H1-13-1 | H2-10-1 | H3-12 | L1-10-1 | L2-8-1 | L3-9-cis7-1 | 269 | IMV membrane protein |
| 38 | 4ffv | κ | H1-13-1 | H2-10-1 | H3-10 | L1-10-1 | L2-8-4 | L3-9-cis7-1 | 730 | Dipeptidyl peptidase 4 |
| 39 | 4fqj | λ | H1-13-1 | H2-10-1 | H3-18 | L1-13-1 | L2-8-1 | L3-11-1 | 304 | Hemagglutinin |
| 40 | 4g6j | κ | H1-13-1 | H2-10-2 | H3-11 | L1-11-1 | L2-8-1 | L3-9-cis7-1 | 158 | Interleukin-1 beta |
| 41 | 4g6m | κ | H1-15-1 | H2-9-1 | H3-12 | L1-11-2 | L2-8-1 | L3-9-cis7-1 | 150 | Interleukin-1 beta |
| 42 | 4h8w | λ | H1-13-1 | H2-10-2 | H3-12 | L1-14-2 | L2-8-1 | L3-11-1 | 353 | gp160 |
| 43 | 4ki5 | κ | H1-13-1 | H2-10-1 | H3-15 | L1-11-2 | L2-8-2 | L3-9-cis7-1 | 183 | Factor VIII |
| 44 | 4lvn | κ | H1-14-1 | H2-9-1 | H3-13 | L1-12-1 | L2-8-1 | L3-9-cis7-1 | 344 | Subtilisin-like SP |
| 45 | 4ot1 | λ | H1-13-1 | H2-10-1 | H3-24 | L1-13-1 | L2-8-2 | L3-10-1 | 129 | Envelope glycoprotein B |
| 46 | 4qci | λ | H1-13-1 | H2-10-2 | H3-13 | L1-11-3 | L2-8-1 | L3-9-2 | 110 | PDGFR Beta |
| 47 | 4xnq | λ | H1-14-1 | H2-9-1 | H3-16 | L1-11-3 | L2-8-1 | L3-9-1 | 212 | Hemagglutinin (Fragment) |
| 48 | 4ydk | κ | H1-13-1 | H2-10-2 | H3-22 | L1-11-1 | L2-8-1 | L3-9-2 | 353 | gp160 |
| 49 | 5b8c | κ | H1-13-1 | H2-10-1 | H3-13 | L1-15-1 | L2-8-1 | L3-9-cis7-1 | 139 | PD1 |
| 50 | 5bv7 | λ | H1-13-1 | H2-10-2 | H3-19 | L1-11-3 | L2-8-1 | L3-10-1 | 422 | PC-sterol acyltransferase |
| 51 | 5d93 | κ | H1-13-1 | H2-10-1 | H3-9 | L1-10-1 | L2-8-1 | L3-9-cis7-1 | 244 | Sulfhydryl oxidase 1 |
| 52 | 5d96 | κ | H1-13-1 | H2-9-1 | H3-12 | L1-11-1 | L2-8-1 | L3-9-cis7-1 | 244 | Sulfhydryl oxidase 1 |
| 53 | 5en2 | κ | H1-13-1 | H2-10-1 | H3-17 | L1-11-1 | L2-8-1 | L3-9-cis7-1 | 141 | Pre-glycoprotein GP |
| 54 | 5f9o | κ | H1-13-1 | H2-10-1 | H3-15 | L1-11-1 | L2-8-1 | L3-8-1 | 352 | gp120 core |
| 55 | 5ggs | κ | H1-13-3 | H2-10-1 | H3-13 | L1-15-1 | L2-8-1 | L3-9-cis7-1 | 123 | PD1 |
| 56 | 5hi4 | λ | H1-13-1 | H2-10-2 | H3-11 | L1-13-2 | L2-8-1 | L3-9-1 | 132 | Interleukin-17A homodimer |
| 57 | 5j13 | λ | H1-13-1 | H2-10-2 | H3-15 | L1-11-3 | L2-8-2 | L3-11-1 | 147 | Thymic stromal lymphopoietin |
| 58 | 5l6y | λ | H1-13-1 | H2-10-1 | H3-15 | L1-11-3 | L2-8-1 | L3-11-1 | 112 | Interleukin-13 |
| 59 | 5mes | λ | H1-13-1 | H2-10-2 | H3-12 | L1-13-1 | L2-8-1 | L3-11-1 | 162 | Mcl-1 homolog |
| 60 | 5nuz | κ | H1-13-1 | H2-10-1 | H3-13 | L1-15-1 | L2-8-1 | L3-9-cis7-1 | 156 | Pre-glycoprotein GP |

For each CDR, the PyIgClassify cluster is given. For H3, only the length is given. VL provides the kappa or lambda identify of the light chain variable domain of the antibody. Antibodies to the same antigen bind in different locations and are not the same antibody.

We define the “%Sampled” as the rate at which the native length or cluster is sampled during the design trajectories. The 5 non-H3 CDRs are very different in terms of the diversity of lengths and clusters that are observed in the PDB [53], with L2 and H1 having more than 90% of CDRs in the PDB with a single length and conformation (clusters L2-8-1 and H1-13-1 respectively), while L1, L3, and H2 are more diverse in both length and conformation. We ran design simulations for the antibodies in the benchmark set by sampling the clusters of each CDR evenly (regardless of length) of all clusters represented in the database by 5 or more unique sequences from the same antibody gene (heavy, λ, or κ antibody CDRs). Thus, the %Sampled of native CDR *lengths* (Fig. 4A) is only 23% for the most length-diverse CDR, L1; followed by L3 (50%) and H2 and H1 (both 66%), and L2 (92%), which is the least diverse (a few λ L2 CDRs are length 12). The %Sampled of native CDR *clusters* is only 10-14% for L1, L3, H1, and H2 and 34% for L2 (Fig. 4B).

**Fig. 4.**
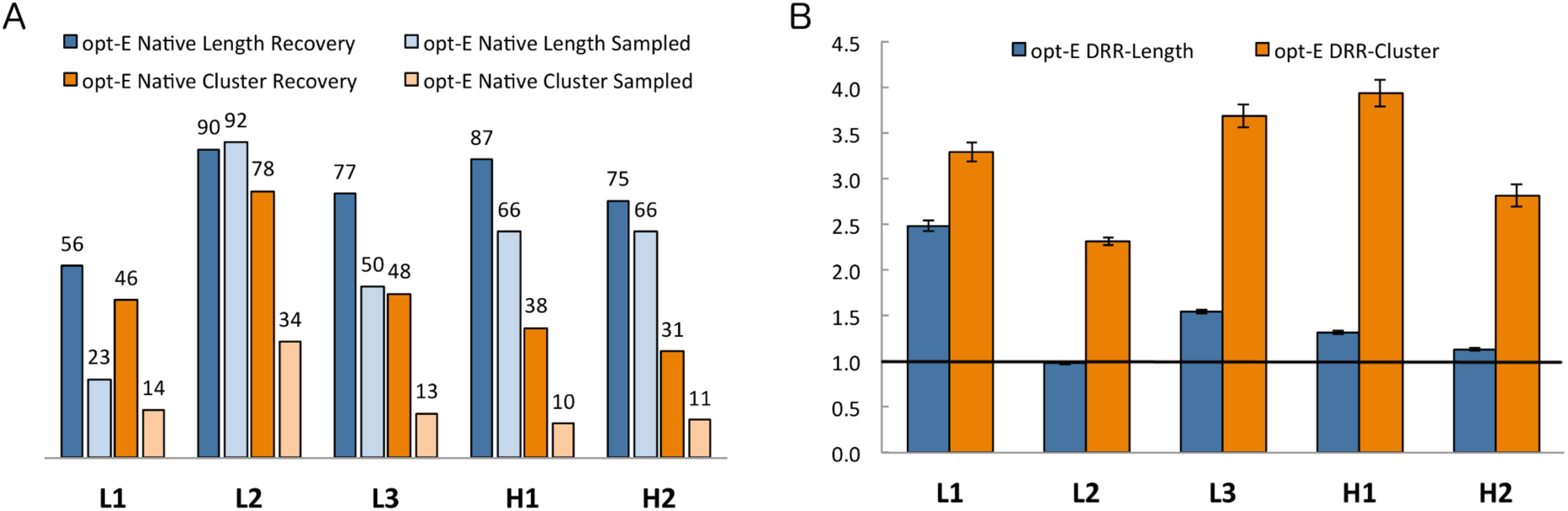
Computational benchmarking of the opt-E protocol. Recovery metrics on 60 antibodies for the opt-E protocol (optimization of total Rosetta energy) for each CDR that underwent GraftDesign in the RAbD design protocol. (A) %Recovered and %Sampled for each CDR length and cluster for the opt-E simulations. (B) Design risk ratios (DRR) for recovery of CDR length and cluster for the opt-E simulations. 95% confidence intervals for the Risk Ratio statistics are calculated as described in Methods.

For each of the 60 antibodies, we ran 100 design trajectories, each with 100 outer design cycles (Fig. 1A) for each experiment (representing a total of 10,000 full design cycles for each antibody) and analyzed the lengths and clusters of the final decoy from each of the 100 Monte Carlo simulations. The %Recovered is then the number of final decoys out of 100 runs that contain the native length or cluster (Fig. 4A) for any given CDR. The %Recovered of length generally runs parallel with the %Sampled with the least length-diverse CDRs (L2 and H1) having higher length recovery than the others. The highest cluster recovery is for L2.

The Design Risk Ratio (DRR) is then defined by Equation 1:

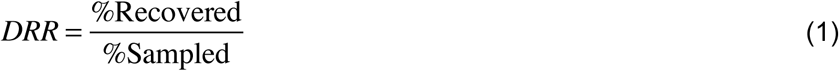

where %Recovered and %Sampled are calculated over the 100 output decoys for all 60 antibodies (6000 total). A DRR greater than 1 indicates that the length or cluster was present in the output decoys more frequently that it was sampled during the trajectories. The DRRs for the length of CDRs are highest for L1 and H3 with values of 2.5 and 1.5 respectively. We do not expect high DRRs for L2, H2, and H1 since their length diversity in the PDB is very limited in the first place.

The DRRs for the clusters are much higher. For the opt-E protocol, the cluster risk ratios are above 2.4 for all 5 non-H3 CDRs, and over 3.5 for L3 and H1. The results demonstrate the utility of the DRR in accounting for the different levels of diversity in length and cluster across the five CDRs in which *GraftDesign* was enabled. This result may come from both more favorable interactions and higher shape complementarity with the antigen–antibody interface using the native cluster(s), as well as local CDR–CDR interactions, which help to enrich certain lengths and clusters together.

There is a possibility that Rosetta scores some CDR structures more favorably than others because of internal interactions within the CDR or interactions with other CDRs or the framework. Some rare clusters may be high in energy or even artifacts of highly engineered antibodies or errors in structure determination. To investigate this, we performed the opt-E protocol without the antigen present in the simulations. We calculated an antigen risk ratio (ARR) from Equation 2 as the ratio of the frequency of the native length or cluster in the final decoys from the antigen-present simulations and the frequency of the native in the final decoys from the antigen-absent simulations:

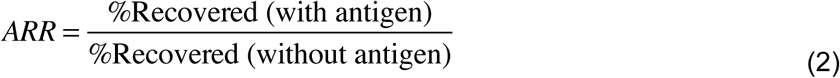

where %Recovered (with antigen) and %Recovered (without antigen) are calculated from 100 design decoys of 60 antibodies (6000 structures each). The recovery values (Fig. 5A) in the presence of antigen are all higher than the recovery values in the absence of antigen, with the exception of L3 where they are approximately equal. This is reflected in the ARR results (Fig. 5B) antigen risk ratios demonstrate that the native lengths and clusters are enriched particularly for the L1 and H2 CDRs in the presence of the antigen. For the other CDRs, the values are a little over 1.0, indicating that Rosetta prefers some of the more common clusters in the PDB, even in the absence of antigen. For the light-chain CDRs, we sampled only from lengths and clusters that contained at least 5 examples of structures from the same light-chain type, either λ or κ. Especially for L3, this choice strongly restricts the number of applicable lengths and clusters, and thus the antigen risk ratios for L3, like H1 and L2, are lower that one might expect otherwise.

**Fig. 5.**
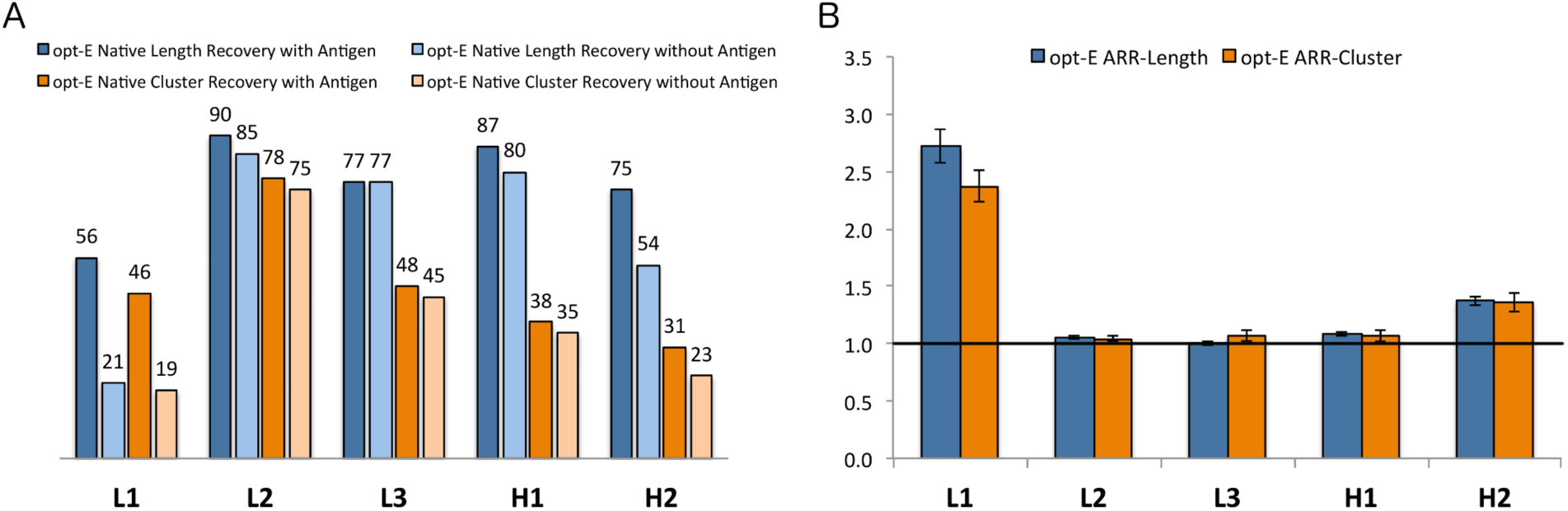
Antigen Risk Ratios for the opt-E protocol. Risk Ratios of benchmarks showing the enrichment in the recovery of native lengths and clusters in the presence of the native antigen compared to simulations performed in its absence. (A) %Recovered length and cluster for the simulations in the presence and absence of antigen. (B) Length and Cluster Antigen Risk Ratios (ARRs) A risk ratio greater than 1.0 indicates enrichment of the native length and cluster in the presence of the antigen over simulations performed in the absence of the antigen.

In addition to optimizing the total energy, design simulations in RAbD can alternatively optimize the interface energy, which for our purposes is defined as the total Rosetta energy of the antibody-antigen complex minus the energy of the separated antibody and antigen after repacking and minimizing the energy of side-chain conformations of the interface residues. The %Recovered is greater than the %Sampled (Fig 6A) for the lengths and clusters of all 5 CDRs, which is reflected in the DRR values. The cluster DRRs are greater than 1.5 for all 5 CDRs, while the length DRRs are only significantly above 1.0 for L1 which is the most length-diverse CDR for both κ or λ antibodies.

**Fig. 6.**
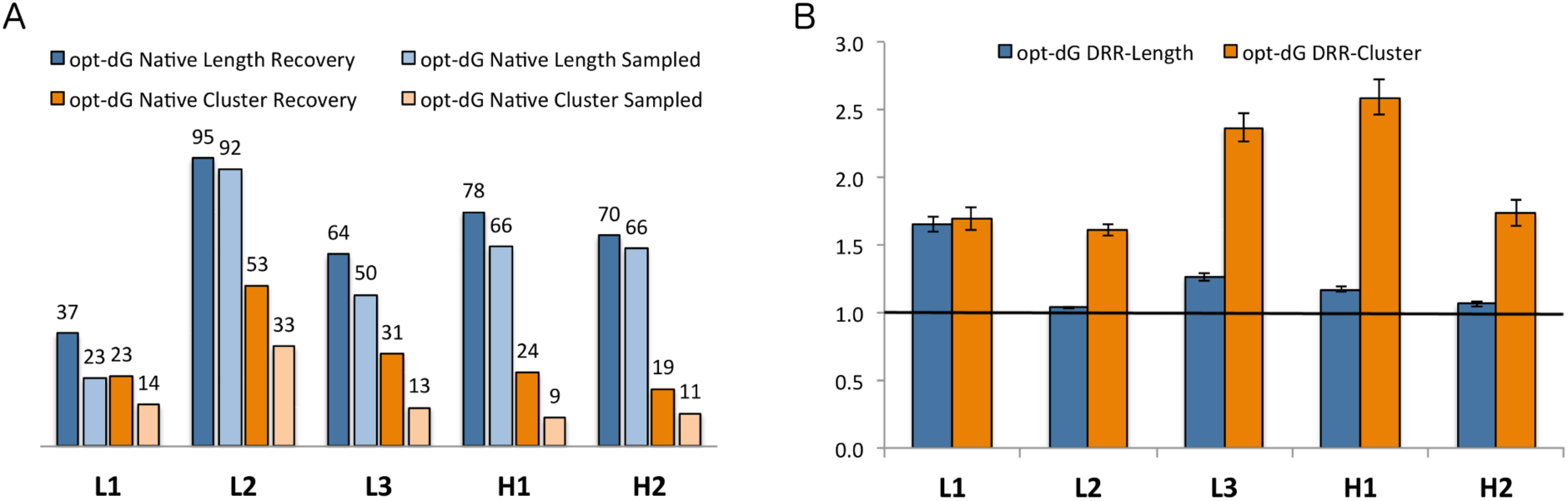
Computational benchmarking of the opt-dG antibody design protocol. Recovery metrics on 60 antibodies for the opt-dG protocol (optimization of Rosetta interface energy) for each CDR that underwent GraftDesign in the RAbD design protocol. (A) %Recovered and %Sampled for each CDR length and cluster for the opt-E simulations. (B) Design risk ratios (DRR) for recovery of CDR length and cluster for the opt-dG simulations. 95% confidence intervals for the Risk Ratio statistics are calculated as described in Methods.

To further investigate the effect of the antigen’s presence during the design phase, we performed sequence design on one CDR at a time (6 per target, including H3) starting from the native sequence and structure with and without the antigen present using the opt-E protocol described above (the opt-dG protocol does not make sense in the absence of the antigen and there is no straightforward way to calculate the sampling rate of amino acid types during the simulations). In each of these twelve simulations (6 CDRs with and without antigen), we produced 100 models for analysis. Fig. 7 shows the sequence recovery and antigen risk ratios separately for residues in contact with the antigen in the starting structures and those not directly in contact with the antigen in the starting complexes. The risk ratios were calculated from Eq. 3:

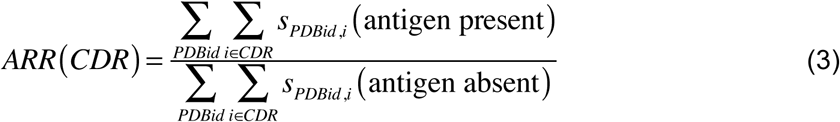

where *s_PDBid,i_* is the fraction of 100 decoys that have the native residue at position *i* of the given CDR in each PDBid.

**Fig. 7.**
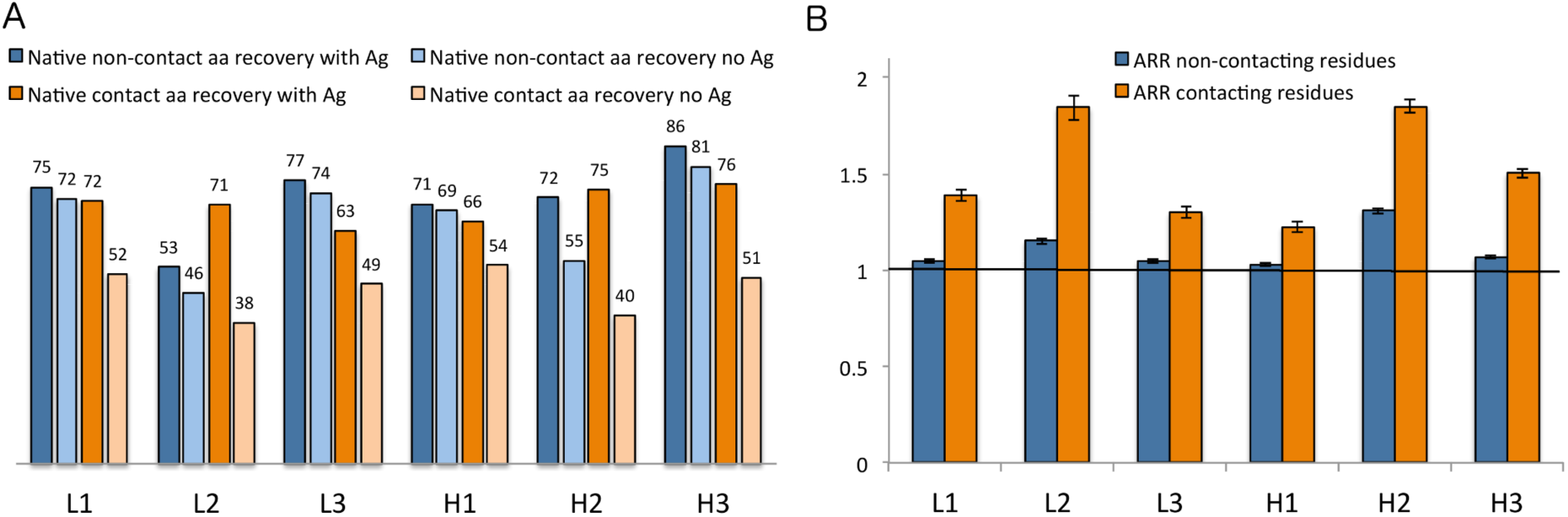
Sequence design with the opt-E protocol on the 60 antibody benchmark. (A) Sequence recovery for amino acids in contact with the antigen and those not in contact with the antigen from the antigen-present and antigen-absent simulations. (B). Antigen risk ratios (ARRs) for the contacting and non-contacting residues. Values greater than 1.0 indicate that the native residue types were present in the design simulations in the presence of the antigen more often than they were present in the design simulations in the absence of the antigen.

For the non-contacting residues in all of the CDRs, most of which are part of the CDR anchors or buried in the hydrophobic core of the variable domains, the sequence recovery rate is 73% during simulations in the presence of the antigen and 67% during simulations in the absence of the antigen. This is an overall antigen risk ratio of 1.084. The resulting ARR values are near 1.0 for all six CDRs (Fig. 7A). Many of these residues are strongly conserved in the PyIgClassify profiles, and their recovery with and without the antigen present is expected.

By sharp contrast, residues in contact with the antigen have a lower recovery of only 48% in the absence of the antigen but a much higher recovery rate of 72% in the presence of the antigen. This is an overall antigen risk ratio for the antigen-contacting residues of 1.50 (95%CI = [1.489, 1.514]). The contact residues ARRs range from 1.2 to 1.9 for the six CDRs (Fig. 7B). Since H3 contributes many residues that contributed to binding free energy, it is gratifying that the H3 risk ratio is above 1.5 and that H3 has the highest sequence recovery rate with the antigen (Fig. 7A).

We investigated the physical properties of the designed antibody-antigen complexes resulting from the opt-E and opt-dG benchmarks. As expected, the opt-dG simulations result in lower interface energies than the opt-E simulations and nearly the same as the native antigen-antibody complexes (Fig. B in S1 Supporting Information). The total energies of the opt-E and opt-dG simulations are relatively similar to each other and somewhat higher than the natives (Fig. C in S1 Supporting Information). The shape complementarities and surface areas of the designed antibody-antigen complexes are also very close to the native structures, with the opt-dG showing a slight improvement over the opt-E simulations (Fig. D and Fig. E in S1 Supporting Information).

#### Experimental Validation

Although computational benchmarking can be extremely useful in optimizing a protocol and its parameters for protein design, the true measure of new protein design methodologies is to test computationally derived sequences experimentally by expressing and purifying the proteins and testing them for desired functionality, including binding affinity and biophysical properties of the designed molecules.

We tested a common scenario for which RAbD was intended – improving the affinity of an existing antigen–antibody complex by grafting new CDRs in place of one or more of the native CDRs. To this end, we tested the ability of the RAbD program to create viable antibody designs that improve binding affinity in two antibody–antigen complexes: an HIV-neutralizing antibody known as CH103 (PDB: 4JAN) [82] that binds to the CD4 binding site of HIV gp120, and an antibody that binds to the enzyme hyaluronidase, which is the main allergen in bee venom (PDB: 2J88) [83]. These antibodies are not dominated by interactions of H3 with the antigen and use common canonical clusters for the CDRs at the binding interface. Using this knowledge and the general knowledge of CDR length and cluster variability, we designed both L1 and L3 together, or H2 in the CH103 antibody. For the 2J88 antibody, we designed either L1 and the light chain DE loop, or H2. The DE loop is a short loop between strands D and E of the variable domain β sheet (residues 82-89 in AHo numbering). The ability of RAbD to treat both the heavy- and light-chain DE loops as CDRs, which are typically considered framework regions, was added later in program development after the elucidation of the role of the loop in both antigen binding and stabilization, especially in regard to intra-CDR contacts with L1 [84]. In light of this, with the L1 design of the 2J88 antibody, we enabled sequence design (but not graft design) of this light chain DE loop, which we call L4.

The CDRs selected for design were set to undergo graft- and sequence-sampling with the relax protocol, allowing for new lengths and clusters in the final antibody design, while the framework residues and antigen residues held their starting amino acid identities. The crystal structures 2J88 and 4JAN were first minimized into the Rosetta energy function before being used as starting points for the designs. An instruction file was given for each CDR design strategy (L1 + L4, H2, and L1 + L3) with different algorithms and selection methods used to choose the final designs. Details of these settings, instruction files, and design and selection strategies can be found in Methods.

For 2J88, 30 designs were chosen, expressed, and purified with no detectable aggregation. These 30 were chosen with no visual or manual inspection, and based purely on a ranking of physical characteristics of each decoy determined by the *AntibodyFeature* reporters for each design strategy and selection characteristic (Methods). Of these 30, 20 showed some degree of binding affinity through an acceptable kinetic sensorgram signal of at least μM binding consisting of L1 loops of length 11, 15, and 17 residues (wild-type: 11) in addition to a single H2 design of 12 residues (wild-type: 9) (Fig. 8A, Table B in S1 Supporting Information).

**Fig. 8.**
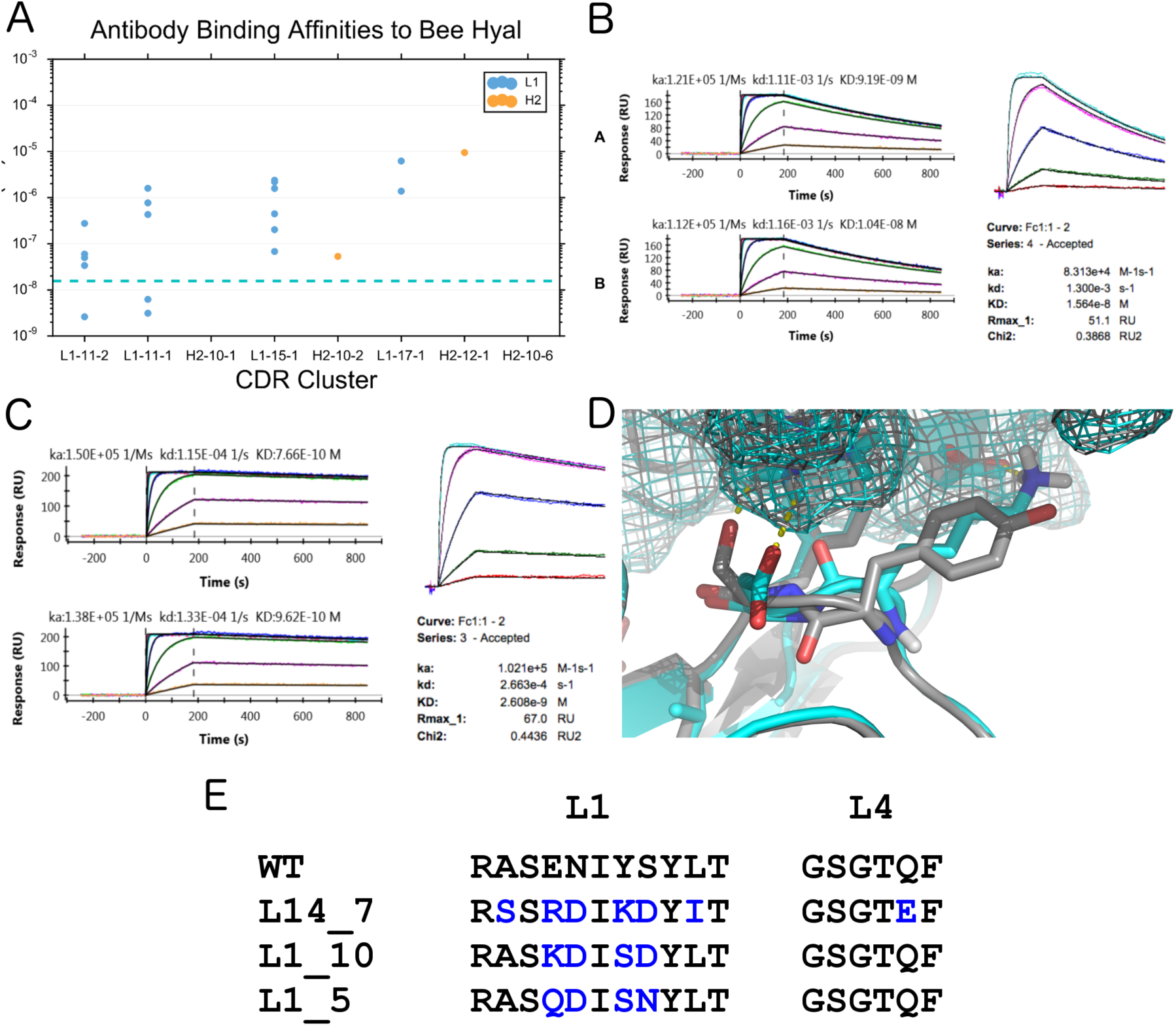
Designed antibodies against bee hyaluronidase. (A) Apparent Binding Affinity (*K*_D_) of expressed and tested antibody designs for bee hyaluronidase (PDB: 2J88), grouped by designed CDR cluster, as determined using Surface Plasmon Resonance (SPR) on a Biacore 4000. The dotted blue line represents the binding affinity of the native antibody on the Biacore machine (1.57×10^−8^ nM). Binding affinity is shown for the 26 designs that had detectable binding affinity (out of 30 tested). The native CDRs are L1-11-2 and H2-9-1. (B) Kinetic sensorgrams of WT 2J88 Antibody to Bee Hyaluronidase. Two repeats of *XPR* (left); *Biacore 4000* (right). (C) Kinetic sensorgrams of design L14_7 to Bee Hyaluronidase Two repeats of *XPR* (left); *Biacore 4000* (right). (D) Model of the interface changes in design L14-7, with designed L1 cluster L1-11-2 (cyan), superimposed onto the WT antibody from PDB ID 2J88 (gray). **(E)** Designed Sequences vs WT.

Three of these designs had improved binding affinity over the wild-type (WT), which binds at 9.2 nM (Fig. 8B), with the best design exhibiting a 12-fold improvement over wild-type with a K_D_ of 770 pM, as determined by Surface Plasmon Resonance (SPR) on a ProteOn XPR (Fig. 8C). This design, designated as L14_7, had a different L1 cluster (L1-11-2) than the wild-type (L1-11-1), with six amino acid differences in the L1 sequence, and a single amino acid difference in the L4 loop (Fig. 8D, 8E), which makes important contacts with the new L1 loop in the design model. Of the L1/L4 design group where docking was enabled, L14_7 had the lowest computational ΔG after filtering out the worst 90% of the designs by total Rosetta energy. The other two designs with better affinity than the native, L1_10 (Fig. F in S1 Supporting Information) and L1_5 (Fig. G in S1 Supporting Information) contained the same cluster as the native, but with 4 amino acid changes in L1 out of 11 positions. Kinetic studies of these designs and WT were done on both a Biacore 4000 and an XPR for a total of 3 replicates. Binding affinity was improved against WT for each of these designs in each replicate.

Thermostability (Tm) of WT and these designs were similar, as measured by Differential Scanning Calorimetry (DSC). While both the WT and designs had two Tm peaks, the major peak of the WT 2J88 antibody was measured at 75.0 °C, while the designs L14_7, L1_10, and L1_5 had similar Tm values of 71.9, 71.5, and 72.1 °C respectively (Fig. H in S1 Supporting Information).

Mutational analysis was done to delineate hotspot residues at the antibody-antigen interface. Rosetta was used to guide this manual analysis through the use of the PyRosetta Toolkit [85] and FoldIt Standalone [86] GUIs. Residue 7 (Lys) of the L1-11 loop of the L14_7 design was selected for mutation due to its proximity to the antigen (making hydrogen bonds to the antigen in the design model – Fig. 8D) and favorable Rosetta energy. This position was mutated back to its sequence-aligned WT residue (K38Y). Binding affinity worsened by approximately 3-4 fold as determined by SPR on a Biacore 4000 (Fig. I in S1 Supporting Information). The reverse experiment was done on the WT antibody for position 38 (Y38K) and the mutant exhibited improved binding (14.2 nM to 4.3 nM) as expected (Fig. J in S1 Supporting Information). Finally, as a proof-of-concept, we improved one of the weaker-binding designs (L1_4), which harbors a very long L1-17-1 loop by 2.5x, through a single S->V mutation at position 36 in the L1 loop (S37V, 655 nM -> 261 nM) (Fig. K in S1 Supporting Information).

We chose 27 design variants of the antibody CH103 to express and test for binding to HIV gp120 by using the AntibodyFeature reporters to rank and select prospective decoys. Of the 27 designs, 21 designs could be purified and tested for binding affinity against a panel of gp120 from different strains of HIV. Of these, 7 designs could bind one or more of the gp120 strains as determined through SPR on a Biacore 4000 (Fig. 9A and Table C in S1 Supporting Information). One of these antibodies, H2-6, improved binding affinity to most of the gp120s tested, with a 54-fold improvement to Core-Bal (91 nM to 1.7 nM) and a 40-fold improvement to HXB2 (52 nM to 1.32 nM) (Fig. 9B, 9C, Fig. L in S1 Supporting Information). H2-6 had the least number of buried unsatisfied hydrogen bonds in the interface in the H2 design group (Methods). This antibody design had a longer H2 loop (cluster H2-10-6) than the native (cluster H2-9-1), came from an unrelated mouse antibody [87] in the antibody design database, and is significantly different than the WT CDR (Fig. 9D, 9E).

**Fig. 9.**
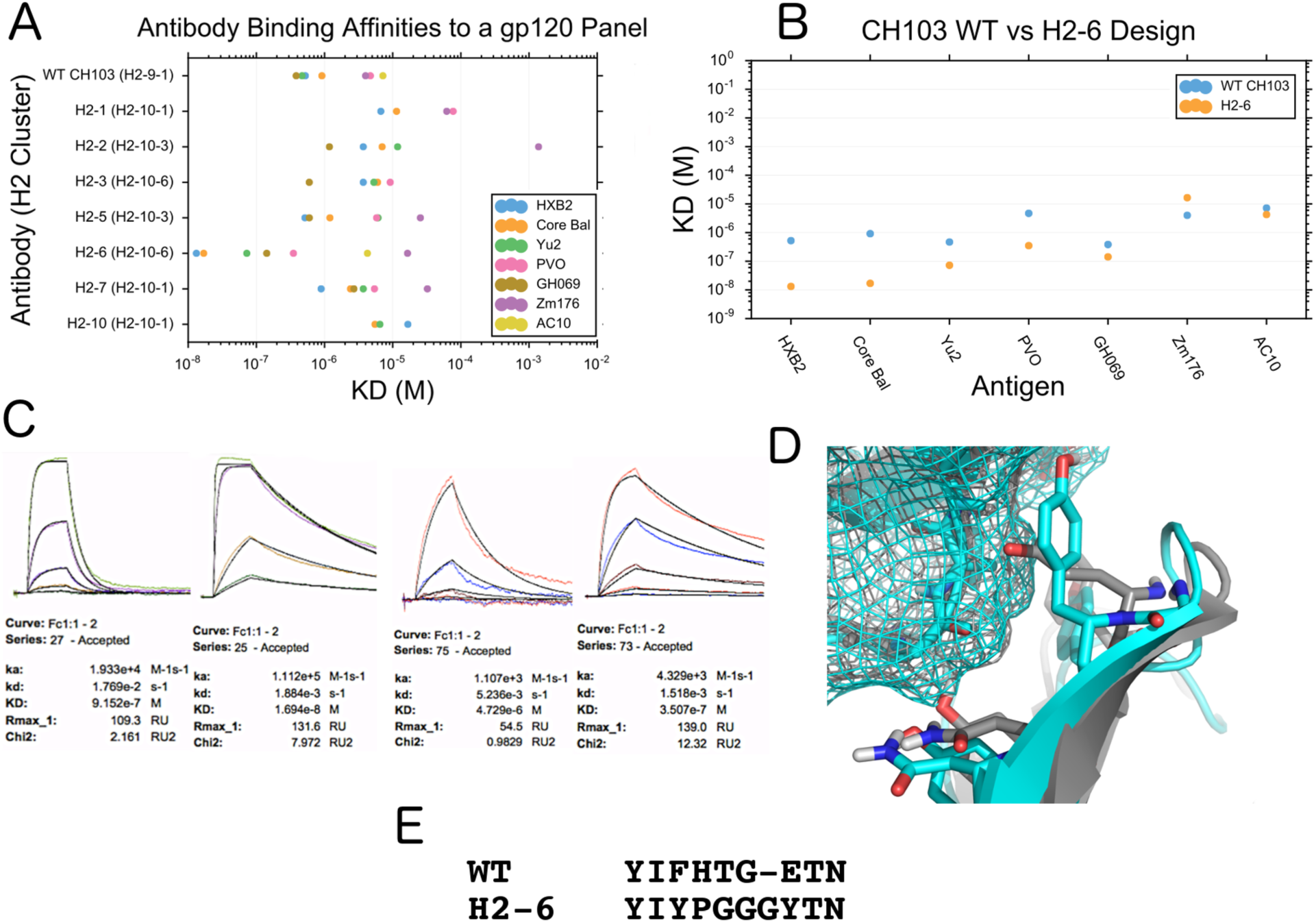
Binding of designed antibodies to HIV gp120. (A) Apparent binding affinity (KD) of WT CH103 antibody and designed antibodies to a panel of gp120 antigens. Here, 30 designs were expressed and tested, where 7 had detectable binding to these gp120s. (B) Binding affinity (KD) of the designed antibody, H2-6, versus the wild-type antibody CH103. (C) Kinetic sensorgrams of CH103 WT and design H2-6 to two select GP120s, Core Bal and PVO as determined through a Biacore 4000. (D) Model of the interface changes in design H2-6, with designed H2 cluster H2-10-1 (cyan), superimposed onto the WT antibody from PDB ID 4JAN (gray) (**E**) Alignment of H2-6 and the WT antibody CH103 from PDB ID 4JAN.

We performed mutational analysis on the CH103 designs and WT antibodies. Based on the sequence alignment of the H2 loops from the H2-6 design and the WT antibody (Fig. 9E) and structural observation using the Rosetta GUIs, we mutated two hypothesized hotspot residues within the H2 loop at positions 3 and 8 of the designed length-10 CDR in the H2-6 design (AHo numbering 59 and 67 respectively) to the aligned WT residue (Y->F and Y->E respectively). Binding affinity was measured for Core HXB2 and Core Bal using SPR on a ProteON XPR. Notably, the position 67 mutant decreased binding significantly (1.7 nM to 30.9 nM for HXB2; 8.8 nM to 212 nM for Core Bal), while the position 59 mutants had a smaller effect (1.7 nM to 2.2 nM for HXB2; 8.8 nM to 12.8 nM for Core Bal) (Fig. M in S1 Supporting Information). The reverse experiment was also done, where the H2-6 residues at the same positions were placed into the WT antibody. This reverse experiment confirmed position 67 as a hotspot residue (Fig. N in S1 Supporting Information); Kd for HXB2 improved 60-fold from 138 nM to 2.3 nM, and Kd for Core Bal improved 93 fold from 1.1 μM to 11.7 nM.

To investigate the role of glycans in CH103 binding, we created glycan knockouts in both the native antibody (Fig. O in S1 Supporting Information) and the ZM176 strain gp120 (Fig. P in S1 Supporting Information). Native antibodies do not usually have N-linked glycosylation sites near the paratope and in all cases except AC10, the glycan-knockout antibodies did not affect binding affinity significantly. However, multiple antibody designs were sensitive to the 463 and/or 386 glycans of gp120, which are in structural proximity to the antibody binding site. A single antibody design that included L1 and L3 CDRs design together was able to bind, but only with the potential 386 glycan knocked out. Meanwhile, two designs bound significantly better to ZM176 when the 386 glycan was knocked out. These glycan knockout studies show the importance of glycan considerations for some antigens and for antibody-design in general.

The results shown here demonstrate that RAbD can be used to successfully improve the binding affinity of antibodies, and that those designs can have different CDR lengths and clusters from the starting antibody.

## Discussion

The knowledge-based RosettaAntibodyDesign framework and application was developed to enable reliable, customizable structure-based antibody design for a wide variety of design goals and strategies based on a comprehensive clustering of antibody CDR structures [53]. To test the ability of RAbD to produce native-like antibody designs before it was used experimentally, we performed rigorous computational benchmarking using novel recovery metrics, the design risk ratio (DRR) and the antigen risk ratio (ARR), which provided needed statistical significance for recovery metrics over random sampling. The results showed that RAbD was able to enrich for native lengths and clusters – even with the large structural diversity of our underlying antibody design database and flat sampling over CDR clusters, while recovering native-like physical characteristics of the interface and antibody. We applied RAbD to two different antigen-antibody systems where the ability to tailor the program to a specific need and the use of our knowledge-based approach to both antibody design and selection led to successful experimental designs that improve binding affinity significantly using different CDR lengths and/or clusters.

While RAbD is highly tailorable, there are only a few choices that must be made for any particular antibody design project. First, after examining the initial structure of the antigen-antibody complex, the user must choose which CDRs to design and whether these CDRs should be subject to graft-design or only sequence-design. It may be the case that one CDR does not contact the antigen at all in the starting structure, and a user may choose to subject only that CDR to graft-design. The other CDRs may or may not require sequence design as well. In other cases, more drastic changes in the starting antibody may be desirable. For example in ab initio design to a new epitope or in redesigning an existing antibody for a homologue of its antigen, the user may choose to perform graft design on multiple CDRs. The user should also decide whether to optimize interface energy (opt-dG), total energy (opt-E) during the Monte Carlo design simulations, or a weighted combination of both,. If the existing antibody has low affinity, then interface dG may be the more relevant choice; however, if the existing antibody has low stability but reasonable affinity, then total energy may be more suitable.

Second, the user may select different optimization protocols for the inner loop of RAbD. This includes whether to perform docking refinements or not and whether to use more computationally intensive relax algorithms. For design against a native antigen present in the starting structure, we recommend not using the additional docking step, since local optimization will usually be sufficient for this purpose. It will usually be better to generate more decoy designs rather than expending CPU time on docking. However, if the antigen is not the same as the one for the starting antibody, either because it is an ab initio design or because it is homologous to the starting antigen, then we recommend including the docking step in the inner loop.

Third, the user can determine the number of inner and outer loop steps and the number of individual design runs to perform. The default values for the inner and outer loops steps are reasonable and usually do not need to be altered. The number of design runs should be at least 1,000 and may be as high as 10,000, depending on CPU availability for the final production run (100 was used for benchmarking purposes).

Finally, significant user input is needed in deciding how many and which antibodies to synthesize and test experimentally. Our rates of success – the number of successfully improved binders out of the number of antibodies expressed and tested in binding studies, were 3 in 30 for the bee venom antibodies and 1 in 27 for the HIV gp120 antibody redesign, which successfully bound better to gp120 from several strains of HIV. These are comparable to applications in other systems in the literature [70,88,89]. Our experience and that of others [90] acts as a guide for employing computational design techniques in real-world applications of computational interface design.

We recommend that users consider both the total energy of the antibody-antigen complex as well as the interface dG of the complex (regardless of which is used to govern the Monte Carlo decision steps). These values are reported in every output decoy file and the associated score file. In our experimental tests, we selected designs to synthesize that had low values of both total energy and interface dG. Other important features may include shape complementarity of the antibody and antigen and the number of hydrogen bonds and unsaturated hydrogen bonds within the interface. The choice of criterion should be based on the stability and affinity of the starting antibody and the goals of the design project.

RAbD is most similar to methods previously developed by two other research groups: OptCDR [41] and OptMAVEn [42], developed by Maranas et al., and AbDesign developed by Fleishman et al. [47] These authors also present computational benchmarking of their methods, and our benchmarking procedures, metrics, and results can be compared with theirs. We believe our benchmarks are better suited to testing computational antibody design methods than the work of previous authors, and that the risk ratios we have used provided needed context and statistical significance missing from earlier studies.

For OptCDR, Pantazas and Maranas constructed 100 decoys for 254 antibodies with CDRs borrowed from other antibody structures and used a simple scoring system that penalized steric conflicts of CDR backbone atoms with any atom of the antigen, favored interactions with a flat score between the sum of van der Waals radii and 8 Å between CDR backbone atoms and atoms of the antigen, and a zero score for longer distances. The native CDR coordinates had better scores than the constructed decoys on average. This is not surprising since almost all of the decoys would have at least one non-native CDR length or cluster, and the antibody as a whole would score worse than the exact native structure, which would have zero clash score and a favorable contact profile. They did not evaluate whether decoys with similar CDR lengths or clusters as the native scored well, as we have done with the length and cluster design risk ratios.

In a second computational experiment, they tested a set of 95 experimentally characterized mutants of a single antibody (anti-VLA1, PDB: 1MHP), 12% of which had improved affinity experimentally [91]. They claim 78% binary total accuracy (Q2) on this set, and a 50% positive predictive value (PPV), which is the percentage of their positive predictions that are true positives (improved affinities). It is impossible to discern a consistent and complete set of evaluative measures typically used in binary prediction methods (TPR, TNR, PPV, and NPV, balanced accuracy, etc.) [92] from these limited pieces of data. Finally, they performed a sequence design test and found that the native sequence scored better than all decoy sequences for two thirds of 38 test cases, although this does not indicate that the method could sample and find these native sequences from scratch, which is the typical sequence recovery metric in protein design. They did not provide any measures of statistical significance of these results, as we have done. By contrast, we measured sequence recovery of residues in contact with the antigen and achieved a 72% recovery in the presence of the antigen and 48% in simulations without the antigen, which is an ARR of 1.50±0.01 (95% confidence interval).

OptMAVEn [42] is based on the MAPS database developed by the same authors. For the heavy chain, κ light chain, and λ light chain, MAPS contains separate PDB files for the structures of V regions (through the beginning of CDR3), CDR3 segments, and post-CDR3 segments (“J regions”). If we count unique sequences: for the variable regions, there are 60 heavy, 34 κ, and 21 λ segments in MAPS; for CDR3, there are 413 heavy, 199 κ, and 39 λ sequences; and for the J regions, there are 3 heavy, 4 κ, and 6 λ sequences. RAbD uses an updated and updateable database of 754 non-redundant sequences per CDR (on average over 6 CDRs) to graft CDRs in any combination onto any starting framework, rather than spending CPU on designing the whole antibody variable domains, which may have already been optimized for stability or other properties. RAbD can mix CDR1s and CDR2s from different sources, rather than restricting them to a given V region from the PDB, as OptMAVEn does. Generally, this is a positive feature, since it allows RAbD to sample sequences and structures that are not likely to be present in an animal immune system or in an antibody display library. RAbD also has the ability to keep CDR sampling within a particular germline, including that of the starting antibody.

In their computational testing of OptMAVEn, Li et al. demonstrated that their grid search over antigen positions and orientations is able to sample (but not rank or score) structures relatively similar to the native structure for 120 antigen-antibody complexes (antigen protein lengths of 4 to 148 amino acids) [42]. This is useful to know but does not represent a recovery metric. They utilized OptMAVEn to produce designs for the same benchmark set and were able to produce designed antibodies with lower calculated interaction energies than the native for 42% of the cases, but this does not show that such antibodies would in fact bind better than the native, nor does it show that the native CDR lengths or conformations or sequences were obtained more frequently than one would expect, as our DRR does. For two antibodies, the authors were able to show that they could recover 20% (HIV VRC01) and 35% (Influenza CH65) of mutations from low-affinity antibodies to high-affinity, matured antibodies.

For their AbDesign method, Lapidoth et al. clustered the V regions of antibodies (up to the beginning of CDR3 of each variable domain) purely by the combination of lengths present at CDR1 and CDR2 [47]. They clustered CDR3 for each domain by length and RMSD. The input data consisted of 788 heavy-chain domains and 785 κ light-chain domains (no λ chains were included), broadly clustered into 5 κ V-regions, 2 L3 conformations, 9 heavy-chain V regions, and 50 H3 conformations. By contrast, RAbD uses the 72 CDR clusters of North et al. to group the non-H3 CDRs and contains 985 unique H3 sequences. Like OptMAVEn, AbDesign combines fragments that comprise the entire variable domains, rather than concentrating CPU on the design of the CDRs that contact the antigen. Thus it is not suitable for many design projects, which usually involve changing the sequences of one or more CDRs rather than a wholesale design of a new antibody, including the frameworks.

AbDesign was computationally tested on only 9 antibodies [47]. The authors compared features such as shape complementarity, buried surface areas, and interaction scores with the native antibodies. The average shape complementarity of their decoys was approximately 0.61, while that of the natives was 0.68. Our opt-dG decoys reached an average of 0.68 in shape complementary scores while the native structures in our benchmark averaged 0.70 (Fig. D in S1 Supporting Information). AbDesign’s 9 designed antibodies achieved an average of −26.1 REU in binding energy, while our 60 designed antibodies averaged −42 REU in the opt-dG simulations (Fig. B in S1 Supporting Information). In terms of recovery of the native structure, they calculated Cα RMSDs to native for each of the CDRs of the top scoring design for each of the 9 antibodies. For all of the non-H3 CDRs and 6 of the H3s, the designs had CDR lengths that matched the native antibodies. For 36 of the 45 non-H3 CDRs (80%), the Cα RMSDs were better than 1.0 Å.

It is difficult to assess the significance of these results, because the source database for sampling in AbDesign must be dominated by the same CDR lengths and clusters found in the native antibodies in the benchmark. To investigate this, we searched PyIgClassify for the 9 antibodies in this benchmark for their clusters according to our nomenclature. Since H3 does not cluster well beyond length 8, we report only the lengths of H3. The representation of clusters in their benchmark is as follows, including the number out of 9 antibodies in their benchmark set: L1-11-1 (4/9); L1-11-2 (3/9); L1-12-1 (1/9); L1-16-1 (1/9); L2-8-1 (9/9); L3-9-cis7-1 (7/9); L3-9-cis7-2 (1/9); L3-9-1 (1/9); H1-13-1 (8/9); H1-13-3 (1/9); H2-10-1 (7/9); H2-10-3 (1/9); H2-9-1 (1/9); H3-9 (1/9); H3-10 (4/9); H3-11 (2/9); H3-12 (2/9). L1-11-1 and L1-11-2 are very similar to each other (<0.5 Å RMSD). As it turns out, the 4 non-H3 CDRs with the largest RMSDs to native (>1.8 Å) are those with less common clusters or lengths: L1-12-1 (1IQD, 1.85 Å RMSD), L3-9-1 (1IQD, 2.12 Å RMSD), H1-13-3 (2CMR, 2.04 Å RMSD), and H2-10-3 (1P2C, 1.90 Å RMSD). For comparison, the very common H1-13-1 cluster is about 1.6 Å from H1-13-3, and the common H2-10-1 is 0.7 Å away from H2-10-3. This indicates that AbDesign is dominated by its sampling database in a way that makes the benchmarking data difficult to interpret. The design risk ratio we developed in this work solves this problem by demonstrating the increase in recovery over the sampling rate of any particular conformation in the database. Similarly, the antigen risk ratio demonstrates that the sampling and scoring is able to choose native-like CDRs when the antigen is present in the simulations more frequently than when it is absent, indicating that the design process is choosing CDR structures and sequences likely to bind the antigen. Finally, Lapidoth et al. achieved a sequence identity of 32% for residues in the binding site of the antibodies in their benchmark, compared to RAbD’s values of 72% in our opt-E benchmark (and a risk ratio of 1.50 over simulations in the absence of the antigen).

RAbD and AbDesign have a number of similarities and several important differences. They both utilize structural clusters of fragments of antibody structure and their associated sequence profiles to build new antibodies during a design simulation. They both utilize Rosetta’s docking and side-chain repacking routines to optimize the structure of the antigen-antibody complex during design.

The clustering of antibody structures and PSSM derivation differ substantially between the two methods. AbDesign breaks up each domain of antibodies into two segments – the V region up to the beginning of CDR3 and a segment containing CDR3 and the rest of the variable domain up to its C-terminus. AbDesign clusters its V regions only by the combination of sequence lengths of CDR1 and CDR2. Thus it samples the entire V domain and merges sequence data from different canonical structures of CDR1 and CDR2. This can lead to problems because many CDR clusters have required residue types, often glycine or proline, at certain positions in order to form the correct loop conformation. The strategy of replacing the entire heavy and light-chain variable domains with different fragments means that AbDesign is not suitable for optimizing existing antibodies, which is a very common task in antibody engineering and therapeutic development projects. Instead, as intended, it is more suitable for ab initio design of antibodies, which is a very challenging task.

Conversely, RAbD treats each CDR separately and samples structures from canonical clusters and their individual sequence profiles as defined in our PyIgClassify database, which is updated on a monthly basis. This allows mixing and matching of CDR1 and CDR2, while our PSSMs are more closely defined by the structural requirements of each canonical conformation. The CDRs are grafted onto the antibody framework provided by the user, which may have already been optimized for specific properties, rather than redesigning the entire variable domains, as AbDesign does.

RAbD and AbDesign are implemented in quite different ways. AbDesign depends on a series of Rosetta scripts, which are xml files that control Rosetta functions. It depends on a mover called splice, which is not documented. It has only been benchmarked on the score12 scoring function of Rosetta, which has not been the default scoring function since 2013. Finally, AbDesign is difficult to customize for specific problems in antibody design, such as sampling defined lengths of a given CDR or sampling from within a particular germline or CDR cluster.

RAbD is a full-fledged Rosetta application, a command-line program that runs the simulation according to command line options and rules provided in an optional CDR Instruction File. The run can be setup as simple as:

`antibody_designer.macosclangrelease -s 2r0l_1.pdb -graft_design_cdrs L1 -seq_design_cdrs L1 L2 L3 -light_chain kappa -nstruct 100`

The Instruction File makes RAbD highly tailorable. One or more CDRs can be designed or not designed and sampling of CDR structures for grafting can be restricted by length, cluster, species, germline, etc. All of RAbD’s dependencies are available in the public release of Rosetta. RAbD is also very well documented so that new users can quickly set up their design runs. RAbD has been benchmarked on the current Rosetta energy function, REF2015 [93], which utilizes our smoothed backbone-dependent rotamer library for protein side chains [94], our smoothed Ramachandran probability densities, and cubic splines for all ϕ,ψ-dependent scoring functions, as described by Leaver-Fay et al. [73]. These scoring functions are important for locally minimizing the Rosetta scoring function by altering backbone and side-chain dihedral angles. The older scoring function used by AbDesign contained very rough surfaces and linear spline estimates for the Ramachandran terms that resulted in poor structure optimization.

A major challenge moving forward, especially in regard to true de novo design, is the difficulty in effective sampling and design of the H3 loop, owing to its extreme variability and lack of clustering. To aid in this, H3-specific design strategies in the program can include up-weighting H3 graft sampling from other CDRs, restricting the use of H3 loops in the CDRSet to kinked-only (See Methods) [95,96], and reducing the search space to only short loop lengths that cluster well; however, much more work is needed to benchmark and test H3-specific design. Recently, we have shown that H3-like loops can be found in non-antibody proteins [95]. These loop structures could be used to supplement existing H3 structures in our antibody database as additional templates for H3 *GraftDesign*. The specific design of antibody H3 loops will be a major challenge in the next phase of antibody design methodology development, but using the RAbD framework and new methods for antibody design benchmarking outlined here should aid in this challenge.

These promising computational and experimental results show that RAbD is able to design antibodies with similar features to the native antibodies and antibodies with improved affinity. It can be used for a variety of antibody design tasks through the use of its highly customizable interface. RAbD represents a generalized framework and program for antibody design and makes many antibody design projects feasible that are either difficult or prohibitive using historical, traditional means, making computational antibody design a tangible reality.

## Methods

### Transfection and Expression

All antibody designs were expressed as IgGs in 293F cells using the pFUSE vectors (pFUSEss-CHIg-hG1 (human heavy), pFUSEss-CLIg-hL2 (human lambda), and pFUSEss-CLIg-hk (human kappa)). Bee Hyaluronidase and gp120 antigens were expressed in 293F cells using pHLsec vectors. Opti-Mem media and FreeStyle293 Expression media were first warmed to 37 °C. 293F cells were checked for viability at 95% and at a concentration greater than or equal to 2.4×10^6^ cells/ml. 6 mls of OptiMem were mixed with 125 μg of heavy chain DNA, and 125 μg of light chain DNA in one 15 ml conical tube, and 250 μg of fectin in the other. After a five minute incubation, the DNA tube was poured into the fectin tube and was left to incubate for 21 minutes. The 293F cells were then diluted to 1.2×10^6^ cells/ml, added to a 500 ml shaker flask, and the fectin/DNA mixture was added to the flask. The 500 ml flask was incubated in a 37 °C, 8.0% CO_2_, 80% humidity shaking incubator for four days. The supernatant was harvested on the fifth day using 500 ml centrifuge tubes and spinning for 20 minutes at 4000 rpm. The supernatant was then filtered in a 500 ml filter unit.

### Purification

Antibodies were purified by first using 1 ml of GE rProtein A Sepharose Fast Flow resin in a chromatography column, and then washing with 10 ml dH_2_O and 10 ml PBS. The antibody supernatant was poured onto the column and then washed with 10 ml of PDB, followed by 10 ml of 0.5 M NaCl in PDB, and then another 10 ml of PBS after all supernatant had passed through the column. The antibody was then eluted with 6 ml of Thermo Scientific IgG Elution Buffer into a 50 ml conical tube of 0.5 ml, 1M Tris-HCl. The eluted antibody was then placed into a Slide-ALyzer cassette and dialyzed against PBS with three changes. After dialysis, the antibody solution was filtered using a 0.22 micron syringe filer and the OD was checked to obtain the final concentration of antibody.

### Binding assays

Kinetics and affinity of antibody-antigen interactions were determined on a Biacore 4000 (GE Healthcare) using Series S Sensor Chip CM5 (BR-1005-30, GE Healthcare) and 1x HBSEP+ pH 7.4 running buffer (20x stock from Teknova, Cat. No H8022) supplemented with BSA at 1 mg/ml. We followed Human Antibody Capture Kit instructions (Cat. No BR-1008-39 from GE Healthcare) to prepare chip surface for ligands capture. In a typical experiment about 9000 RU of capture antibody was amine-coupled in appropriate flow cells of CM5 Chip. 3M Magnesium Chloride was used as a regeneration solution with 180 seconds contact time and injected once per each cycle. Raw sensorgrams were analyzed using Evaluation software (GE Healthcare), double referencing, Equilibrium or Kinetic with Langmuir model or both where applicable. Analyte concentrations were measured on NanoDrop 2000c Spectrophotometer using Absorption signal at 280 nm.

Antibody-Antigen binding kinetics were confirmed on a ProteOn XPR36 (Bio-Rad) using GLC Sensor Chip (Bio-Rad) and 1x HBS-EP+ pH 7.4 running buffer (20x stock from Teknova, Cat. No H8022) supplemented with BSA at 1mg/ml. We followed Human Antibody Capture Kit instructions (Cat. No BR-1008-39 from GE) to prepare chip surface for ligand capture. In a typical experiment, about 6000 RU of capture antibody was amine-coupled in all 6 flow cells of GLC Chip. 3M Magnesium Chloride was our regeneration solution with 180 seconds contact time and injected four times per each cycle. Raw sensorgrams were analyzed using ProteOn Manager software (Bio-Rad), interspot and column double referencing, Equilibrium or Kinetic with Langmuir model or both where applicable. Analyte concentrations were measured on NanoDrop 2000c Spectrophotometer using Absorption signal at 280 nm.

### Thermostability assays

Differential scanning calorimetry (DSC) experiments were performed on a MicroCal VP-Capillary differential scanning calorimeter (Malvern Instruments). The HEPES buffered saline (HBS) buffer was used for baseline scans and the protein samples were diluted into HBS buffer to adjust to 0.6 mg/ml. The system was set to equilibrate at 20 °C for 15 min and then heat up until a temperature of 125 °C was reached at a scan rate of 90 °C/h. Buffer correction, normalization, and baseline subtraction were applied during data analysis using Origin 7.0 software. The non-two-state model was used for data fitting.

### The RosettaAntibodyDesign Program

The RAbD protocol consists of repeated execution of an outer loop and an inner loop (Fig. 1). Most Rosetta protocols utilize Monte Carlo + minimization algorithms to optimize sequence and structure effectively. By allowing occasional increases in energy, we enable structures to overcome energy barriers to escape local energy wells in order to drive the energy down further. In order to traverse the energy landscape more effectively, the Monte Carlo criterion is applied during the design simulation for both the outer and inner loops of the algorithm.

The general RosettaAntibodyDesign protocol consists of 4 major tasks:

1. *Choosing a CDR structure to graft:* In the other loop, randomly choosing a CDR from those CDRs set to design, choosing a CDR cluster for that CDR, and choosing a structure from the design database for that CDR cluster (Fig. 1A).
2. *Grafting a CDR:* Grafting that CDR onto the antibody framework. This structure is then passed to *N_inner_* cycles of the inner loop.
3. *Sequence design and side-chain repacking in the inner loop:* Sequence design in Rosetta consists of a Monte Carlo side-chain repacking procedure in which residues to be designed sample rotamers of multiple residue types. All residues in the grafted CDR passed from the outer loop to the inner loop can be redesigned in one round of the inner loop.
4. *Local energy minimization and application of the inner-loop Monte Carlo criterion*. After sequence design and repacking, local steepest-descent energy minimization (or optionally Rosetta’s Relax algorithm) is applied, which alters the dihedrals of the backbone and the side chains. The inner cycle Metropolis Monte Carlo criterion is then applied to the resulting structure using either the total energy (opt-E) or the interface energy (opt-dG) after each cycle of the inner loop.
5. *Application of the outer-loop Monte Carlo criterion:* Once a structure exits the inner loop after *N_inner_* cycles (default 1), the structure is then passed back to the outer loop where the Monte Carlo criterion is applied, and the algorithm continues with Step 1. The outer loop Metropolis Criterion can either be applied on the Total Energy (opt-E) or the Interface Energy (opt-dG). The cycle repeats (Step 1-5) for *N_outer_* cycles (default 25). The output design is the structure with the lowest energy observed during the simulation.

The entire procedure may be repeated many times (1,000-10,000) so that an ensemble of designs is produced from which some number of the top-ranking sequences may be chosen for synthesis and testing. The outer and inner loops of RAbD can be tailored for a variety of design projects and design strategies through the optional CDR Instruction File, an abundant set of command-line options, and object-oriented code design, which enables RosettaScript-able [97] framework components. Each of the five basic steps is described below in turn. Further details are provided in the Supplemental Methods.

#### (1) Choosing a CDR structure to graft

The CDRSet instructions tell the program which CDR lengths, clusters, and specific structures to include or exclude from the antibody design database for graft-based design. By default, every CDR length is enabled. The light chain type (κ or λ) can be specified on the command-line in order to limit the CDRSet to those that originate from that gene, which is aimed at increasing stability of the final antibody. No light chain is specified for camelid antibody design.

There are three simple algorithms that control how the CDR structure is chosen from the database during the *GraftDesign* stage. The default is to choose a CDR cluster from the list of available clusters and then choose a structure from that cluster (*even_cluster_mc*). One can also choose a structure from all available structures, which samples according to the prevalence of that length and cluster in the database (*gen_mc*). Or the outer loop can choose a length randomly and then a cluster given that length, and finally a structure from that cluster (*even_length_cluster_mc*). We recommend *even_cluster_mc* for most purposes. Finally, when designing a single CDR, the *deterministic_graft* algorithm can be used to graft every structure available. The lengths, clusters, and particular structures that are sampled and grafted can be controlled through the CDR Instruction File.

RosettaAntibodyDesign uses an SQLITE3 database to house all antibody and CDR data needed for the program, including full structural coordinates of CDRs, CDR length, cluster, species, and germline identifications, as well as CDR cluster sequence profile data. The publicly available release of Rosetta includes a smaller database (about 30 MBytes) that includes only the structures in the data analyzed by North et al in 2011. As with other large databases utilized by Rosetta [98], the current database is too large to be distributed with Rosetta by default. It can be obtained from PyIgClassify [54], which is typically updated every month and reflects data from the current PDB. All computational benchmarking in this paper utilized a recent version of the database (August 2017). The experimental tested designs utilized a version from November 2016.

The up-to-date database consists of only non-redundant CDR data at a 2.8 Å resolution and 0.3 R factor cutoff. CDR cluster outliers are then removed as described in the Supplemental Methods. In order to cull for non-redundancy in the remaining CDR loops, the CDR is selected in the order of: highest resolution à lowest R factor à lowest normalized distance to the cluster centroid. These databases are used in all aspects of the antibody design algorithm, including the *GraftDesign* step, which uses the raw coordinates in the database and the *SeqDesign* step, which uses an additional table for CDR cluster profiles (residue probabilities at each position) created from the non-redundant sequence data. The CDR Instruction File helps enable additional culling during the *GraftDesign* step, controlling which lengths, clusters, species, germlines, and structures, are used or left out.

The majority of H3 structures contain a “kink” at the C-terminus involving a Cα-Cα-Cα-Cα dihedral around 0° for the last three residues of H3 and the conserved tryptophan residue immediately following H3. More than 80% of H3 structures contain this kink, whose function in part is to break the β-sheet and allow the H3 CDR to form diverse non-β structures [95]. H3-specific control is available, such as limiting the H3 CDRSet to kinked-only structures. The kink option is useful if H3 is being sequence-designed, as some mutations in kinked H3s may make the H3 adopt an extended β-strand-like conformation and vice-versa. In addition, by default, we disable sequence design of the H3 stem region, which is known to influence the H3-kink [95,96].

The span of framework residues with AHo numbering 82-89 comprise what is commonly referred to as the “DE loop” (Chothia residues 66-71 in the light chain and 71-78 in the heavy chain). These variable residues form a loop physically in contact with CDR1 and are occasionally observed making contacts with antigen [53,84]. In RosettaAntibodyDesign, we denote the DE loop region as L4 and H4 for the light and heavy chains respectively. The typical kappa L4 is different in sequence and conformation than the typical lambda L4, so that lambda L4s should be used with lambda L1 CDRs and frameworks [84] and κ L4s should be used with kappa L1 CDRs and frameworks. Both loops can be considered CDRs in the application and can be specified just as any other CDR except for the *GraftDesign* stage. However, since the conformation of L4 and H4 is largely conserved among κ, λ, and heavy chain variable domains, and these loops do not usually contact the antigen, we typically do not set them to graft-design and in most cases do not set them to sequence-design either.

#### (2) Grafting a CDR

In order to sample whole structures of CDR conformations from our design database, a way to graft them onto a given antibody was needed that was quick and accurate enough to minimally perturb the CDR region without leaving breaks in the structure or non-realistic peptide bond lengths and angles. Our grafting algorithm (*CCDEndsGraftMover*) first superimposes three residues on either end of the CDR to be grafted onto the template framework. Those residues are then deleted and the Cyclic Coordinate Descent (CCD) algorithm of Canutescu and Dunbrack [71] is used to attach the CDR to the framework using two residues of the framework and the first residue of the CDR (on both sides of the CDR), while all other residues are held fixed. Each closure attempt first perturbs the backbone ϕ and n of these residues and the energy of these residues is minimized after the attempted closure.

A graft is considered closed if the peptide bond C-N distance is less than 1.5 Å and both the Cα-C-N and C-N-Cα angles are less then 15 degrees away from the ideal min and max values determined by Berkholz et al. [99] (114.5°, 119.5° and 120°, 126° respectively). If the graft is not closed after a specific number of cycles, we use the grafting algorithm from the older Anchored Design Protocol [81] followed by a minimization of the CDR and connecting residues with tight dihedral constraints on all residues. This protocol is generally much slower and can result in larger perturbations to the overall structure of the CDR loop relative to the framework, but can close most grafts due to the mobility of the entire insert region. When both terminal ends are closed during the protocol in either algorithm, we continue the design protocol.

Using this combined grafting algorithm, most CDR grafts can be completed in less than a second and we accomplish 100% of CDR graft closure while minimally perturbing the internal CDR structure, if at all. This algorithm is now also used for grafting within the main antibody application of RosettaAntibody [79], fixing many loop closure imperfections of the original application.

#### (3) Sequence Design and Side-chain Optimization

The *SeqDesign* options control which strategy to use when doing sequence design (primary strategy), and which strategy to use if the primary strategy cannot be used for that CDR (fallback strategy), such as conservative design or no design.

In Rosetta, the optimization of side-chain conformations is referred to as packing (or repacking). Packing in Rosetta consists of traversing the set of residues to be optimized randomly until no residues are left in the pool and selecting the best rotamer of all rotamers defined for the given residue [64]. Design is accomplished in the packing algorithm by sampling all rotamers (using the 2010 Dunbrack Rotamer Library [94]) of a specified set of residue types allowed at a given position.

In general, we use a probability distribution of residue types for each CDR position, embedded in the antibody design database. In the RAbD framework, it is how we sample from the profile of a given CDR cluster type. Each time packing is applied, a residue type for each position is chosen based on distributions from the antibody design database. If this residue is different from the starting residue, it is added to the design types. This process can be performed repeatedly to increase the sampling according to the distributions. This methodology helps to maintain the residue profile of a given CDR cluster. This is in use in the Antibody Design framework by default if enough statistics for that CDR cluster are present.

Alternatively, we use a set of conservative mutations as design types for each specified position. The conservative mutations for each residue type are composed of the substitutions for each residue which score ≥ 0 in one of the BLOSUM matrices [100]. All BLOSUM matrices can be used for conservative design, and are specified through the use of a command-line option. The numbers of the matrix (such as BLOSUM62) indicate the sequence similarity cutoffs used to derive the BLOSUM matrices, with higher numbers being a more conservative set of mutations. By default, the conservative mutations from the BLOSUM62 matrix are used to strike a balance between variability and conservation. This methodology is the default fallback sequence design strategy but can be used generally instead of profile-based sequence design.

By default we disable sequence design for prolines, cysteine residues involved in disulfide bonds, and the H3 kink-determining stem region (first 2 and last 3 residues of H3) [95,96] in order to limit large, unproductive perturbations of the CDR loops from disruptive sequence changes.

Users may further disallow amino acid types for *all* positions through a command-line option, for *specific CDRs* through the CDR instruction file, and for *specific positions* through the use of the Rosetta *resfile* format. The *resfile* can also be used to disable specific positions from design or packing altogether.

Within the protocol, both antibody-antigen interface residues and neighbor residues (Fig.2) that are computed for side-chain packing and design are updated on-the-fly before each packing/design step. This allows the algorithm to continually adapt to the changing environment and is accomplished through Rosetta’s graph-based neighbor detection algorithms.

#### (4) Local energy minimization and application of the inner-loop Monte Carlo criterion

Following sequence design via repacking (Step 3), the conformations of the grafted CDR, its neighboring CDRs, and nearby framework residues are optimized. The type of minimization and which CDRs are minimized as neighbors to other CDRs during the protocol can be specified through the *MinProtocol* section of the *Instruction File*. Many minimization types are implemented. The default is the standard lbfgs_armijo_nonmonotone minimizer in Rosetta with a tolerance of 0.001 REU. But other options include the backrub motion protocol [63,101], and the Relax algorithm, which includes alternating cycles of reducing and then ramping up the repulsive van der Waals energy term, and at each step performing side-chain repacking and local dihedral angle space energy minimization.

In order to achieve flexible-backbone design and environmental adaptation of the packing/design algorithm as described above, we updated the *FastRelax* algorithm [60] to enable sequence design during backbone and side-chain optimization. These changes to *FastRelax*, which we call *RelaxedDesign*, were used in the optimization step of Jacobs et al.to general success [102]. This is an optional alternative for the minimization step

One further option is in the inner loop is integrated sampling of the antibody-antigen orientation during design uses the underlying framework and docking algorithms of *RosettaDock* [45,103–106]. A ‘dock cycle’ consists of a low-resolution docking step, side-chain repacking of the interface residues (defined as residues of the antibody or antigen that are within the set interface distance of each other (8 Å default), minimization of the rigid body orientation (the *‘jump’* in Rosetta parlance) between the antigen and the antibody, and a shortened high-resolution dock consisting of 3 outer cycles and 10 inner cycles as opposed to 4 and 45, which is used for a full *RosettaDock* high-resolution run. During high-resolution docking, the current interface side chains are optimized, while any CDRs or specific residues set to design are designed. In this way, sequence design and antibody-antigen orientation optimization are coupled in the same vein as sequence design is accomplished during CDR structural optimization.

#### (5) Application of the outer-loop Monte Carlo criterion

Once a structure exits the inner loop after *N_inner_* cycles (default 1), the structure is then passed back to the outer loop where the Monte Carlo criterion is applied, comparing the structure that entered the inner loop and the structure that exited the inner loop. The outer loop Metropolis Criterion can either be applied on the Total Energy (opt-E) or the Interface Energy (opt-dG) or some weighted combination of the two. Once *N_outer_* cycles have been completed (Steps 1-5, default 25), the output design is the structure with the lowest energy observed during the simulation.

### Computational Benchmarking

In order to reduce artifacts, all benchmarking complexes and starting complexes for antibody design were first minimized into the Rosetta energy function using the Pareto-optimal protocol of Nivon et al. [62], which relaxes [60] the starting structure using restraints on the backbone and side-chain atoms to strike a balance between deviation from the starting structure and minimization of the energy. This Pareto-optimal method produces models with all-atom Root Mean Square Deviations (RMSD) to the starting structures at a mean of 0.176 Å [62]. For all starting structures, the lowest-energy model of ten decoys was used as the starting structure. An example command to run this protocol and the flags are given in the Supplemental Methods.

All antibodies were renumbered into the AHo numbering scheme [107] using PyIgClassify. The *CDRClusterFeature, InterfaceFeature*, and *AntibodyFeature* reporters (Table D, Table E and Table F in S1 Supporting Information) were used to determine CDR length and cluster information and physical characteristics of the decoys for benchmarking and design selection. In general, analysis was done using the Feature Reporters to analyze decoys and create databases with physical data, and the public, open-source Jade repository (https://github.com/SchiefLab/Jade) was used for benchmarking calculations and selections.

Antibody-protein complexes used for benchmarking consist of 46 κ and λ 14 structures. These complexes were chosen with the following criteria: 1) resolution ≤ 2.5 Å; 2) interface surface area of ≥ 700 Å2; 3) CDR1 and CDR2 within 40° of one of the cluster centroids in PyIgClassify (most L3 CDRs were also within 40°, Table A in S1 Supporting Information); 4) contacts with the antigen from both the heavy and light-chain CDRs with a preference for contacts of all 6 CDRs; 5) non-redundant such that no two antibodies in the benchmark contacted the same antigen with overlapping epitopes; 6) a diversity of CDR lengths and clusters.

For each input antibody, we ran a total of 100 simulations, and the best decoy observed during each design run was output. The benchmarking was run on a compute cluster in parallel via MPI. The RunRosettaMPIBenchmarks.py script of the Jade github repository was used to help launch and configure benchmarks on the cluster (https://github.com/SchiefLab/Jade). The number of outer cycles for each parallel run was set to 100, so each input antibody underwent 10,000 total design cycles for each experimental group.

All 5 non-H3 CDRs were allowed to undergo *GraftDesign*, while all 6 CDRs went through *SequenceDesign*. The starting CDR for each non-H3 CDR was removed at the start of the program and a random CDR from the CDRSet was grafted onto the starting antibody through the option –*random_start*.

To calculate the risk ratios over the entire benchmark, we calculated the percent recovered and the percent sampled over the 100 decoys for the 60 antibodies in the benchmark. Thus, the recovery frequency was the number of native clusters observed in the 6000 decoys of the benchmark for each CDR divided by 6000. Similarly, the sampled frequency was calculated as the number of native cluster CDRs grafted divided by the total number of grafts for a CDR during the 6000 simulations (for each CDR, (100 outer loops) x (1/6 CDRs) x 100 simulations x 60 antibodies = 100,000). For the antigen risk ratios, the frequencies of recovered CDR lengths and clusters were calculated for the final decoys from the antigen-present and antigen-free simulations.

The confidence intervals are calculated as described by Gertsman [108]. If *RR* = *p_Recovered_/p_Sampled_* than:

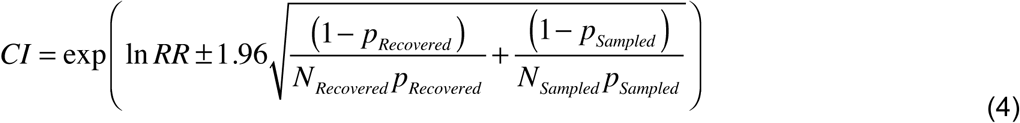

Similarly, for the antigen risk ratio, if *RR* = *p_antigen_/p_noantigen_*, where *p* represents the frequency of the native cluster, length, or residue type in the antigen or no-antigen simulations, then

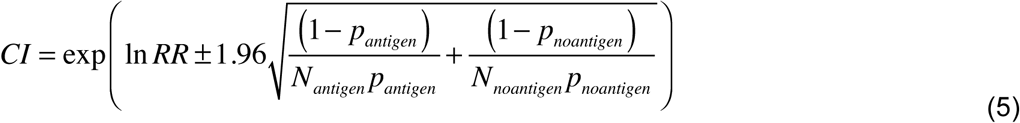

### Designs for experimental testing of computational antibody design

The starting antibody complex, was obtained from the Protein Data Bank with ID 2J88 [109], renumbered using PyIgClassify, and minimized into the Rosetta energy function as described above. To begin design, we used a multiple-strategy approach including with and without docking and the explicit use of only CDR clusters which have cluster profiles (more than 10 non-redundant members in the database), as well as differential CDR design for both graft-based and sequence-based design (H2 vs. L1). When designing the L1 loop, we also included a strategy in which we allowed L4 to undergo sequence design, for a total of 6 antibody design strategies. The WT L1 or H2 CDR was removed and a random CDR from the database was grafted in order to start design with the non-native CDR, as well as start with a potentially higher-energy structure. All CDR structures from the WT 2J88 antibody were left out of the CDRSet.

For the strategies in which docking was used, automatic epitope SiteConstraints were enabled to constrain the antibody paratope to the starting epitope. A total of 1000 top decoys were output as separate Monte Carlo trajectories in parallel for each design strategy using a compute cluster via MPI, with 100 outer cycles for each parallel run, for a total of 100,000 design rounds per strategy. The RunRosettaMPI Bio-Jade python application was used to aid the cluster run (https://bio-jade.readthedocs.io/en/latest/). The command to run the application, flags, and CDR Instructions are given in the Supplemental Methods.

Decoys were analyzed by the Rosetta Feature reporter framework in the exact same manner as the benchmarking. The feature databases were then used in the RAbD Jade Antibody Design GUI in order to sort them for selection (Fig. Q in S1 Supporting Information).

For both relaxed and unrelaxed sets of decoys and each antibody design strategy, we sorted the models according to their computed interface energy (dG) after culling to only the top 10% of the models by total energy (dG_top_p_total), or by the lowest density of unsaturated hydrogen bonds per interface area (delta_unsats_per_1000_dSASA) [90] for a total of 24 sorted groups (6 design strategies x 2 decoy discrimination methods (relaxed/unrelaxed) x 2 sorting methods).

For the three design strategies where docking was enabled and sorted by unsaturated hydrogen bond density (3 design strategies x 2 decoy discrimination methods x 1 sorting strategy), the best two models had antibodies that were too far from the native binding site, even with the use of epitope *SiteConstraints*. This could be due to not using the constraint energy as a filter in this case, only to guide the design –i.e., not in the sorting of the total energy. Due to this, these 6 groups were left out for a total of 18 groups. The best two models from the sorted, unrelaxed groups (9 groups, 18 designs) and the best model of the sorted, relaxed groups were expressed (9 groups, 9 designs) (i.e., chosen with no human intervention). Three other models of the sorted relaxed groups for L1 design were added to the expression group, as these were the second-best scoring models of the L1 relaxed groups, for a total of 30 antibodies selected for expression. Sequences were obtained from the decoys and processed for inclusion into the expression vector sequences using the *get_seq* application of Jade.

The starting antibody complex was obtained from the Protein Data Bank with ID 4JAN [82], renumbered using PyIgClassify, and minimized into the Rosetta energy function as described above. A total of four antibody design strategies were used where either H2 or L1+L3 were designed and the CDRSet included only clusters with enough data to use profiles.250 top decoys were output for parallel Monte Carlo trajectories in parallel for each antibody design strategy with the outer cycle rounds set to 200, for a total of 50, 000 design cycles per design strategy. Commands, flags, and CDR Instructions are given in the Supplemental Methods.

Decoys were analyzed with both the RosettaFeature reporters and physical data and sorted as described for 2J88. In addition to sorting by the top dG of the top 10% of total energy (*dG_top_Ptotal*) and density of unsaturated hydrogen bonds per interface area (*delta_unsats_per_1000_dSASA*), we sorted by the Lawrence and Colman Shape Complementarity score [110] (*sc_value*) through the Jade RAbD GUI. Sorts were done for both relaxed and nonrelaxed decoys to aid in decoy discrimination. The sorts were done for individual antibody design strategies and all combined for a total of 28 sorted groups. Jade was used to output PyMol sessions of each group. The top 10 designs from each group were visually analyzed in PyMol and 27 designs were selected based on physical characteristics such as good shape complementarity, hydrogen bonding, interface, and total energies, as well as cluster and sequence redundancy in the designs. Generally, the top design selected from each sort was expressed, unless it was redundant or the structure held some abnormality, such as bad shape complementarity. Sequences were obtained from the decoys and processed for inclusion into the expression vector sequences using the *get_seq* application of Jade.

### Availability

RosettaAntibodyDesign is distributed with the Rosetta Software Suite (www.rosettacommons.org) and is included with Rosetta versions starting at 3.8. All RosettaAntibodyDesign framework classes are available for scripting within the *RosettaScripts* framework [97], including the main application. The public Rosetta distribution includes a database of the original North-Lehmann-Dunbrack clustering data [53]. Up-to-date antibody design databases for use with RosettaAntibodyDesign can be obtained from PyIgClassify (http://dunbrack2.fccc.edu/pyigclassify). Documentation on the use of RosettaAntibodyDesign, including instructions for installing an up-to-date PyIgClassify database, can be found with the RosettaCommons documentation:

https://www.rosettacommons.org/docs/latest/application_documentation/antibody/RosettaAntibodyDesign.

Bio-Jade is an open-source python package, with scripts and modules created specifically for RAbD (https://bio-jade.readthedocs.io/en/latest/).

## Supplemental Methods

### 1. Rosetta Commands

For the benchmarking set, full commands and CDR Instruction File contents are given below. Note that some options are the default and do not need to be specified, but are given in the file explicitly anyway. To prepare the structures for design, they are relaxed with the Rosetta force field using the FastRelax protocol with these commands:

*Command*:

~~~
relax.mpi.linuxgccrelease –l PDBLIST.txt –nstruct 10
@**pareto_optimal_flags.txt**
~~~

*Contents of **pareto_optimal_flags.txt** file:*

~~~
  -no_optH false
  -flip_HNQ
  -use_input_sc
  -constrain_relax_to_start_coords
  -relax:ramp_constraints false
  -relax:coord_constrain_sidechains
  -ignore_unrecognized_res
  -ignore_zero_occupancy false
  -pdb_comments
  -ex1
  -ex2
  -out:pdb_gz
  -other_pose_to_scorefile
  -scorefile_format json
  -jd2:delete_old_poses
  -load_PDB_components false
~~~

The benchmark antibodies are designed with the following commands (with variations for each protocol):

*Command:*

~~~
Antibody_designer.mpi.linuxgccrelease @**common_flags.txt**
@**experimental_flags.txt** –l PDBLIST.txt
~~~

*Contents of **common_flags.txt** file:*

~~~
  -graft_design_cdrs L1 L2 L3 H1 H2
  -seq_design_cdrs L1 L2 L3 H1 H2 H3

  -output_ab_scheme AHo_Scheme
  -nstruct 100
  -outer_cycle_rounds 100
  -random_start True
  -add_graft_log_to_pdb
  -ignore_zero_occupancy false
  -ignore_unrecognized_res
  -pdb_comments
  -ex1
  -ex2
  -use_input_sc
  -out:pdb_gz
  -other_pose_to_scorefile
  -scorefile_format json
  -flip_HNQ
  -delete_old_poses
  -load_PDB_components false
~~~

*Contents of **experimental_flags.txt** file (in separate files for each example listed below as separated by the # sign):*

~~~
  #Lambda/opt-E
  -light_chain lambda

  #Lambda/opt-E/No antigen
  -light_chain lambda
  -remove_antigen True

  #Lambda/opt-dG
  -light_chain lambda
  - mc_optimize_dG

  #Lambda/opt-dG/No antigen
  -light_chain lambda
  -mc_optimize_dG
  -remove_antigen True

  #Kappa/opt-E
  -light_chain kappa

  #Kappa/opt-E/No antigen
  -light_chain kappa
  -remove_antigen True

  #Kappa/opt-dG
  -light_chain kappa
  - mc_optimize_dG

  #Kappa/opt-dG/No antigen
  -light_chain kappa
  - mc_optimize_dG
  -remove_antigen True
~~~

*Contents of CDR Instruction File:*

~~~
  #RAbD Defaults (Used always, unless command override):
  L1 MinProtocol Min_Neighbors L2 L3
  L2 MinProtocol Min_Neighbors L1
  L3 MinProtocol Min_Neighbors L1 H3
  H1 MinProtocol Min_Neighbors H2 H3
  H2 MinProtocol Min_Neighbors H1
  H3 MinProtocol Min_Neighbors L1 L3
  ALL MinProtocol MinType min

  #Experiments
  ALL CDRSet CLUSTER_CUTOFFS 5
~~~

For the redesign of the 2j88 antibody, the Rosetta command and files were:

*Command:*

~~~
  antibody_designer.mpi.linuxgccrelease @**common_flags.txt**
  @**experimental_flags.txt –cdr_instructions cdr_instructions.txt** –s
  2j88_pareto_optimal.pdb –nstruct 1000
~~~

*Contents of **common_flags.txt** file:*

~~~
  -output_ab_scheme AHo_Scheme
  -add_graft_log_to_pdb
  -ignore_zero_occupancy false
  -ignore_unrecognized_res
  -pdb_comments
  -ex1
  -ex2
  -use_input_sc
  -out:pdb_gz
  -other_pose_to_scorefile
  -scorefile_format json
  -flip_HNQ
  -delete_old_poses
  -load_PDB_components false
~~~

*Contents of **experimental_flags.txt** file, Docking Off:*

~~~
  -outer_cycle_rounds 100
  -s input_pdbs/pareto_2j88_renum_0002.pdb.gz
  -run_relax
  -random_start
  -design_protocol even_cluster_mc
  -light_chain kappa
~~~

*Contents of **experimental_flags.txt**, Docking On:*

~~~
  -outer_cycle_rounds 100
  -run_relax
  -random_start
  -design_protocol even_cluster_mc
  -light_chain kappa
  -use_epitope_constraints
  -do_dock
  -inner_cycle_rounds 2
~~~

*Contents of **cdr_instructions.txt**, H2 Design:*

~~~
  H2 ALLOW
  ALL GraftDesign mintype relax
  H2 MinProtocol Min_Neighbors H1 H3
  H2 CDRSet Cluster_Cutoffs 10
  ALL CDRSET EXCLUDE PDBIDs 2J88
~~~

*Contents of **cdr_instructions.txt**, L1 Design:*

~~~
  L1 ALLOW
  ALL GraftDesign mintype relax
  L1 GraftDesign Min_Neighbors L3 L4
  L1 CDRSet Cluster_Cutoffs 10
  ALL CDRSET EXCLUDE PDBIDs 2J88
~~~

*Contents of **cdr_instructions.txt**, L1/L4 Design:*

~~~
  L1 ALLOW
  ALL GraftDesign mintype relax
  L1 GraftDesign Min_Neighbors L3 L4
  L1 CDRSet Cluster_Cutoffs 10
  ALL CDRSET EXCLUDE PDBIDs 2J88
  L4 SeqDesign ALLOW
~~~

Note that many options, such as which CDRs to design, can alternatively be set via simple command-line options.

For the redesign of the 4JAN antibody, the Rosetta command and files were:

*General Command:*

~~~
  *antibody_designer.mpi.linuxgccrelease @**common_flags.txt –***
  *cdr_instructions cdr_instructions.txt –s 4JAN_pareto_optimal.pdb –*
  *nstruct 250*
~~~

*Contents of **common_flags.txt** file:*

~~~
  -outer_cycle_rounds 200
  -run_relax
  -random_start
  -design_protocol even_cluster_mc
  -light_chain lambda
  -add_graft_log_to_pdb
  -ignore_zero_occupancy false
  -ignore_unrecognized_res
  -pdb_comments
  -ex1
  -ex2
  -use_input_sc
  -out:pdb_gz
  -other_pose_to_scorefile
  -scorefile_format json
  -flip_HNQ
  -delete_old_poses
  -load_PDB_components false
~~~

*Contents of **cdr_instructions.txt**, H2 Design, only clusters with profiles:*

~~~
  H2 ALLOW
  H2 MinProtocol Min_Neighbors H1 H3
  H2 CDRSet Cluster_Cutoffs 10
  ALL CDRSet Exclude PDBIDs 4JAN
~~~

*Contents of **cdr_instructions.txt**, H2 Design, all:*

~~~
  H2 ALLOW
  H2 MinProtocol Min_Neighbors H1 H3
  ALL CDRSet Exclude PDBIDs 4JAN
~~~

*Contents of **cdr_instructions.txt**, L1/L3 Design, only clusters with profiles:*

~~~
  L1 ALLOW
  L3 ALLOW
  L3 MinProtocol Min_Neighbors L1 H3
  L1 MinProtocol Min_Neighbors L3 H3
  L1 CDRSet Cluster_Cutoffs 10
  L3 CDRSet Cluster_Cutoffs 10
  ALL CDRSet Exclude PDBIDs 4JAN
~~~

*Contents of **cdr_instructions.txt**, L1/L3 Design, all:*

~~~
  L1 ALLOW
  L3 ALLOW
  L3 MinProtocol Min_Neighbors L1 H3
  L1 MinProtocol Min_Neighbors L3 H3
  ALL CDRSet Exclude PDBIDs 4JAN
~~~

The command for the AntibodyFeature reporters (described below) used in the analysis of the antibodies was:

*Command:*

~~~
rosetta_scripts.macosxrelease –l DECOYS.txt @common_flags –parser:protocol
features_script.xml
~~~

*Contents of features_script.xml:*

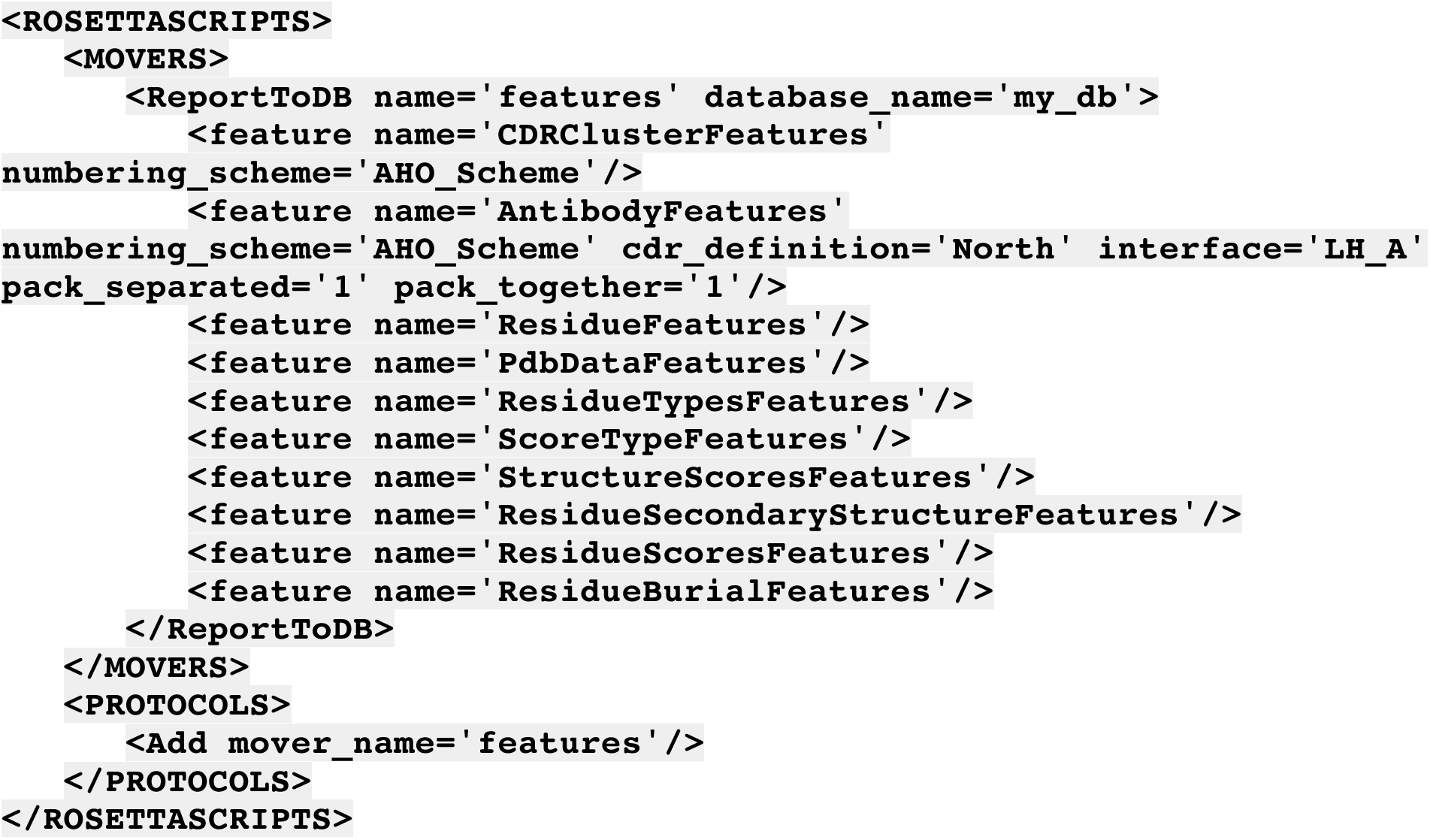

Once each database was created, extra output in each decoy PDB file (added through the option *-add_graft_log_to_pdb)*, was used to calculate recoveries and risk ratios through the creation and use the Bio-Jade AnalyzeRecovery module of the RAbD_BM subpackage.(https://bio-jade.readthedocs.io/en/latest/).

### 2. Dihedral, Epitope, and Paratope Constraints

Several constraint types are used by the Antibody Design framework to limit unproductive structural perturbations of the CDR regions and the relative orientation of the antibody-antigen interface while docking in the program. There are many constraint types with associated function types implemented in Rosetta. These constraints are evaluated via terms added to the Rosetta energy function. The Rosetta energy minimizer (which optimizes the conformation of the structure by finding the local energy minimum) can use these constraints to find optimal values that help to satisfy all the energy terms including the constraints. Within the Rosetta Antibody Design framework, the weight of these constraints can be set from command-line options. We can set parameters that govern whether these constraints are used throughout the protocol (where they also act as structural filters) or only in certain situations like minimization or docking (where they act only to guide the structure to an optimum conformation that satisfies the constraints).

The set of general Antibody constraint movers that were implemented consist of the *CDRDihedralConstraintMover, ParatopeSiteConstraintMover*, and the *ParatopeEpitopeSiteConstraintMover*.These Movers (a mover applies some change to a Pose or structure [1]) can be fine-tuned for specific design strategies using a number of user-accessible options and RosettaScripts [2].

The *CDRDihedralConstraintMover* places Circular Harmonic constraints on each ϕ and ψ dihedral angle of a given CDR as cluster-specific and general-use constraints. The equation for the Circular Harmonic constraint is as follows where x_0_ is the starting dihedral angle, x is the changed dihedral angle, and s is the standard deviation of x:

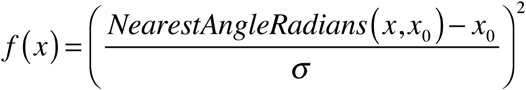

Constraints are added to help Rosetta maintain a particular loop structure during any backbone optimization. Dihedral constraints are used instead of coordinate constraints (which try to keep each atom at a particular Cartesian coordinate) in order to allow more natural, hinge-like motion of the CDR loops. Users of the protocol who wish to design antibodies without these constraints can set the weight of the dihedral constraint to zero via a command-line option.

The cluster-specific constraints have the value of x_0_ and standard deviation for each backbone ϕ and ψ dihedral angle at the angle mean and standard deviation of the members of the cluster. These constraints are output by PyIgClassify using a high-quality set of non-redundant data. The default behavior of the *CDRDihedralConstraintMover* is to add these constraints for a particular CDR only if there are enough members in the cluster to have reliable data. If data are scarce, then general dihedral constraints are added, with means at the current angles and a standard deviation that was originally compiled by taking the mean of the standard deviations of all CDR clusters. These angles can be set via command-line options. By default, we use these general dihedral constraints for H3, since it does not cluster well.

For the cluster-specific constraints (and other places in the protocol), we generally filter out outliers in the data as described below. This can be turned off through the use of an option that will load a different set of constraints compiled with structures that are not filtered for outliers. This can be useful if using outliers elsewhere in the protocol.

*SiteConstraints* are a set of atom-pair constraints that evaluate whether a residue interacts with some other chain or region – roughly, that it is (or is not) in a binding site. More specifically, if we have a *SiteConstraint* on a particular residue, that *SiteConstraint* consists of a set of distance constraints on the Cα atom from that residue to the Ca atom of all other residues in a set, typically the set being specific residues on another chain or chains. After each constraint is evaluated, *only* the constraint giving the lowest score is used as the *SiteConstraint* energy for that residue. These *SiteConstraints* use a Flat Harmonic function by default:

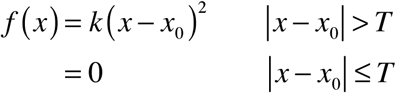

Values of the standard deviation are set at 1 Å, while the tolerance is set at the interface distance of the protocol (8 Å default), which means that there is no penalty for the *SiteConstraint* except at distances greater than this distance.

The *ParatopeSiteConstraintMover* adds *SiteConstraints* between each CDR residue and the antigen. This helps to keep the CDR paratope at the interface during docking; without it, docking can use the whole of the antibody surface instead of just the paratope and this can be seen in resulting models. These paratope constraints are added automatically in the program, and the CDRs of the paratope can be controlled through an option.

The *ParatopeEpitopeSiteConstraintMover* adds *SiteConstraints* from the epitope residues to the paratope residues and from the paratope to the epitope. Target epitope residues can be specified via command-line or automatically detected via the set interface distance. These constraints are off by default, but if they are enabled, they are set instead of the *ParatopeSiteConstraintMover* and help to keep the paratope and the epitope in contact during design when the docking component of the algorithm is enabled.

### 3. Outlier Control

In our original clustering of the antibody CDR structures, an affinity propagation clustering technique was used on a carefully curated dataset of high-resolution structures and few outliers [3]. In order to match new CDR structures with a proper cluster from that original clustering, we use the dihedral angle metric originally used for the affinity propagation, but measure it against the centroid (representative structure) of all clusters of the same length. The cluster with the lowest dihedral distance is assigned as the cluster for that structure [4].

While this is useful to assign CDRs of known length to a particular cluster, many structures become outliers of the particular cluster and would have formed their own cluster if clustering was repeated (Kelow and Dunbrack, in preparation). To optimize our CDR profiles, constraints, and other aspects of the design program for an updated database, we needed to quantitatively define what would be considered an outlier.

We used both the dihedral distance metric and RMSD of all backbone atoms to help define an outlier. In order to visualize the breadth of each cluster, we generated PyMol sessions of each of the clusters using python and PyRosetta [5] by aligning the CDRs to their cluster center either using all backbone heavy atoms or by aligning only the stem region (three framework residues on either side of the CDR loop). We also generated plots of dihedral distance versus RMSD and length versus RMSD for both alignment types and for each length and cluster, where high RMSD can be seen even with lower dihedral distance, especially when only the stem was aligned. We then used these plots and the PyMol visualizations for each CDR cluster to define two outlier definitions – one conservative and one liberal (used as the default). We calculate the RMSD for these definitions through the full CDR alignments as the stem alignment can result in very high RMSD for low dihedral angle distances, attributable to hinge-like motions in the CDR:

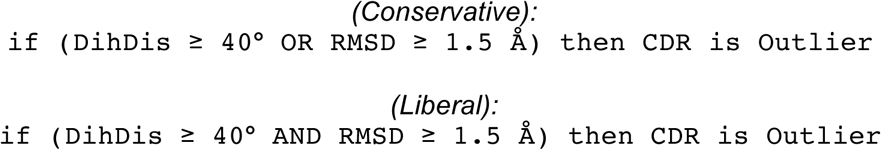

Outlier control is handled as an option in the Antibody Design framework, where each set of data used by the framework is first generated with and without outliers and using both the liberal and conservative definition of an outlier. For smaller clusters or H3 (which only cluster well at lengths ≤ 9), outliers may be useful in the design search and an option will switch all aspects of the framework to include outliers for sequence and dihedral constraint statistics as well as graft sets. By default, outliers are left out, but used for H3 since it does not cluster well.

### 4. Antibody Feature Analysis

Three FeatureReporters were developed as a part of the Rosetta Feature Reporter framework [6–8] to aid in the modeling and design of antibody structures. Each of these can be used through the RosettaScript framework on a list of structures. The physical attributes reported are output to a relational database, such as SQLITE3, across multiple tables for further analysis. These databases can easily be converted into CSV files or read by available packages in R and Python.

The *CDRClusterFeature* reporter identifies all North/Dunbrack CDR clusters in an antibody to the closest cluster centroid using the same metric described in PyIgClassify as well as information pertaining to the dihedral distances of the CDRs [4]. It is the primary FeatureReporter used in benchmarking length and cluster recovery. The database tables output by the *CDRClusterFeatures* are detailed in Table D in S1 Supporting Information.

The *InterfaceFeature Reporter*, detailed in Table E in S1 Supporting Information, analyzes protein-protein and protein-ligand interfaces, outputting a number of different tables and physical data. Much of the analysis is done through the Rosetta InterfaceAnalyzer [9,10] which we have updated. The InterfaceAnalyzer calculates differences in scoring (such as an estimate of the interface **Δ**G – the enthalpic component of the full binding free energy) by physically separating the interface components (such as antibody from antigen) and optimizing interface residue side chains - both in the complexed and separated conformations. An interface distance of 6 Å is used as the default interface distance.

Separate tables are output for the overall complex, the individual proteins in the complex, and the interface residues. The main data output by this Reporter are the estimated binding energy (**Δ**G) of the complex in Rosetta Energy Units (REU), the change in solvent accessible surface area upon binding (**Δ**SASA) using the Le Grand SASA calculation method [11], the Lawrence and Colman shape complementary of the interface (sc_value) [12], the packing quality (packstat) [13], and the number of unsaturated hydrogen bonds in the complex [10].

We added alternative SASA radius sets to Rosetta, with the standard, now-defunct radii changing from the default to ‘legacy’. We implemented a variety of radius sets found in the literature and used in various structural modeling programs in which they either implicitly or explicitly include hydrogen atoms. Once a particular radius set is used, the SASA machinery will change its consideration of implicitly or explicitly including hydrogen atoms during the calculation depending on the set.

The atomic radius set with implicit hydrogens is the one used by the program Naccess, a popular program used for the calculation of SASA [14]. This set was derived by Chothia in his seminal 1976 paper [15], while explicit hydrogen radius sets include the legacy radii, the Rosetta Lennard-Jones (LJ) radii (which are mostly the same as the LJ radii from the CHARMM molecular dynamics program [16]), and the radii used by the program *reduce* (a program for the placement of hydrogens onto molecular models and crystal structures) [17], originating from physical data obtained from Bondi [18] and Gavezzotti [19]. The reduce radius set is now the default in Rosetta.

We implemented the *AntibodyFeature* Reporter, a type of *InterfaceFeature* Reporter specific for antibody and antibody-antigen interfaces, while outputting a number of *additional* metrics for antibodies and CDRs. Some of the main metrics include CDR, antibody, and paratope charge, ΔG and ΔSASA, H3 kink statistics, number of contacts, and packing angle statistics [20]. The packing angle is a measure of the relative orientation between the light and heavy antibody chains. It uses four conserved residues of each chain in the framework beta-sheets at the VL and VH interface and principal component analysis to define four centroid points and a dihedral angle for which to quantify the orientation [21].

A full list of the metrics and tables output by the *AntibodyFeature* Reporter can be found in Table F in S1 Supporting Information. All tables output by the *InterfaceFeature* Reporter are output by the *AntibodyFeature* Reporter for specific antibody interfaces specified where A is the antigen: LH-A, LH, L-A, H-A.

## S1 Supporting Information

**Adolf-Bryfogle et al., RosettaAntibodyDesign: A general and flexible framework for computational antibody design**

**Fig. A.**
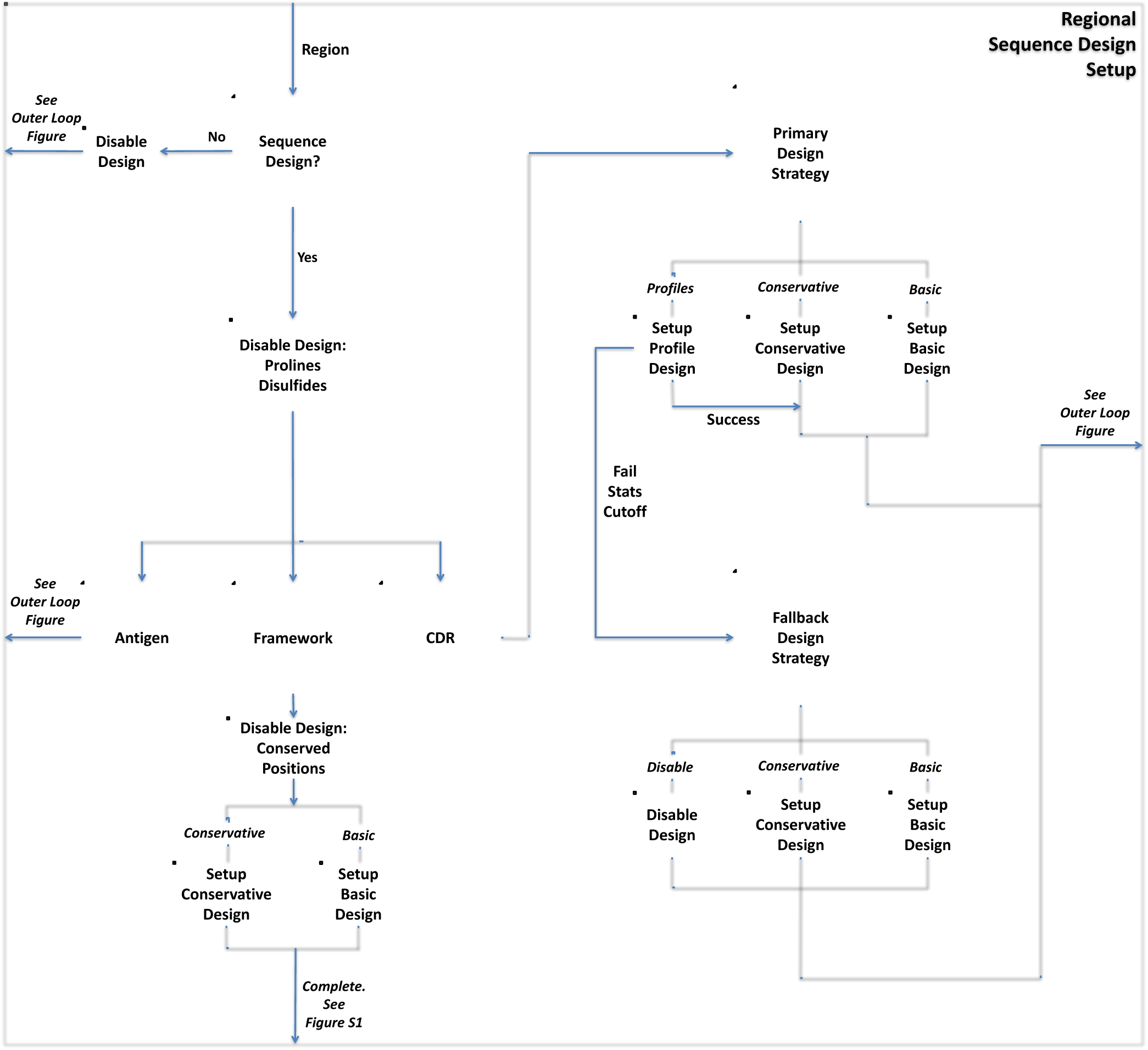
Regional sequence design setup. For each region (Antigen, Framework, Each CDR), whether that region is set to design is determined from the program options and CDR Instructions File. If the region is set to design, design is disabled for prolines and disulfide-bonded cysteine residues within the region by default. If it is an antigen region, we use “basic design” (standard Rosetta design) and exit the setup. By default, the framework region is held fixed in sequence space. But if framework design is enabled, we disable design on completely conserved positions, such as the tryptophan immediately after the H3 loop, and perform conservative or basic design on the rest. Finally, when designing a CDR region, we set up the Primary Sequence Design strategy that is set by the CDR Instructions File. If the Primary Sequence Design strategy is to use the CDR cluster-based profiles and there is scarce data, we use the set Fallback Design Strategy. After the CDR Sequence Design strategy is done, we have completed the setup for sequence design.

**Fig. B.**
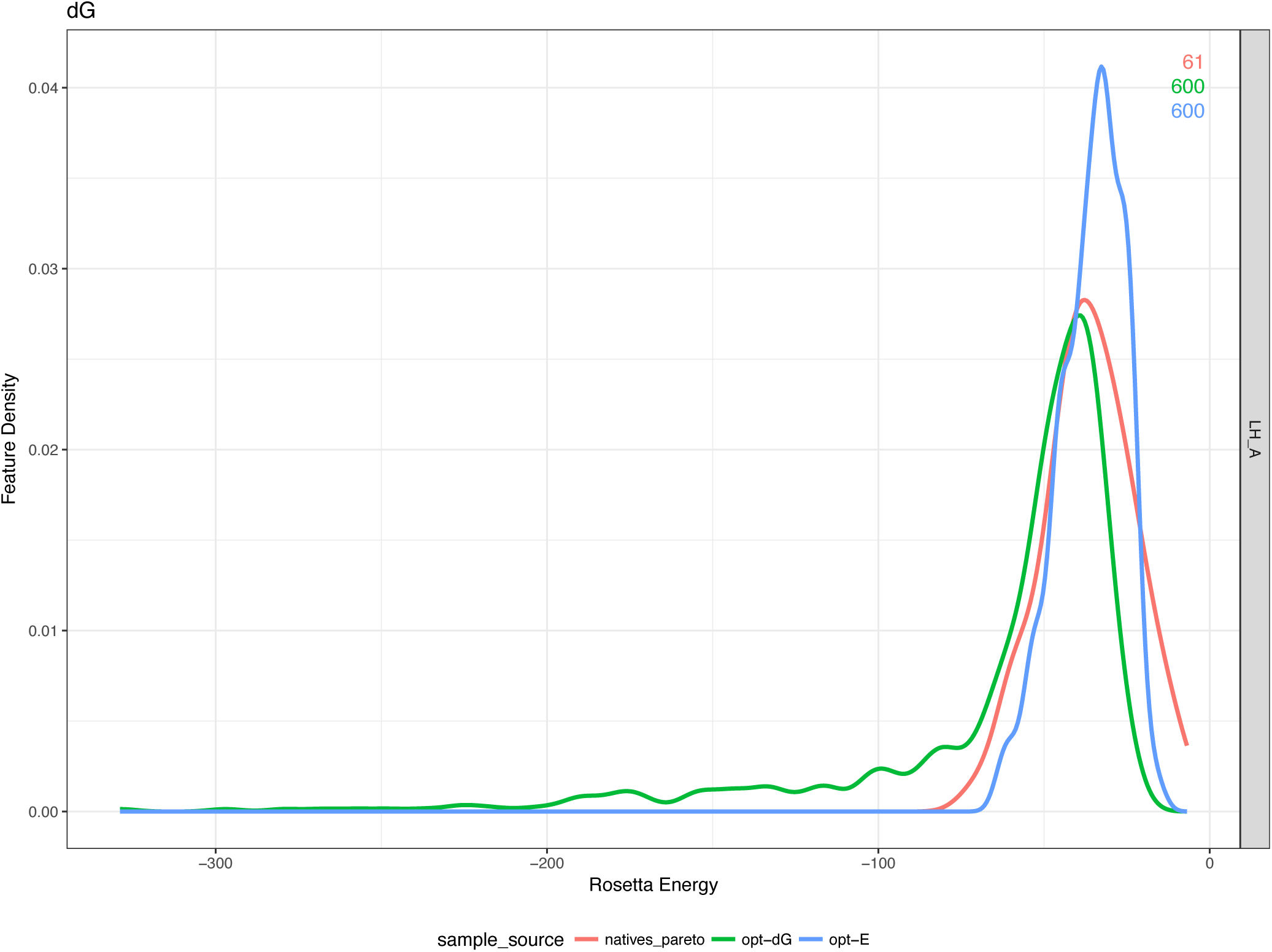
Benchmark Rosetta interface energies. Kernel density estimates (KDE) and averages of the Rosetta Interface Energy (dG) of the top 10% of the 100 decoys for each of the 60 antibody-antigen complexes in the opt-E and opt-dG benchmarks as well as the natives after optimization. As expected, opt-dG produces lower dG scores that are more similar to native than opt-E.

**Fig. C.**
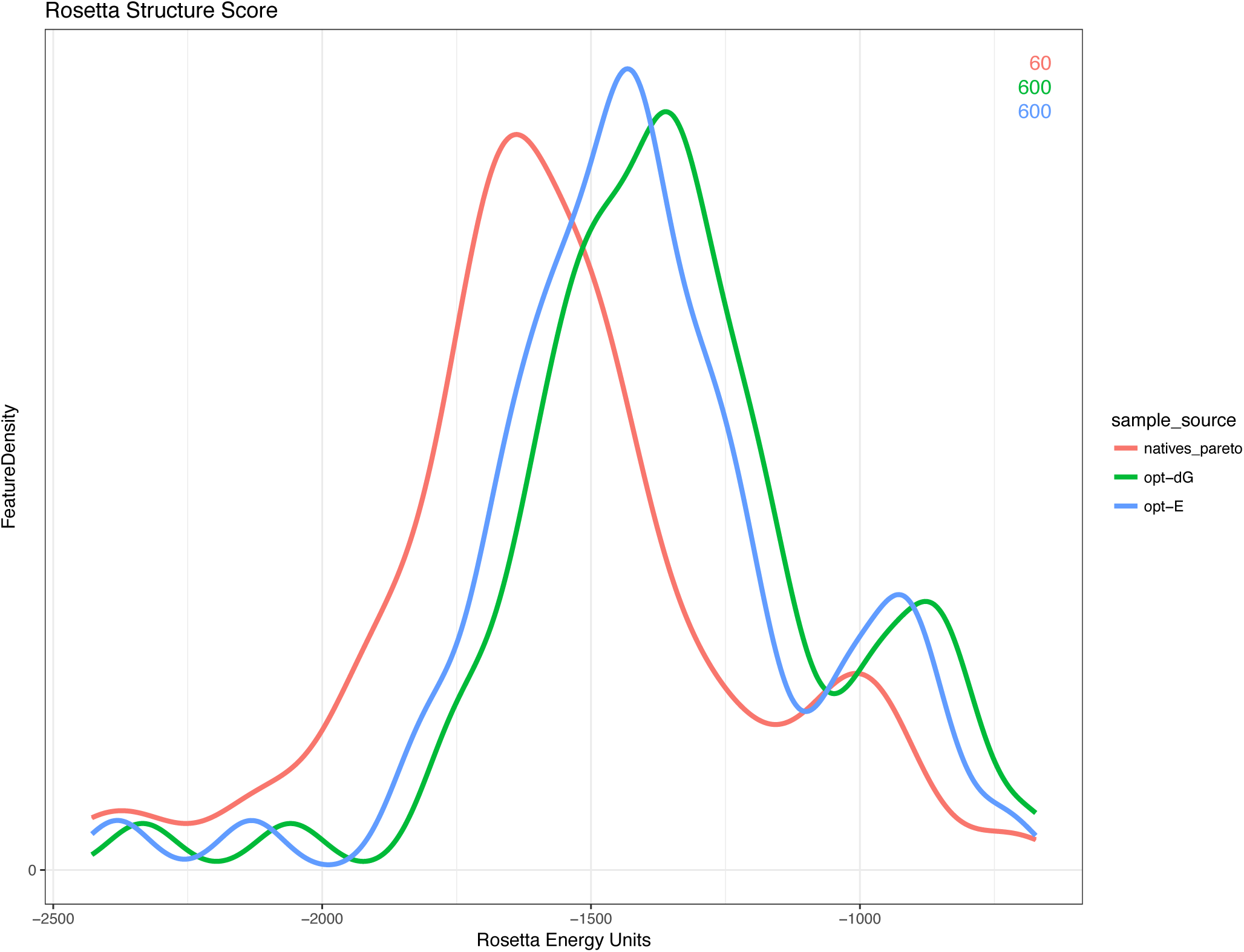
Benchmark total Rosetta energies. Kernel density estimates of total Rosetta energy (REU) of the opt-E vs opt-dG benchmark decoy set using the current Rosetta Energy function (REF2015). Densities for the natives and the top 10% of decoys are shown.

**Fig. D.**
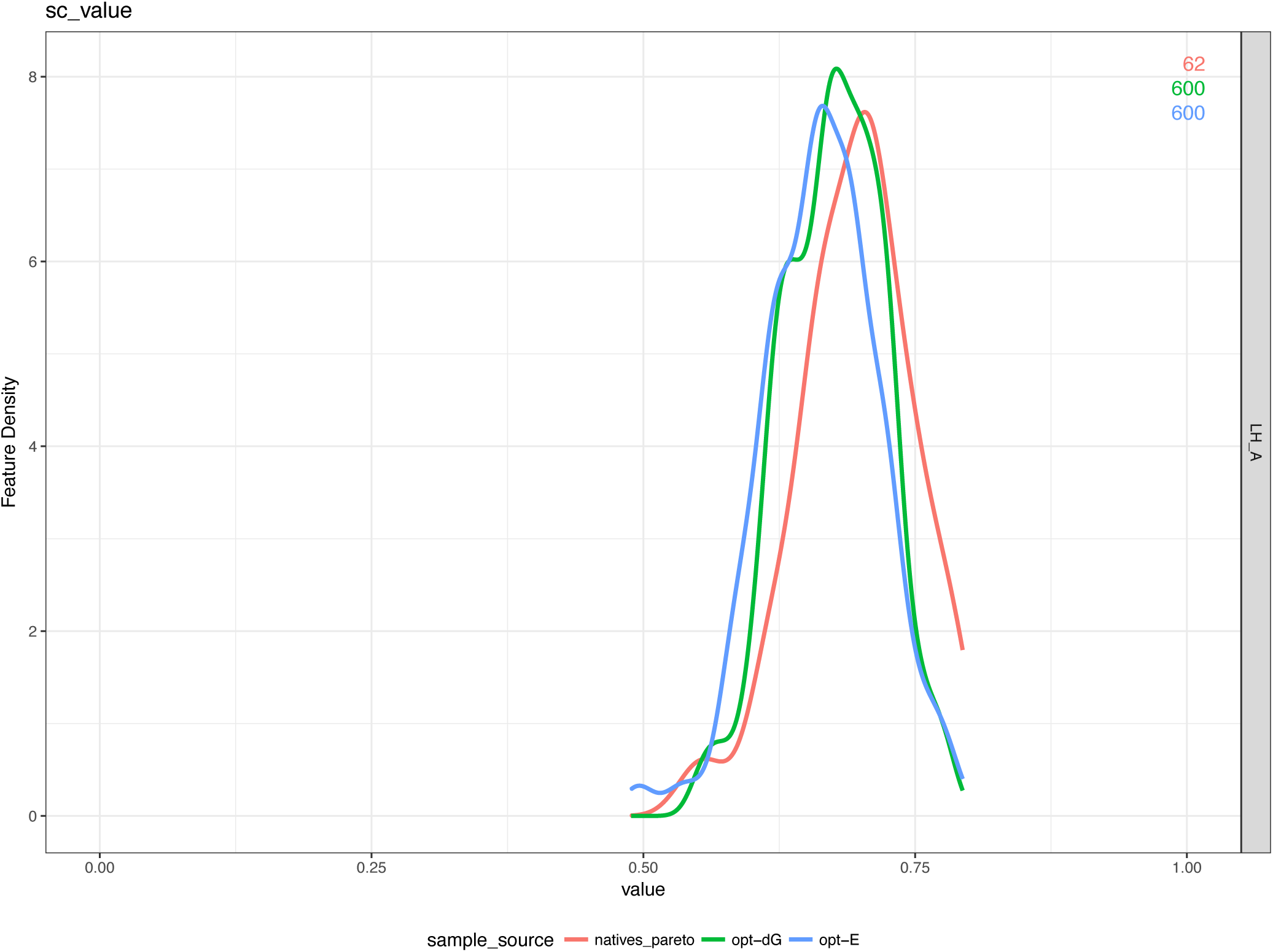
Benchmark shape complementarities. Kernel density estimates of the Lawrence and Colman Shape Complementarity value (*sc_value*) of the opt-E and opt-dG benchmark decoys for 60 antibody-antigen complexes using the Rosetta *AntibodyFeature* reporter.

**Fig. E.**
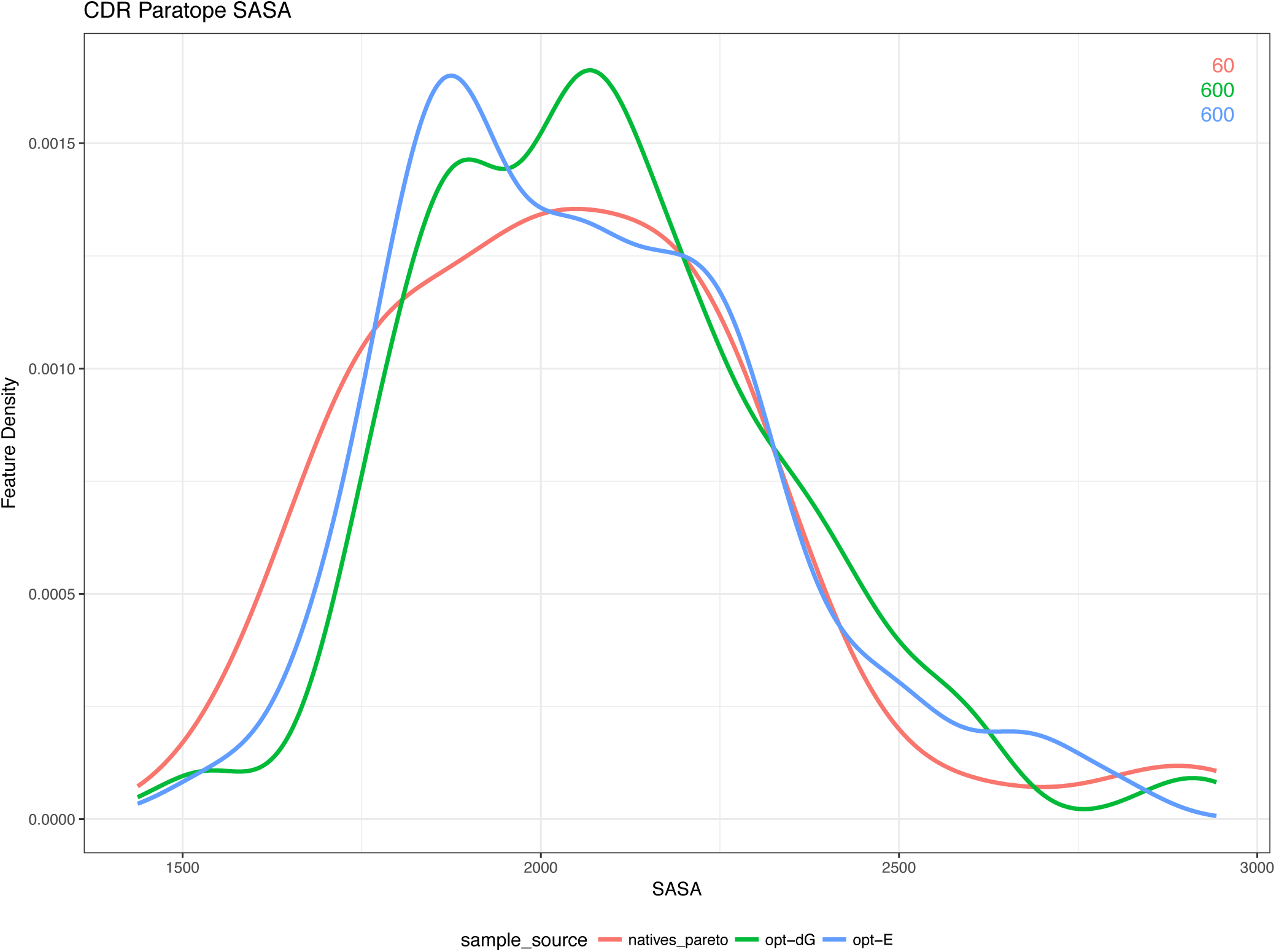
Benchmark solvent accessible surface areas. Kernel density estimates of the buried Solvent Accessible Surface Area (**Δ**SASA) of the opt-E and opt-dG 60 antibody benchmark decoy set at the antibody/antigen interface compared to the relaxed native structures.

**Fig. F.**
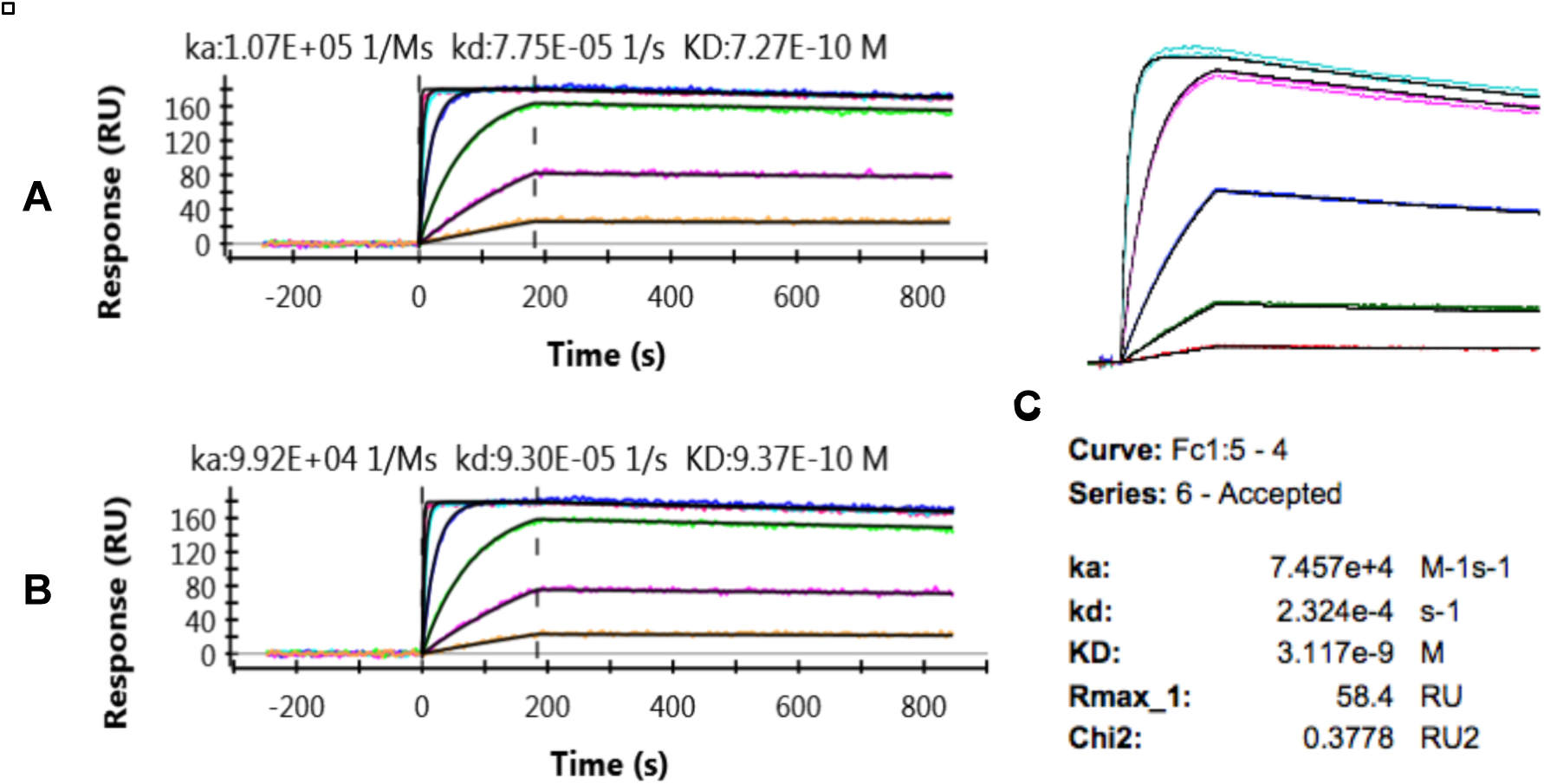
Kinetic sensorgrams of design L1_10 to Bee Hyaluronidase. **(A)** *XPR Repeat 1*; **(B)** *XPR Repeat 2;* **(C)** *Biacore 4000*.

**Fig. G.**
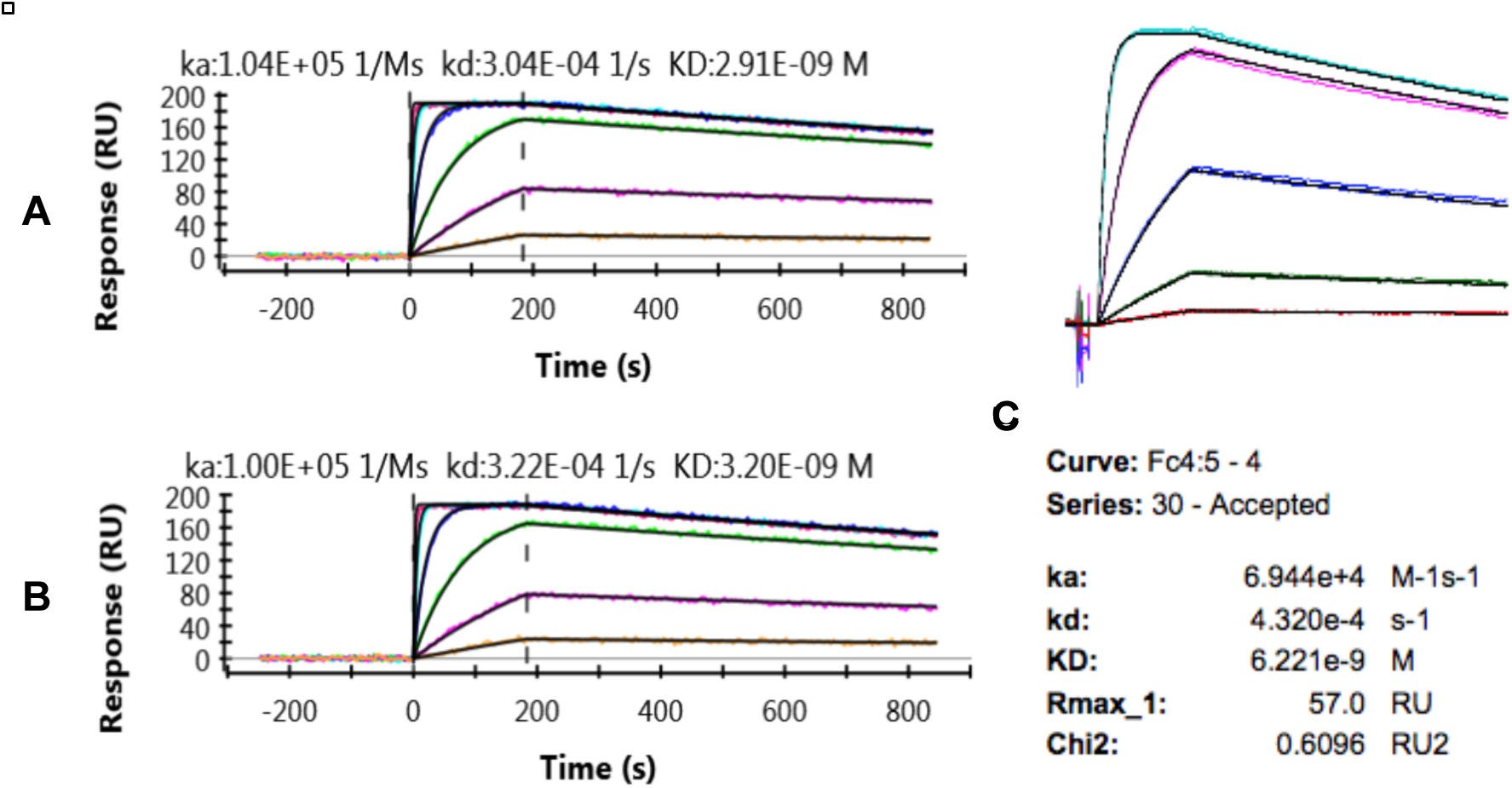
Kinetic sensorgrams of design L1_5 to Bee Hyaluronidase. **(A)** *XPR Repeat 1*; **(B)** *XPR Repeat 2*; **(C)** *Biacore 4000*.

**Fig. H.**
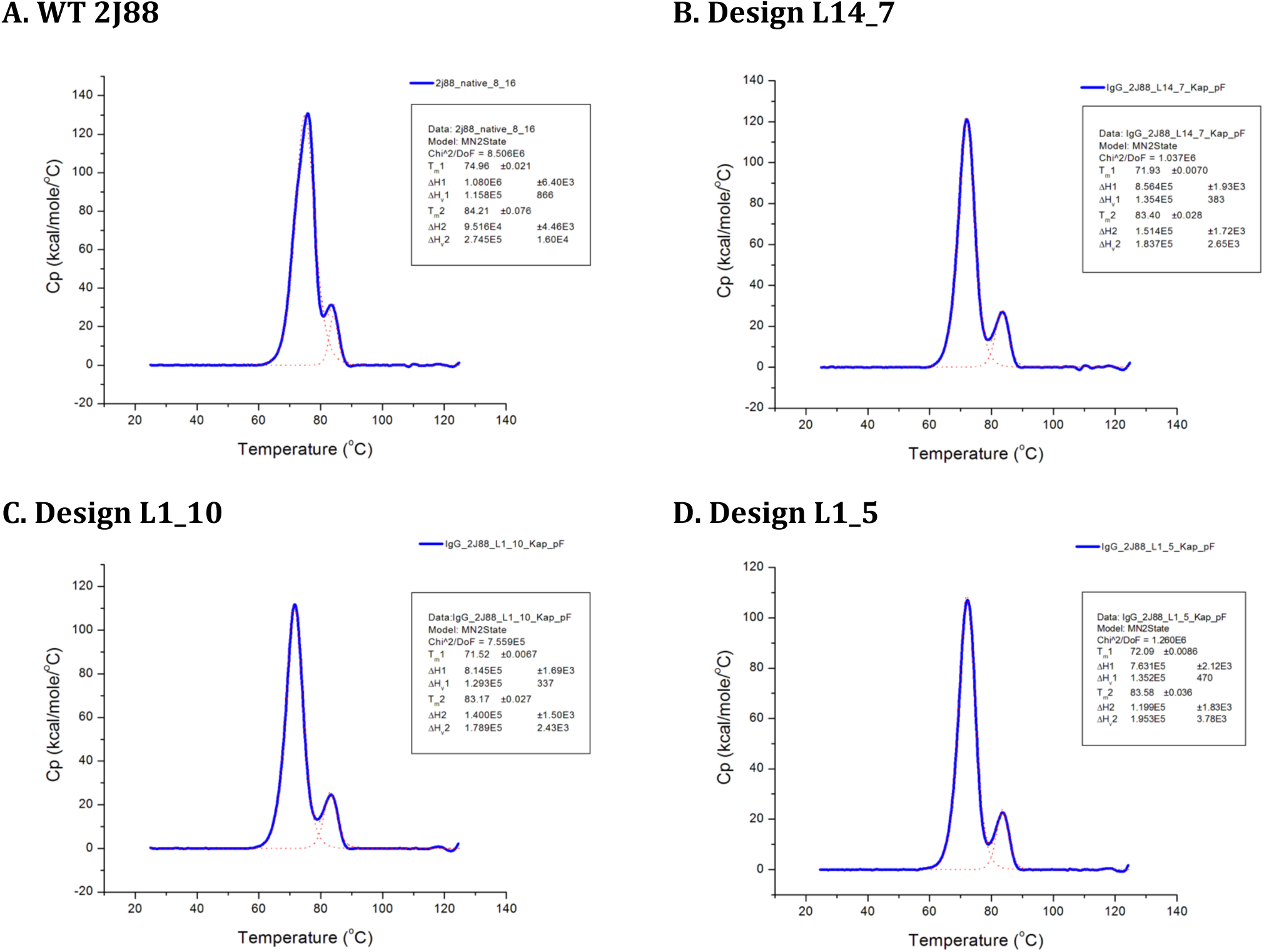
Thermostability measurements of WT 2J88 antibody and designs by Differential Scanning Calorimetry (DSC). (**A**) WT 2J88 thermostability; **(B)** L14_7 design thermostability; **(C)** L1_10 design thermostability; **(D)** L1_5 design thermostability.

**Fig. I.**
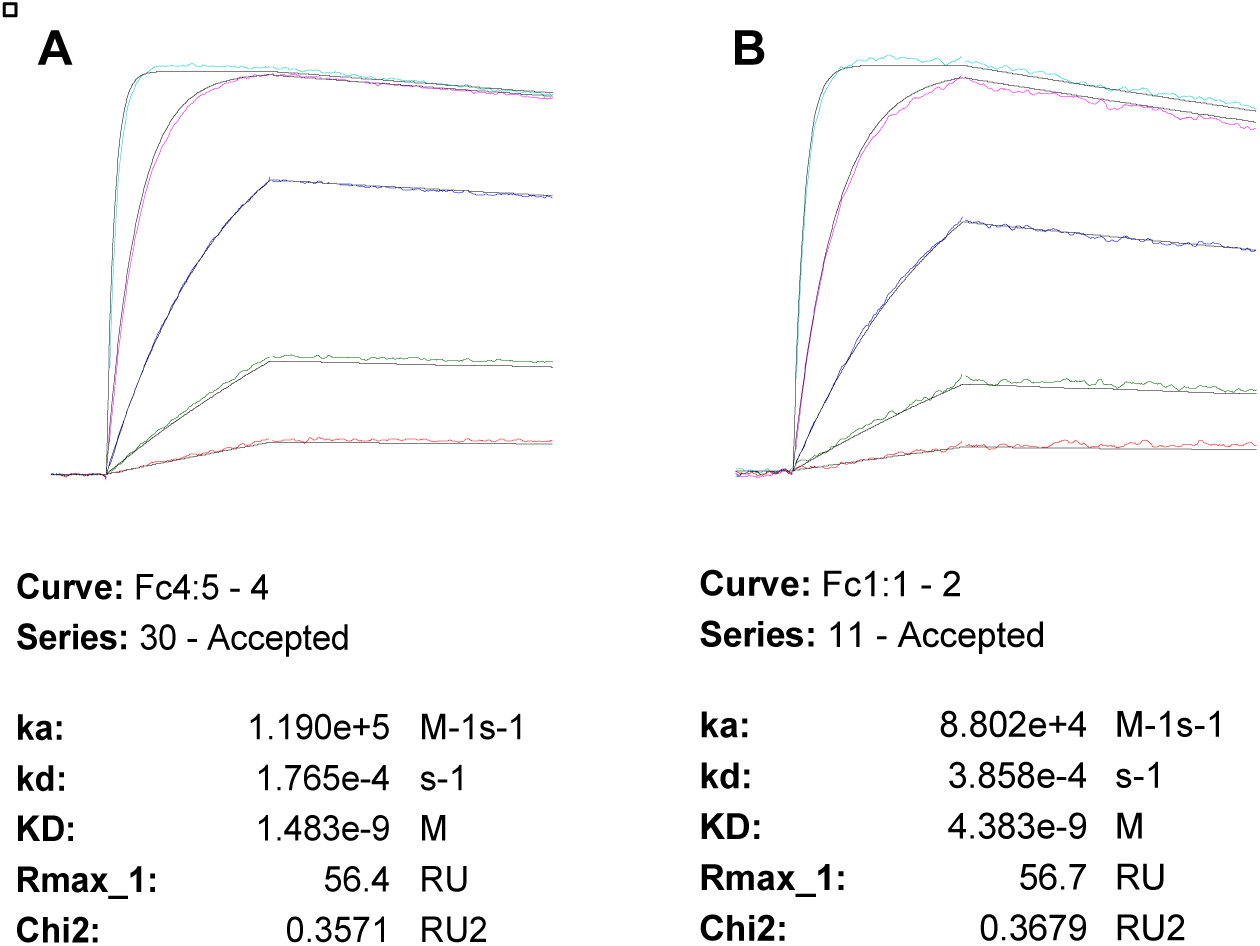
L14_7 Mutants with WT residue at position 38. (**A**) L14_7. (**B**) L14_7 K38Y.

**Fig. J.**
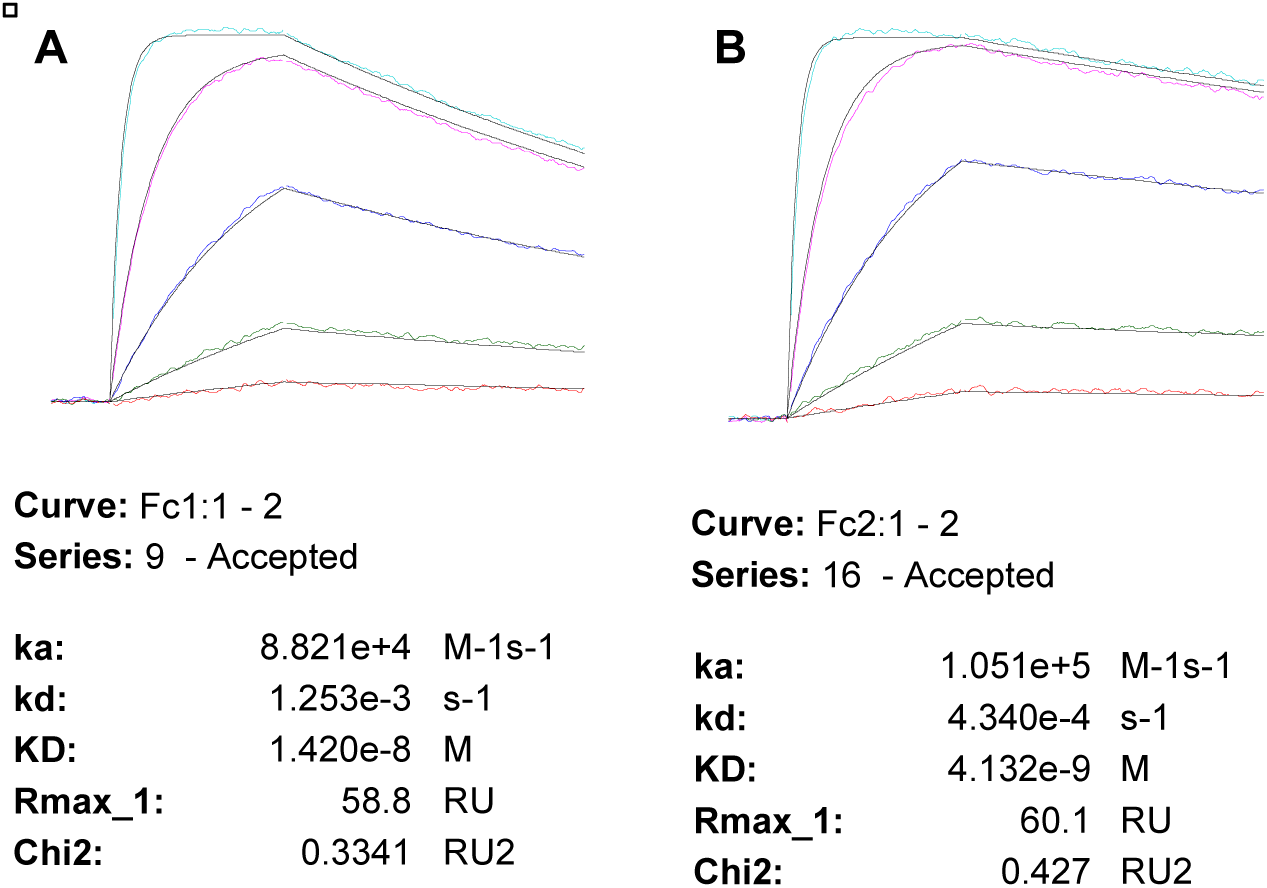
2J88 WT Mutants with L14_7 residue at position 38. **(A)** 2J88 WT; **(B)** 2J88 Y38K

**Fig. K.**
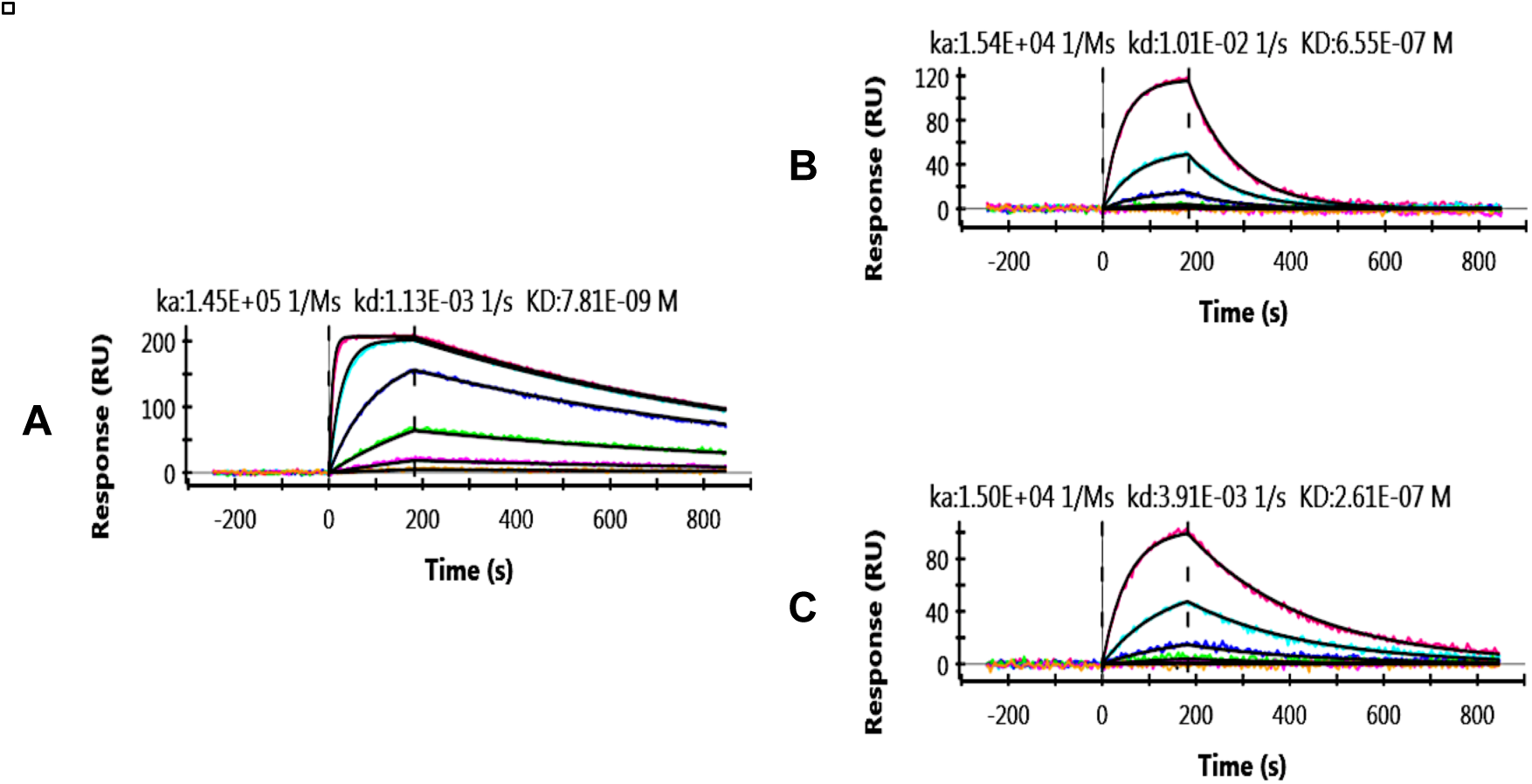
2J88 WT Mutants with L14_7 residue at position 38. **(A)** 2J88 WT; **(B)** L1_4 Design; **(C)** L1_4 S36V.

**Fig. L.**
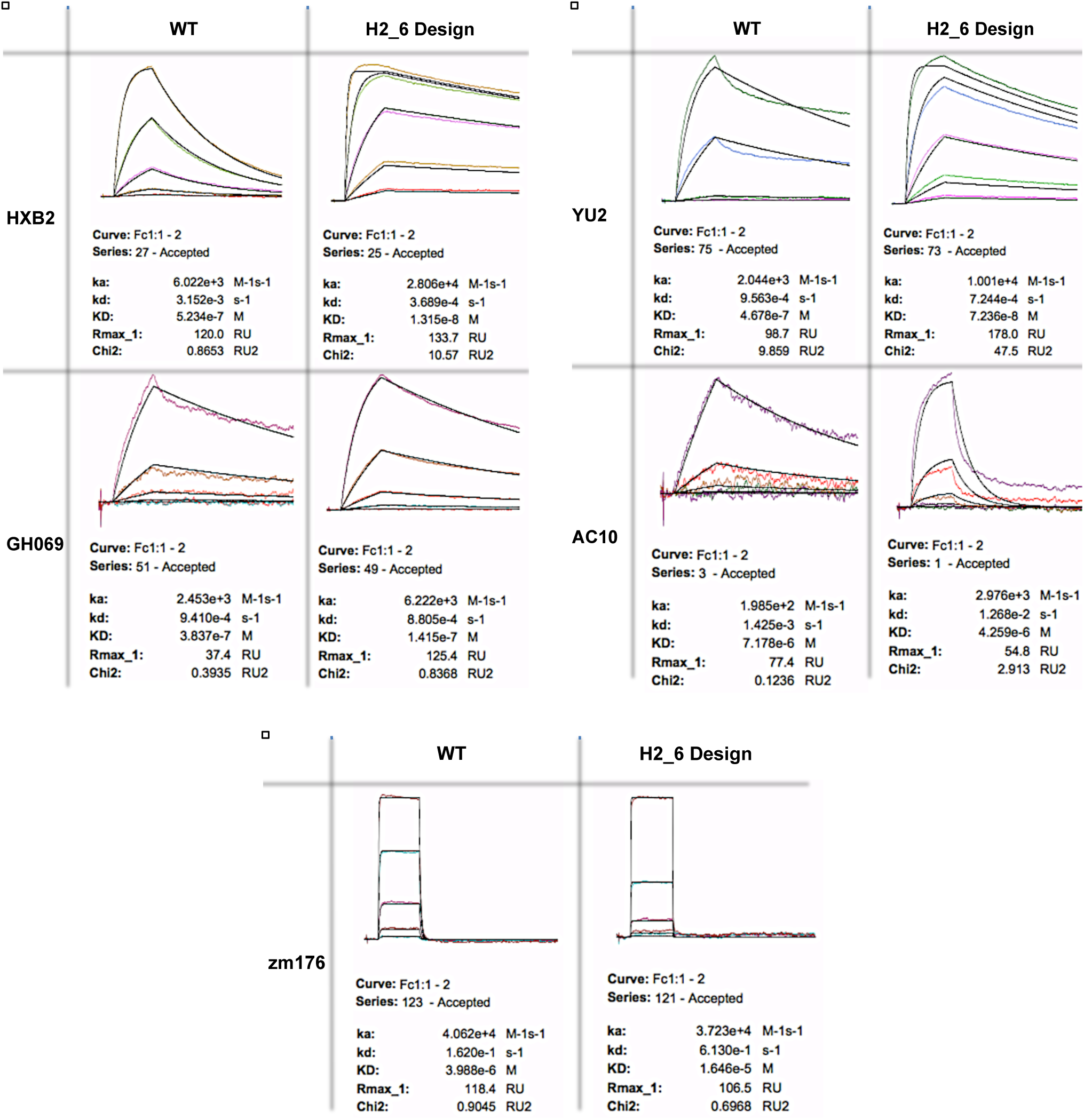
Kinetic sensorgrams of CH103 antibody and H2_6 design to a panel of gp120s. Binding studies were performed on a Biacore 4000.

**Fig. M.**
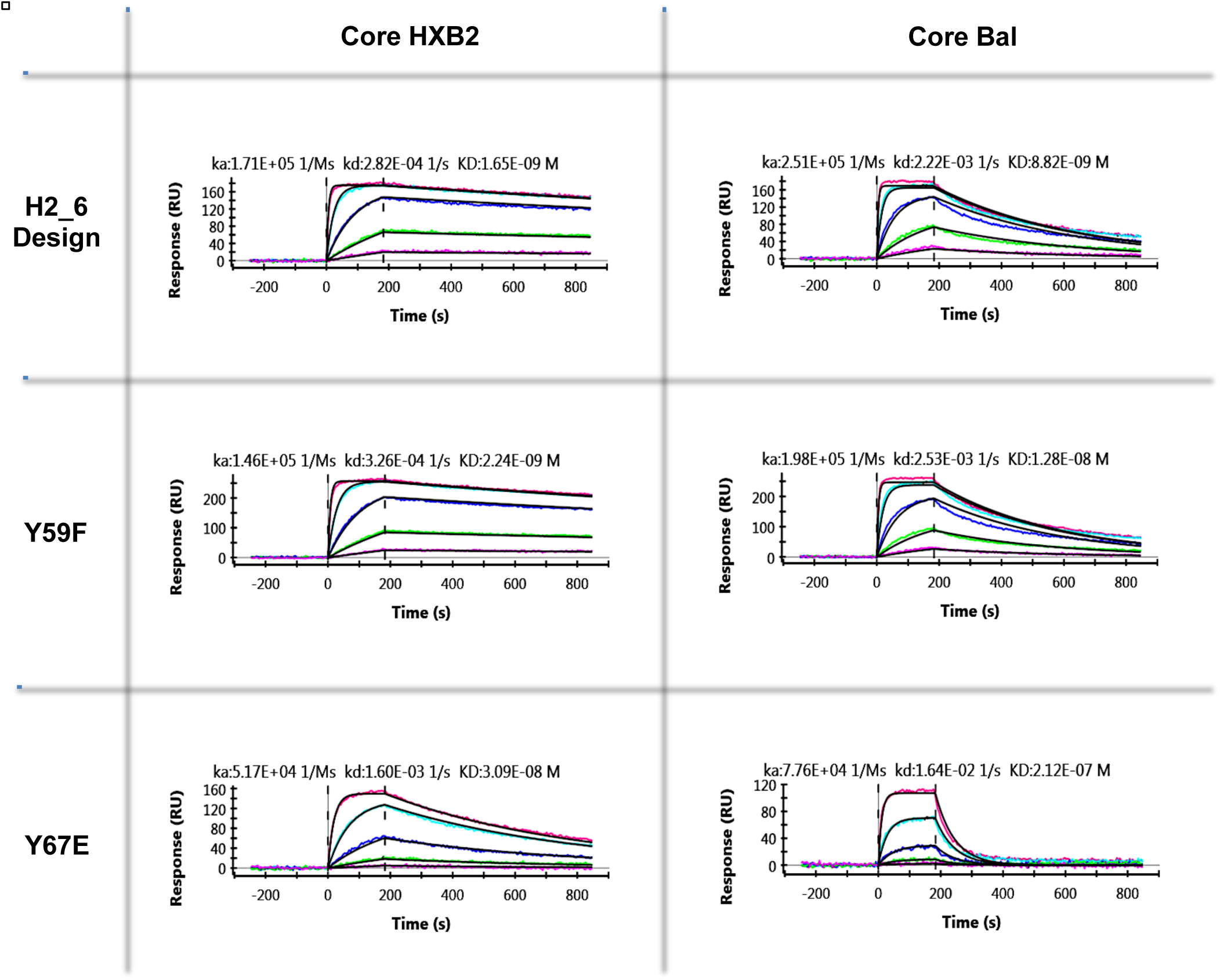
Kinetic Sensorgrams (ProteON XPR) of CH103, H2_6 design mutants with residues from CH103 WT.

**Fig. N.**
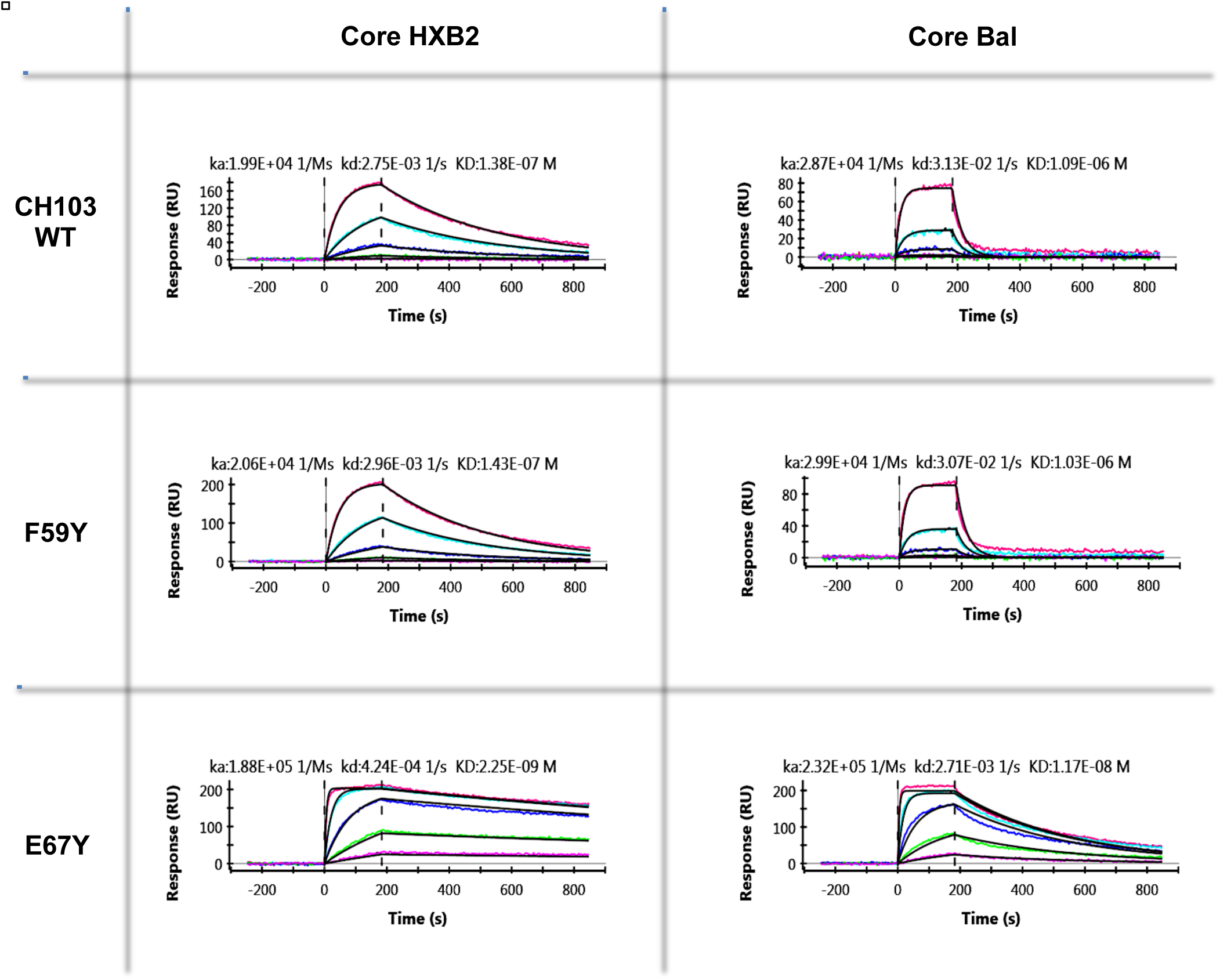
Kinetic Sensorgrams (ProteON XPR) of CH103 WT with mutations based on the H2_6 design.

**Fig. O.**
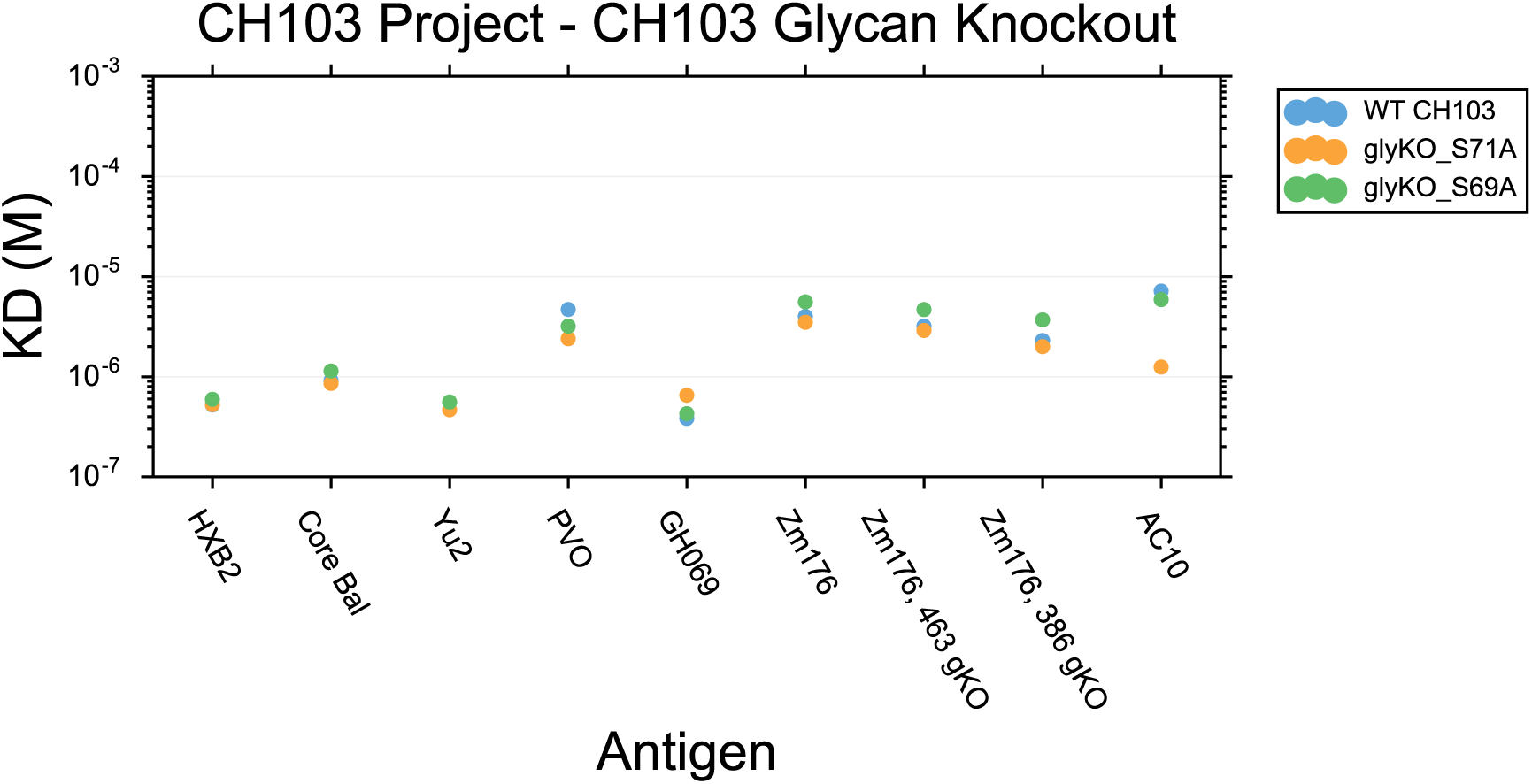
Paratope mutations. Two potential glycan sites were located in proximity to the antibody paratope for the CH103 antibody (PDB: 4JAN). These sites were mutated from serine to alanine mutations at position 69 and 71 in the heavy chain (AHo-Numbering). Binding affinity (K_d_) is shown from Biacore experiments to nine expressed GP120s.

**Fig. P.**
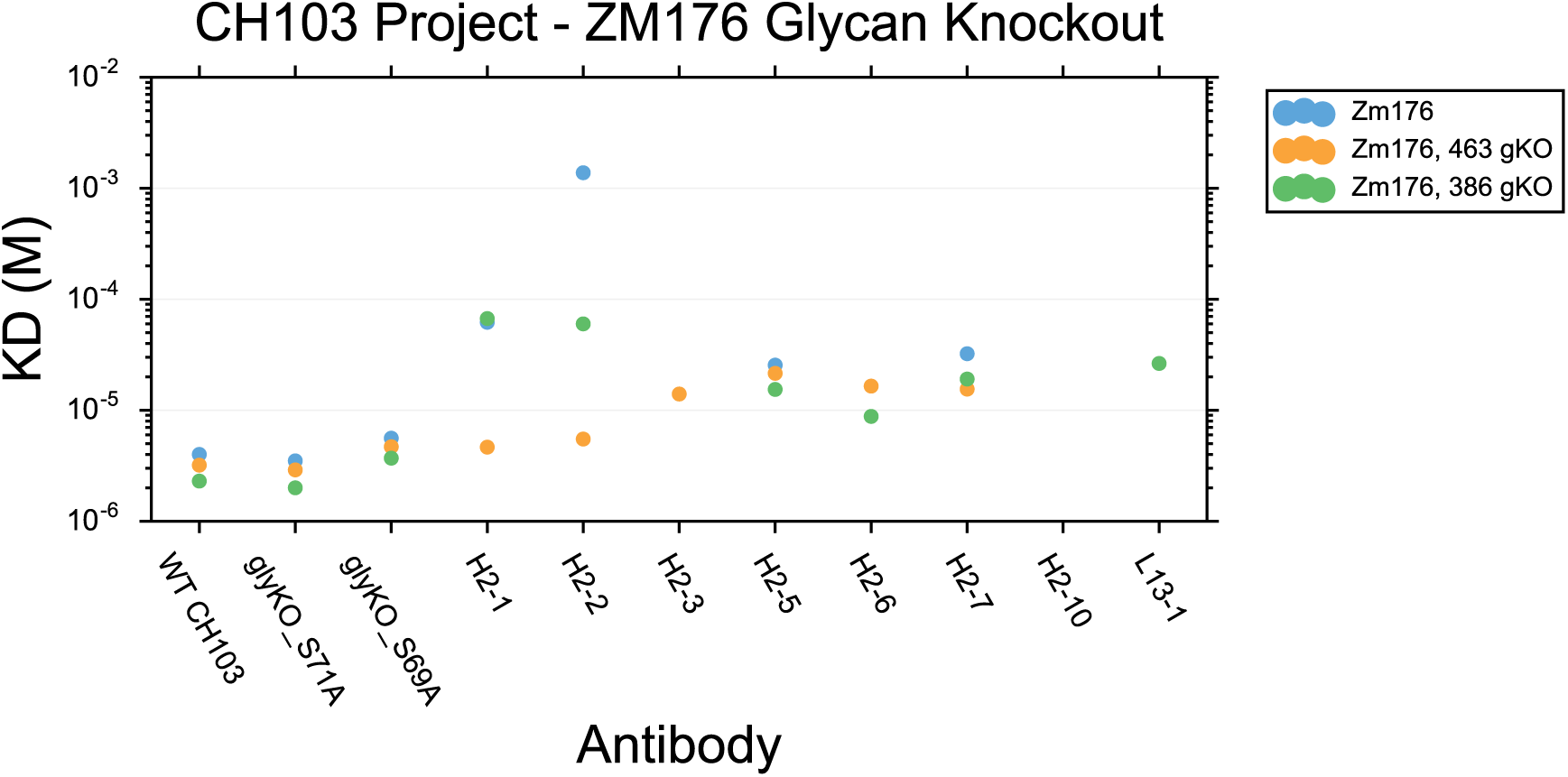
Epitope mutations. Potential glycan sites were located in proximity to the GP120-CH103 epitope at positions 386 and 463 of Zm176 (PDB ID 4JAN). Each of these sites were knocked out using Serine to Alanine mutations. Binding affinity (K_d_) is shown from Biacore experiments to nine expressed GP120s.

**Fig. Q.**
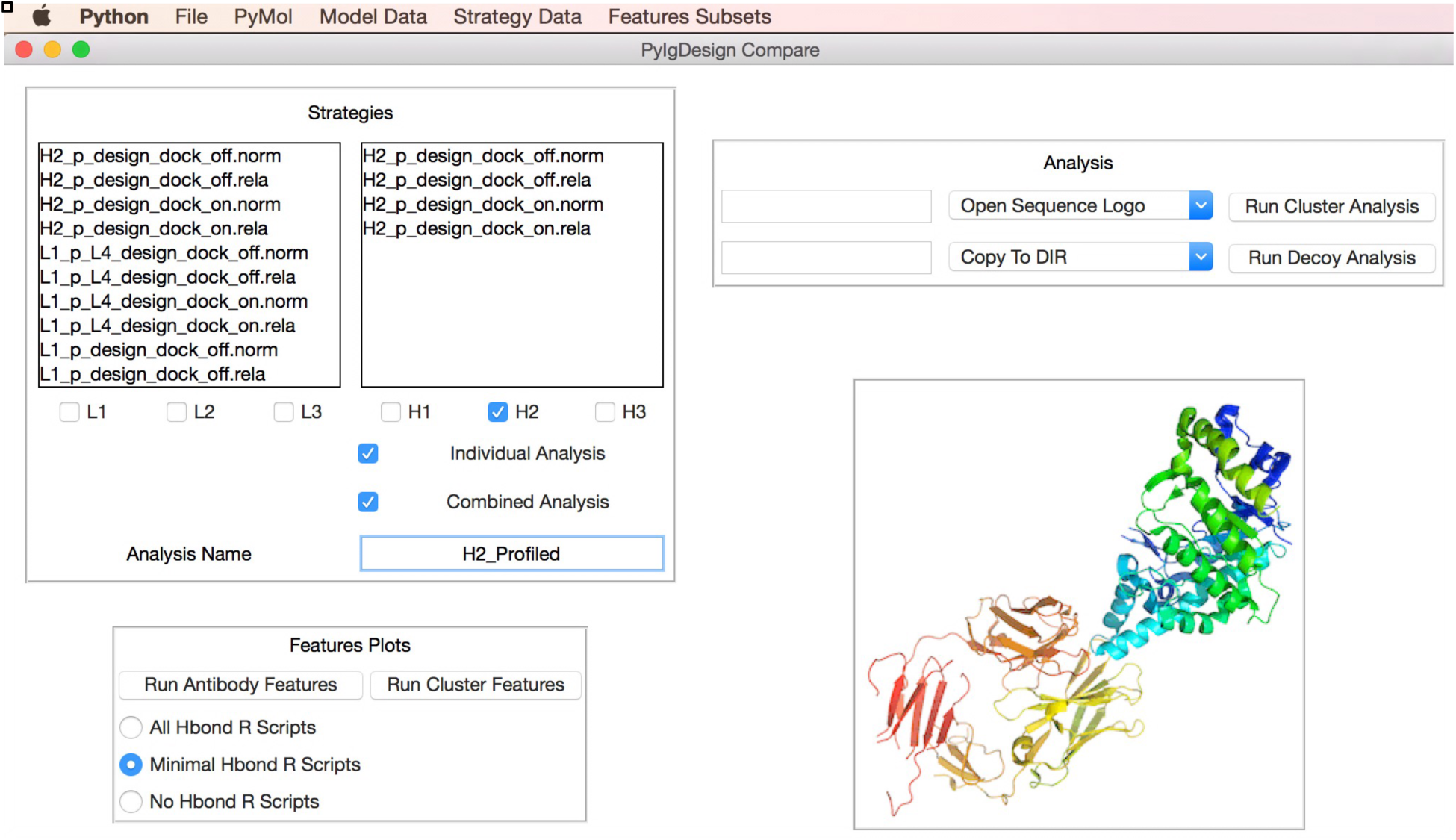
Jade Antibody Design analysis Graphical User Interface (GUI)

**Table A.**
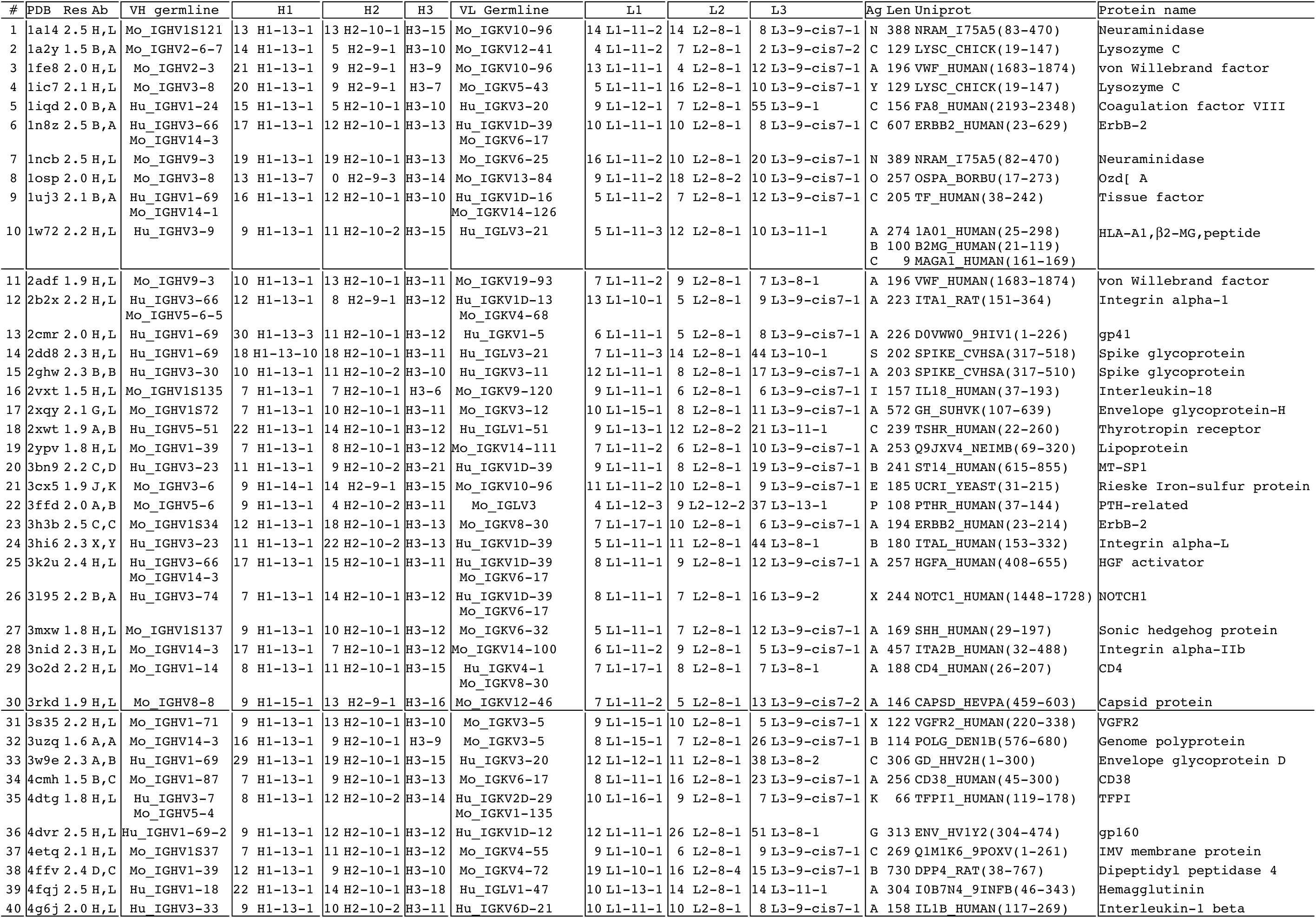
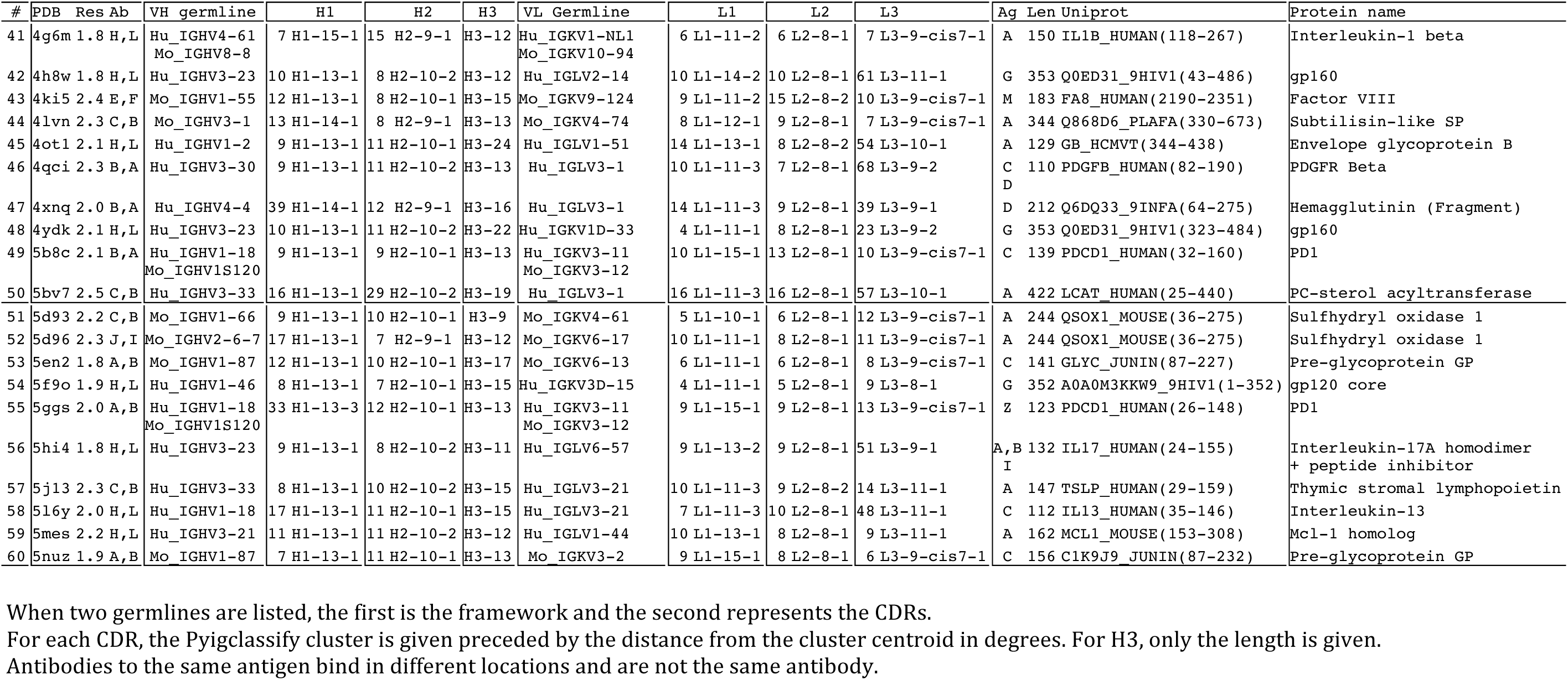
Detailed Data on Benchmark Antibody Complexes

**Table B.**
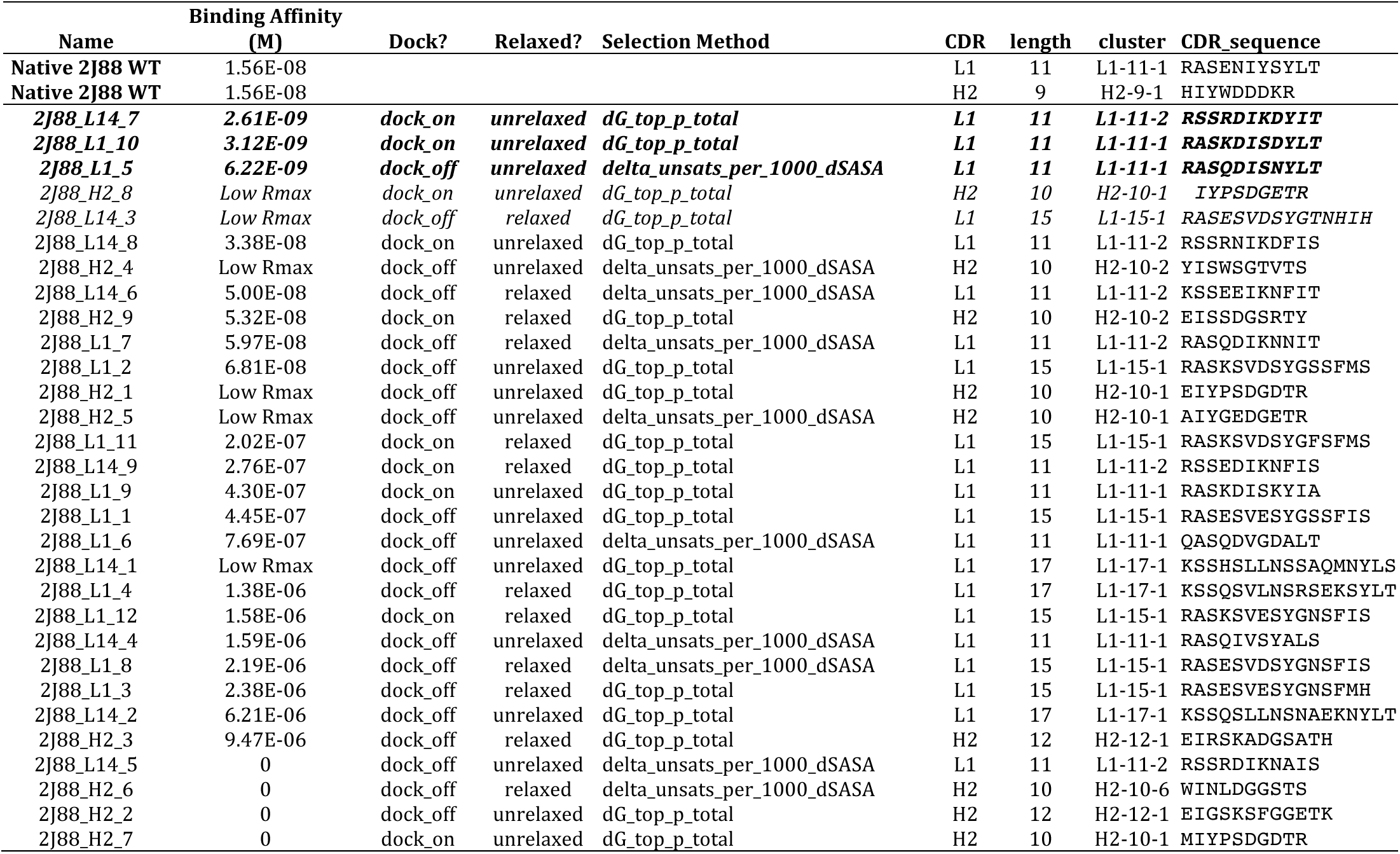
Binding affinity, CDR design identity, and selection strategies of the expressed and tested 2J88 designs from the Biacore 4000 results.

**Table C.**
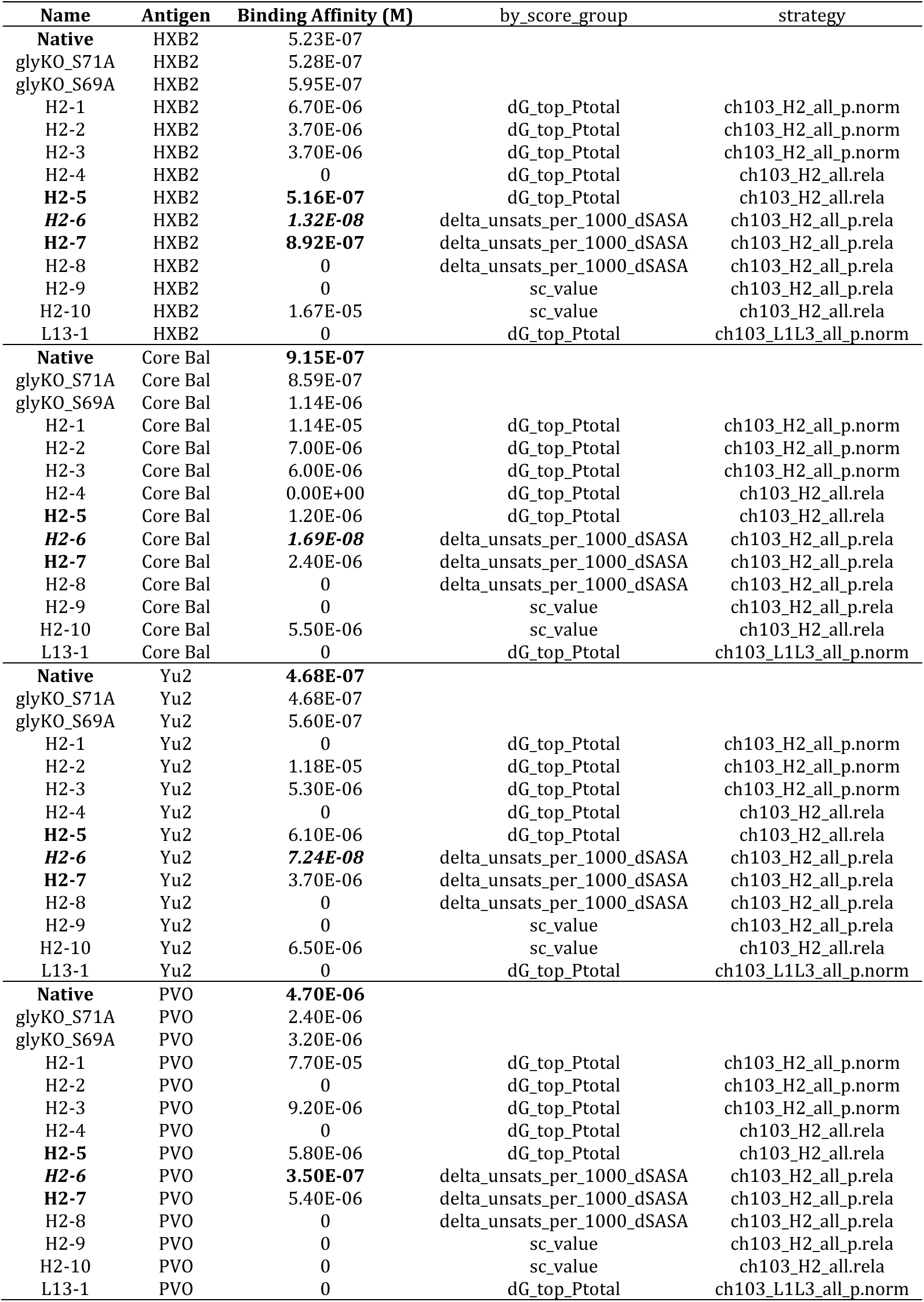
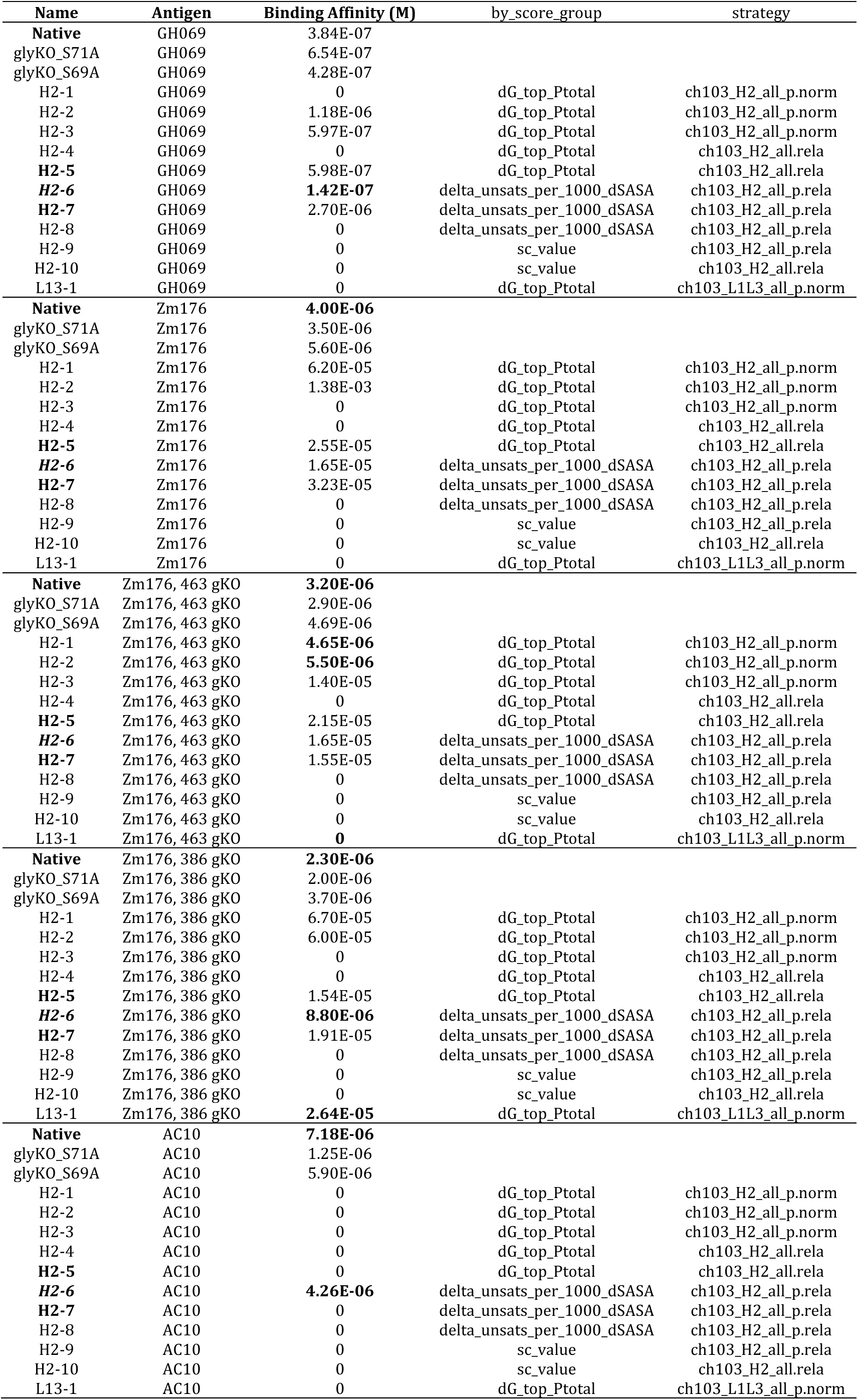
Binding affinity of expressed and purified CH103 designs and their selection strategies.

**Table D.** *CDRClusterFeature* reporter tables

| Column | Description |
| --- | --- |
| ***Table cdr_clusters*** |  |
| struct_id | Structure number – used to query other tables for same loaded structure |
| cdr_start | Starting number of CDR |
| cdr_end | Ending number of CDR |
| chain | Chain of CDR – L or H |
| CDR | Identity of CDR |
| length | Length of CDR |
| cluster | Identified cluster of CDR |
| dis | Calculated Dihedral Distance to cluster center |
| normDis | Dihedral Distance normalized by length (radians) |
| normDis_deg | Dihedral Distance normalized by length (degrees) |
| sequence | Amino acid sequence of the CDR |

**Table E.** *InterfaceFeature* reporter tables

| Column | Description |
| --- | --- |
| ***Table interfaces*** |  |
| struct_id | Structure number – used to query other tables for same loaded structure |
| interface | Interface ID – The interface that is being analyzed. The ID lists the chains that make up the interface - such as L_A or LH_A. Used to query other tables |
| chains_side1 | Interface Side 1 ID – chains that make up interface side 1 such as L or LH. Used to query other tables |
| chains_side2 | Same as above, but for side 2 |
| nchains_side1 | Number of chains that make up side 1 |
| nchains_side2 | Number of chains that make up side 2 |
| dSASA | Difference in Buried Solvent Accessible Surface Area as described above<br>(*SASA separated - SASA together*) |
| dSASA_hphobic | Hydrophobic component of dSASA |
| dSASA_polar | Polar component of dSASA |
| dG | Difference in Rosetta energy as described above<br>(*E together - E separated*) |
| dG_cross | Total cross interface energy of pair-wise terms. Close to dG. |
| dG_dev_dSASAx100 | dG normalized by dSASA. Roughly an energy density calculation. |
| dG_cross_dev_dSASAx100 | dG_cross normalized by dSASA. Roughly an energy density calculation. |
| hbond_E_fraction | Fraction of dG attributable to cross interface hydrogen bonds. |
| sc_value | Shape Complementarity value |
| packstat | Packing quality measure |
| nres_int | Total interface residues |
| nres_all | Total structure residues |
| complex_normalized | Total energy/Total residues |
| ***Table interface_sides*** |  |
| struct_id | Structure number – used to query other tables for same loaded structure |
| interface | Interface ID – The interface that is being analyzed. The ID lists the chains that make up the interface - such as L_A or LH_A. Used to query other tables |
| side | The Side that this data pertains to. side1 or side2. |
| chains_side1 | Interface Side 1 ID – chains that make up interface side 1 such as L or LH. Used to query other tables |
| chains_side2 | Same as above, but for side 2 |
| interface_nres | Total number of residues for interface side |
| dSASA | dSASA of interface side |
| dSASA_sc | dSASA of the side chains of the interface side |
| dhSASA | Hydrophobic dSASA of the interface side |
| dhSASA_sc | Hydrophobic dSASA of the side chains of the interface side |
| dG | dG of the interface side |
| energy_int | Energy of the interface residues complexed |
| energy_sep | Energy of the interface residues separated |
| avg_per_residue_energy_dG | Average per-residue dG of the side |
| avg_per_residue_energy_int | Average per-residue energy of the side while complexed |
| avg_per_residue_energy_sep | Average per-residue energy of the side while separated |
| avg_per_residue_dSASA | Average per-residue dSASAS of the side |
| avg_per_residue_SASA_int | Average per-residue total SASA of the side while complexed |
| avg_per_residue_SASA_sep | Average per-residue total SASA of the side while separated |
| aromatic_fraction | Fraction of total residues that are aromatic |
| aromatic_dSASA_fraction | Fraction of aromatic dSASA from total dSASA of the side |
| aromatic_dG_fraction | Fraction of aromatic dG from total dG of the side |
| interface_to_surface_fraction | Fraction of side interface residues to total surface residues |
| ss_sheet_fraction | Fraction of side interface residues that are identified as beta-sheet from DSSP |
| ss_helix_fraction | Fraction of side interface residues that are identified as helix from DSSP |
| ss_loop_fraction | Fraction of side interface residues that are identified as loop from DSSP |
| **Table interface_residues** |  |
| struct_id | Structure number – used to query other tables for same loaded structure |
| interface | Interface ID – The interface that is being analyzed. The ID lists the chains that make up the interface - such as L_A or LH_A. Used to query other tables |
| resNum | Residue number of particular residue for which this data reports. |
| chains_side1 | Interface Side 1 ID – chains that make up interface side 1 such as L or LH. Used to query other tables |
| chains_side2 | Same as above, but for side 2 |
| side | The Side that this residue is part of. Side1 or side2. |
| dSASA | dSASA of the residue |
| dSASA_sc | dSASA of the side-chain of the residue |
| dhSASA | Hydrophobic dSASA of the residue |
| dhSASA_sc | Hydrophobic dSASA of the side-chain of the residue |
| SASA_int | Total SASA of the residue while complexed |
| SASA_sep | Total SASA of the residue while separated |
| relative_dSASA_fraction | Fraction of dSASA of the residue relative to the total SASA separated. 0 indicates no burial, while 1 indicates total burial. |
| dG | dG of the residue |
| energy_int | energy of the residue while complexed |
| energy_sep | energy of the residue while separated |

**Table F.** Additional *AntibodyFeature* reporter tables

| Column | Description |
| --- | --- |
| **Table ab_metrics** |  |
| struct_id | Structure number – used to query other tables for same loaded structure |
| numbering_scheme | The numbering scheme used to define the antibody |
| cdr_definition | The CDR Definition used to define the start and end points of the CDR |
| cdr_residues | The total number of CDR residues |
| antigen_present | Is an antigen present? |
| antigen_chains | If so, what are its chains? |
| net_charge | Net charge of the antibody |
| paratope_charge | Net charge of the paratope |
| paratope_SASA | Total SASA of the paratope |
| paratope_hSASA | Total Hydrophobic SASA of the paratope |
| paratope_pSASA | Total Polar SASA of the paratope |
| VL_VH_packing_angle | VH/VL Packing Angle |
| VL_VH_distance | VH/VL Distance |
| VL_VH_opening_angle | VH/VL Opening Angle |
| VL_VH_opposite_opening_angle | VH/VL Opposite Opening Angle |
| is_camelid | Is the antibody a Camelid antibody? |
| **Table cdr_metrics** |  |
| struct_id | Structure number – used to query other tables for same loaded structure |
| CDR | The identity of the CDR |
| length | The length of the CDR |
| start | Starting resNum of the CDR |
| end | Ending resNum of the CDR |
| ag_ab_contacts_total | Total unique Antigen/Antibody contacts from this CDR. An atomic contact is defined as at least 5 atoms of the Ag that are within 5 Å of an Ab atom |
| ag_ab_contacts_nres | Total number of residues making contact with Antigen |
| ag_ab_dSASA | dSASA of CDR/Antigen interface |
| ag_ab_dSASA_sc | dSASA of only side chains of CDR/Antigen interface |
| ag_ab_dhSASA | Hydrophobic dSASA of CDR/Antigen interface |
| ag_ab_dhSASA_sc | Hydrophobic dSASA of only side chains of CDR/Antigen interface |
| ag_ab_dG | dG of CDR/Antigen interface |
| SASA | Total SASA of the CDR |
| charge | Total charge of the CDR |
| energy | Total energy of the CDR |
| anchor_CN_distance | Distance from C of first residue to N of last residue of the CDR |
| aromatic_nres | Total aromatic residues of this CDR |
| **Table cdr_residues** |  |
| struct_id | Structure number – used to query other tables for same loaded structure |
| resNum | Residue number of particular residue for which this data reports. |
| CDR | The CDR for which this residue belongs |
| position | The position (1 – length) for which this is in the CDR |
| ag_contacts | The total number of Antigen contacts this residue makes as described above. |
| **Table ab_h3_kink_metrics** |  |
| struct_id | Structure number – used to query other tables for same loaded structure |
| kink_type | The identified Kink type – either Kinked or Extended. |
| begin | Start of the kink |
| end | End of the kink |
| anion_res | Identity of the anion residue of the Kink |
| cation_res | Identify of the cation residue of the Kink |
| RD_Hbond_dis | Distance of the sc-sc Hbond across the strands at the beginning of the H3 kink (typically Asp-Arg) |
| bb_Hbond_dis | Returns of the bb-bb Hbond across the strands at the beginning of the kink (typically Asp-Arg) |
| Trp_Hbond_dis | Distance of the Trp sc-bb Hbond across the H3 kink residues (n-1 to n+2) |
| qdis | Distance from the four kink residues of the H3 C-terminal end that make up the qdih below. |
| qdih | Dihedral angle from the four kink residues of the H3 C-terminal end |

